# Tissue-to-Analysis Framework Enables Multiscale Mapping of the Architectural and Cellular Organization in the Human Dorsal Root Ganglion

**DOI:** 10.64898/2026.01.28.702323

**Authors:** Matthew Adam Hunt, Sven David Arvidsson, Gustaf Wängberg, Zhening Zhang, Zerina Kurtović, Emerson Krock, Lisbet Haglund, Camilla I Svensson

## Abstract

Transcriptomic studies have helped us understand the dorsal root ganglia’s cellular milieu, yet our knowledge of protein expression and spatial organization/architecture remains less defined. Here we establish a comprehensive resource from processing through analysis of hDRG tissue. We optimize tissue-handling strategies and evaluate 114 antibodies targeting neuronal and non-neuronal cell types, identifying protocols that preserve neuronal morphology and antigen retain specificity. Integrating these workflows with our Deep Learning-assisted image analysis pipelines, we quantify size, expression, and spatial organization across 35,721 neurons from 15 donors. Female donors exhibited significantly larger neuronal somata, indicating sexual dimorphism. Neuronal subpopulations display clear spatial clustering. We further characterized the perineuronal niche, marked by dense vascularization, nuclear remodeling in perineuronal cells, and age-related increased turnover of neuron-associated macrophages. Together, this resource provides standardized methodologies and quantitative frameworks for reproducible protein-level interrogation of human sensory biology and pain mechanisms.

## Introduction

The dorsal root ganglia (DRG) house the cell bodies of the primary sensory neurons responsible for signaling pain, touch, temperature, and itch from the periphery to the central nervous system. Dysfunction of the DRG is considered a major contributor to chronic pain and sensory disorders. Despite its critical role, direct investigation of human DRG (hDRG) biology has historically been limited by access to tissue and technical challenges. Foundational insights into somatosensory neuron diversity emerged from electrophysiological and anatomical studies that characterized A-beta, A-delta and C-fibers^1–3^ and later studies that identified their relationship with neuronal soma size^4,5^. Advances in single-cell (sc) and single-nucleus (sn) RNA sequencing (RNAseq) have refined this framework, identifying between 13-22 transcriptionally distinct sensory neuron subtypes in rodent and human DRGs^6–12^. These transcriptomic atlases have transformed our understanding of sensory neuron diversity and provided an essential molecular foundation/resource for the field.

However, transcriptomic identity alone is insufficient to fully capture the functional and organizational complexity of the hDRG. While neuronal subtypes in rodent models display discrete and mutually exclusive expression of canonical markers^13^, those in the hDRG exhibit a broader expression profile^8,14–16^. Moreover, relationships between molecular identity, soma size, and spatial organization are less clearly defined in hDRGs^6,7,10,17^, making direct translation of animal models to the human system challenging.

In parallel, nociception is shaped not only by neurons, but also the environmental niche of the DRG and complex interactions with surrounding cell types. The DRG is highly vascularized^18,19^ and contains a diverse population of satellite glial cells (SGC), immune cells, endothelial cells and stromal cells^9,18,20–23^ that can act on neuronal somata. Rodent studies have shown that SGCs^24–26^ and macrophages^27–31^ can modulate neuronal excitability and pain-like behaviors. And recent studies using hDRGs have further revealed distinct species-specific differences in immune organization, including the presence of CX3CR1-expressing perineuronal macrophages that are largely absent in rodents^18,21^. Despite their abundance and strategic localization, systematic protein-level and spatial characterization of non-neuronal cell populations in hDRG remains limited.

A major barrier to progress is the lack of standardized methodologies for protein-level and spatial analysis of hDRG tissue. Most hDRG samples are obtained post-mortem from organ donors, often without clinical pre-selection, and exhibit substantial biological and technical variability. Tissue quality and antigen preservation are highly sensitive to collection, fixation and processing protocols^32–35^. Donor-to-donor variability has hindered the development of high-throughput technologies for analyzing neuronal populations and the cytoarchitecture of the hDRG. Given these obstacles, cell-to-cell interactions risk being overlooked, marker based functional interpretations become muddied and reproducibility of antibody-based studies remain limited.

Here, we address this gap by establishing a comprehensive resource for immuno-staining and imaging-based spatial analysis of the hDRG. We systematically evaluate how commonly used tissue processing and fixation strategies affect tissue integrity and immuno-staining interpretability. We perform large-scale screening and validation of antibodies targeting neuronal and non-neuronal cell types, providing practical guidance on marker reliability and protocol sensitivity. To enable quantitative and scalable analysis, we developed deep-learning image-analysis workflows that integrate automated segmentation and morphological/spatial analyses that are robust to donor and technical variability. Together, this work provides a robust and accessible foundation for reproducible studies of hDRG biology and enables deeper integration of molecular atlases with spatial and functional investigations relevant to pain and sensory disorders.

## Results

### Tissue quality and immunohistochemistry optimization of the hDRG

The lack of standardized tissue-processing guidelines and antibodies validated on post-mortem hDRG tissue represents a major obstacle for reproducible immunohistochemistry (IHC). To aid the community, we first compiled a list of published antibodies targeting 72 proteins said to work on hDRG tissue (Supplementary Table 1). Leveraging cell-type specific expression patterns from single-nucleus and single-cell RNAseq datasets expanded the list to 114 candidate antibodies (Supplementary Table 2-4).

### Effects of tissue processing on neuronal morphology

We evaluated the impact of tissue processing methods on hDRG morphology and antigen preservation. Post-mortem hDRGs were either immediately fixed in 4% PFA (iFix; donors 1-11), immediately frozen and subsequently fixed in 4% PFA (iFr-F; donors 1-6, and 12-16) or cryosectioned directly from immediately frozen and fixed on the slide (iFr-Fos; donors 12-17) (Fig. 1a, Supplementary Table 5). Both iFix and iFr-F protocols were cryoprotected in 30% sucrose prior to embedding, while iFr-Fos were not. Across samples, neuronal soma frequently appeared distorted relative to surrounding tissue, consistent with differential contraction or expansion during processing, resulting in apparent “shrunken” neuronal somata surrounded by tissue voids. To objectively evaluate preservation of neuronal morphology, we defined the Tissue Quality Index (TQI) as the fraction of observed neuronal soma area relative to the expected, where expected was the soma plus adjacent voids. Neuronal somata were identified via NF200 staining together with DAPI to visualize surrounding nuclei and tissue architecture (Fig. 1b-c). iFr-Fos samples showed a markedly reduced TQI (0.73±0.13), indicating substantial loss of tissue integrity. In contrast, both iFix and iFr-F protocols preserved neuronal morphology more effectively, with iFix samples exhibiting the highest TQI (0.97±0.02) and significantly higher precision than iFr-F TQI (0.93±0.04, Fig. 1b-c). These data indicate that cryoprotection is essential for preserving hDRG neuronal architecture, and that immediate fixation provides the most consistent tissue integrity.

**Fig. 1:**
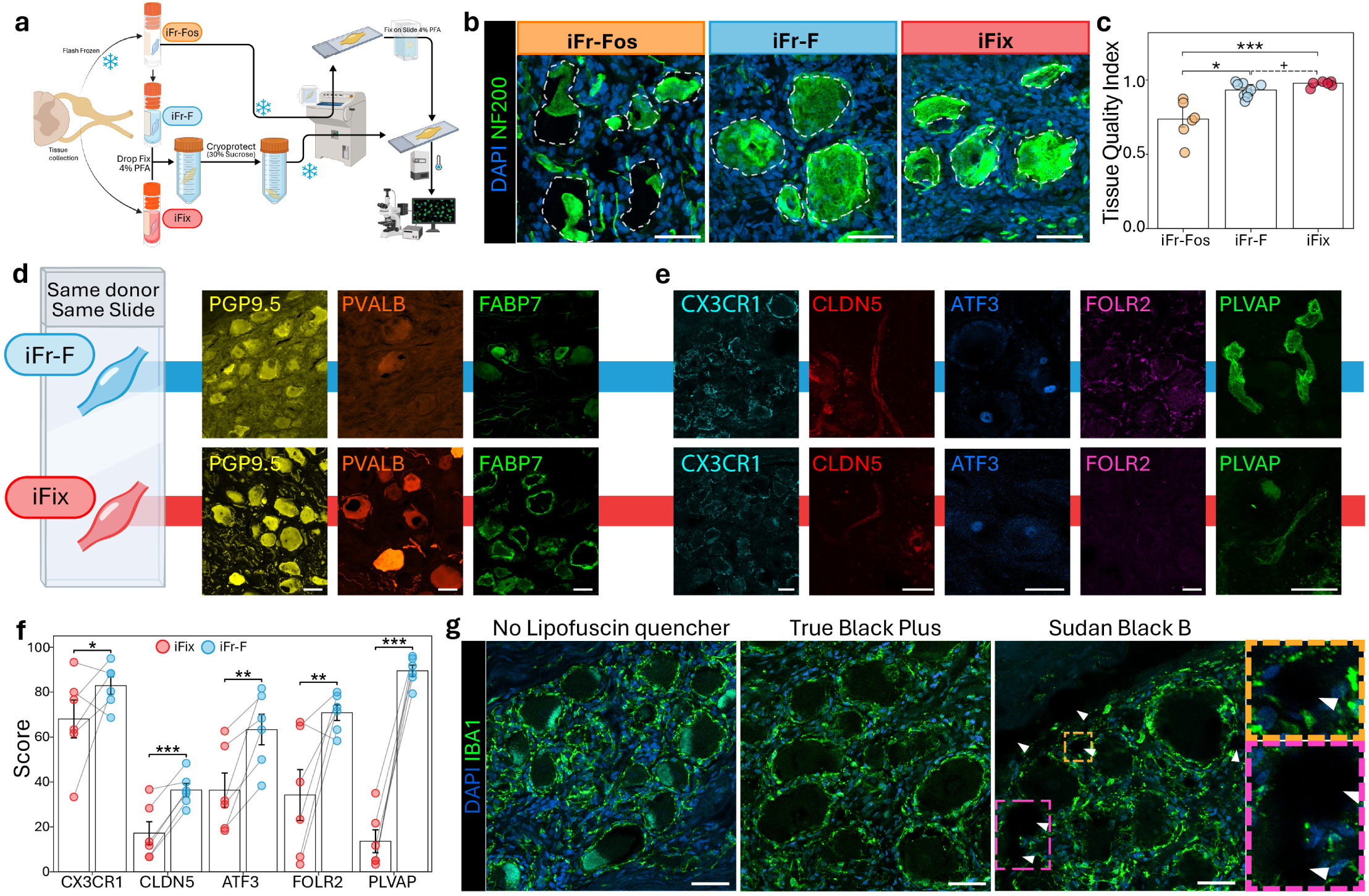
Optimization of hDRG staining. **a**, Schematic representation of preprocessing protocols used. **b**, Representative images of pan-neuronal staining NF200 (green) and DAPI (Blue) showcasing the effect of different preprocessing protocols or neuronal soma. Dotted line simulates the expected coverage of neuronal tissue. **c**, Quantification of neuronal preservation in iFr-Fos (n=6), iFr-F (n=9), and iFix (n=6) samples. \**P* < 0.05, \*\*\**P* < 0.001. + indicates significant (*P* < 0.05) difference in precision by comparing variance in Levene’s test **d**, Representative images from paired samples of PGP9.5 (yellow), PVALB (orange), and FABP7 (green) showing a loss of specificity in iFr-F samples while maintaining a clean signal on iFix samples. **e**, Representative images of antibodies with diminished signal in iFix samples while keeping specific signal in iFr-F samples. **f**, Average from 3 different investigators scoring staining quality (0-100) across paired samples from 6 different donors for PLVAP, CLDN5, ATF3, FOLR2, and CX3CR1. Paired T-test with False Discovery Rate (FDR) calculated with two-stage step-up method of Benjamini, Krieger and Yekutieli. \**q* <0.05, \*\**q* < 0.01, \*\*\**q* < 0.001. **g**, Representative images of same donor stainings for DAPI (Blue) and IBA1 (green) showing regular amount of lipofuscin (No Lipofuscin quencher) and the loss of lipofuscin signal following True Black Plus post-treatment or Sudan Black B 0.02% pre-treatment. Zoom-in regions in the Sudan Black B staining highlight the unspecific accumulation of Sudan Black B. Scale bar = 50 μm.

### Fixation duration differentially affects epitope preservation

While immediate fixation is commonly recommended to preserve ultrastructure and minimize protein diffusion^33,36^, prolonged fixation can compromise epitope accessibility and limit downstream applications^32,35,37,38^. We assessed the impact of fixation duration on sensitive epitopes (FASN, CD31, PLVAP, CX3CR1) by staining quartered iF-Fr hDRGs from a single donor fixed for 2 or 8 hours (donor 14). CX3CR1 and FASN staining were largely unaffected by fixation duration, whereas CD31 signal intensity was reduced by longer fixation even though signal-to-noise ratio remained similar. In contrast, PLVAP staining showed a clear reduction in signal-to-noise ratio after prolonged fixation (Extended Data Fig. 1a-b). Based on these observations, we adopted a fixation of 3 hours for quartered iFr-F samples and 8–12 hours for iFix to account for larger tissue volumes and ensure adequate penetration. This window also accommodated variation in post-mortem collection timing, which contributed to potential inter-sample variability.

### Fixation-dependent antibody performance

To directly compare IHC performance between fixation strategies, paired iFix and iFr-F hDRGs from the same donors were processed on the same slides and stained in parallel (Fig. 1d-f). Among the 114 antibodies screened, 12 exhibited fixation-dependent differences (Supplementary Table 2). While pan-macrophage marker IBA1 showed minor differences in perinuclear staining, cell specificity was preserved across protocols (Extended Data Fig. 1c). PGP 9.5, PVALB, and FABP7 antibodies all exhibited markedly lower signal-to-noise ratios in iFr-F tissue (Fig. 1d), to the point where FABP7 completely lost any recognizable SGC specificity, which was confirmed using a second validated FABP7 antibody (Extended Data Fig. 1d). Antibodies against nuclear and membrane-associated markers (ATF3, PLVAP, CD31, FOLR2, CX3CR1, PIRT, and KCNA2) showed consistently superior performance in iFr-F samples, whereas staining in iFix samples remained cell specific but exhibited lower signal-to-noise ratios (Fig. 1e and Extended Data Fig. 1e). Of note, fixation-sensitive CD31 staining was restricted to a single antibody clone (AF806).

Five fixation-sensitive markers (ATF3, PLVAP, CLDN5, FOLR2, and CX3CR1) were further evaluated by three blinded investigators using a 0-100 scoring scale in paired samples from six donors. Following our initial observations, all antibodies consistently scored higher in iFr-F samples (Fig. 1f).

### Limited utility of signal amplification for low-affinity targets

We next assessed whether tyramide signal amplification (TSA) could rescue staining for antibodies with low signal to noise ratio independent of fixation, including TRPM8 (NBP1-97311SS) and CHRNA3 (15183), or high variability including PGP9.5 (RBK064-05) and FASN (ab128870). TSA substantially increased signal to noise ratio and staining quality for FASN but did not improve specificity for PGP9.5 or enhance staining quality for low-signal antibodies (Supplementary Fig. 1). These results indicate that, for the antibodies tested here, TSA cannot compensate for freezing-induced epitope loss or intrinsically weak antibody performance.

### Quenching neuronal autofluorescence

Lipofuscin, an age-associated aggregate of lipids and protein, frequently exhibits a punctate autofluorescent pattern within neuronal somata^39^. Lipofuscin’s broad emission spectrum can obscure or confound detection of neuronal and perineuronal markers^40,41^. Reducing lipofuscin autofluorescence is therefore advantageous. We compared an ethanol-based Sudan Black B (SBB) pretreatment with a water-soluble TrueBlack Plus post-treatment. Both approaches effectively attenuated lipofuscin autofluorescence while preserving the signal to noise ratios (Fig. 1g). However, SBB frequently formed insoluble aggregates that persisted after washing and produced artifactual voids (Fig. 1g), additionally its ethanol-based formulation was incompatible with TSA protocols. TrueBlack Plus provided superior performance and was therefore used for all subsequent analyses when required.

### Neuronal protein expression in the hDRG extend beyond transcriptomic predictions

Using the optimized IHC workflow described above, we examined how neuronal protein expression compares to published scRNAseq expression (Fig 2). Neuronal subpopulations, clustered via integration of publicly available hDRG RNAseq datasets^8,10^, were classified via shared gene expression profiles across datasets, and annotated to match the hDRG atlas^9^. Among 47 antibodies predicted to label neurons, 29 (24 markers) produced neuron-specific staining patterns (Supplementary Table 2). Protein signal for 20 neuronal markers was quantified using one full section each. All neurons were manually classified as negative, detectable, or bright and compared with RNA expression.

**Fig. 2:**
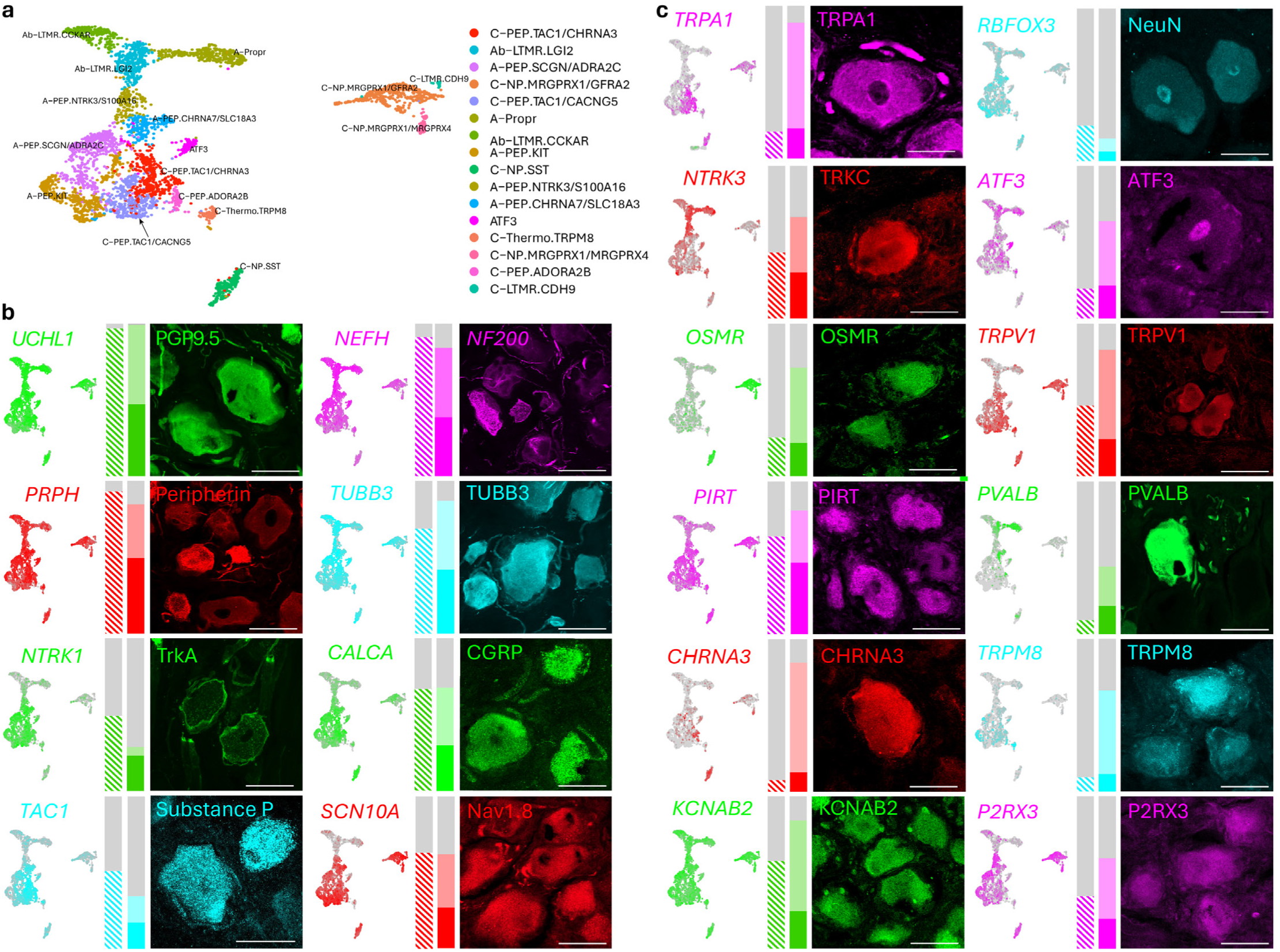
Protein and transcriptomics correlation of neuronal markers. **a**, UMAP showing identified neuronal subpopulations in an integrated dataset from 4 studies matched to previous single soma populations and newly identified populations from the hDRG transcriptomics atlas. **b**-**c**, Comparison of RNA and protein. Feature plots representing RNA transcript expression in the previously define neuronal populations followed by the percentage of neurons with log2Fc>0.5 for each associated marker to the left. To the right, a column showing single slide quantifications (210-966 neurons) showing neurons with negative staining (gray), detectable protein expression (dim colored portion), or bright protein expression (strong colored portion) followed with a representative 40x image highlighting high expressors. Columns in B show genes and proteins with a high similarity between RNA percentage and detectable protein. Column C shows genes and proteins with high similarity between RNA percentage and bright expressors.

Proteins reported as broadly expressed across sensory neurons, PGP9.5 and NF200, had proportions of positive neurons consistent with detectable RNA (Fig. 2b). In contrast, β3-tubulin (TUBB3) displayed a higher proportion of detectable neurons than was reported via RNA expression, nonetheless neurons expressing these pan-neuronal marker associated genes were present in all RNAseq subpopulations (Fig. 2b and Extended Data Fig. 2). Genes encoding proteins highly enriched in human nociceptors^16,42–45^, such as TrkA, Substance P, CGRP and Nav1.8, exhibited RNA expression profiles more similar to detectable protein than high expressors. Most additional evaluated subset markers had a broader neuronal protein distribution than suggested by RNAseq data, where transcriptomic prevalence matched the percentage of neurons exhibiting high expression rather than detectable levels (Fig. 2c). These results indicate broad protein expression in human neurons, and that RNA-defined populations correspond preferentially to high-level protein expressors.

### hDRG organized into laminar neuron rich islands

During qualitative inspection of hDRGs we observed a striking laminar structural organization of alternating neuron-rich and fiber-rich regions, which extended radially inward from the capsule. This organization was particularly evident following optical clearing of a quartered hDRG and light-sheet imaging, which revealed repeated neuronal strata separated by thick axonal bundles (Fig. 3a-b). While some studies have described neuronal/fiber density^46^ in the hDRG to our knowledge, this, laminar, radial stratification has not been previously described in hDRG.

**Fig. 3:**
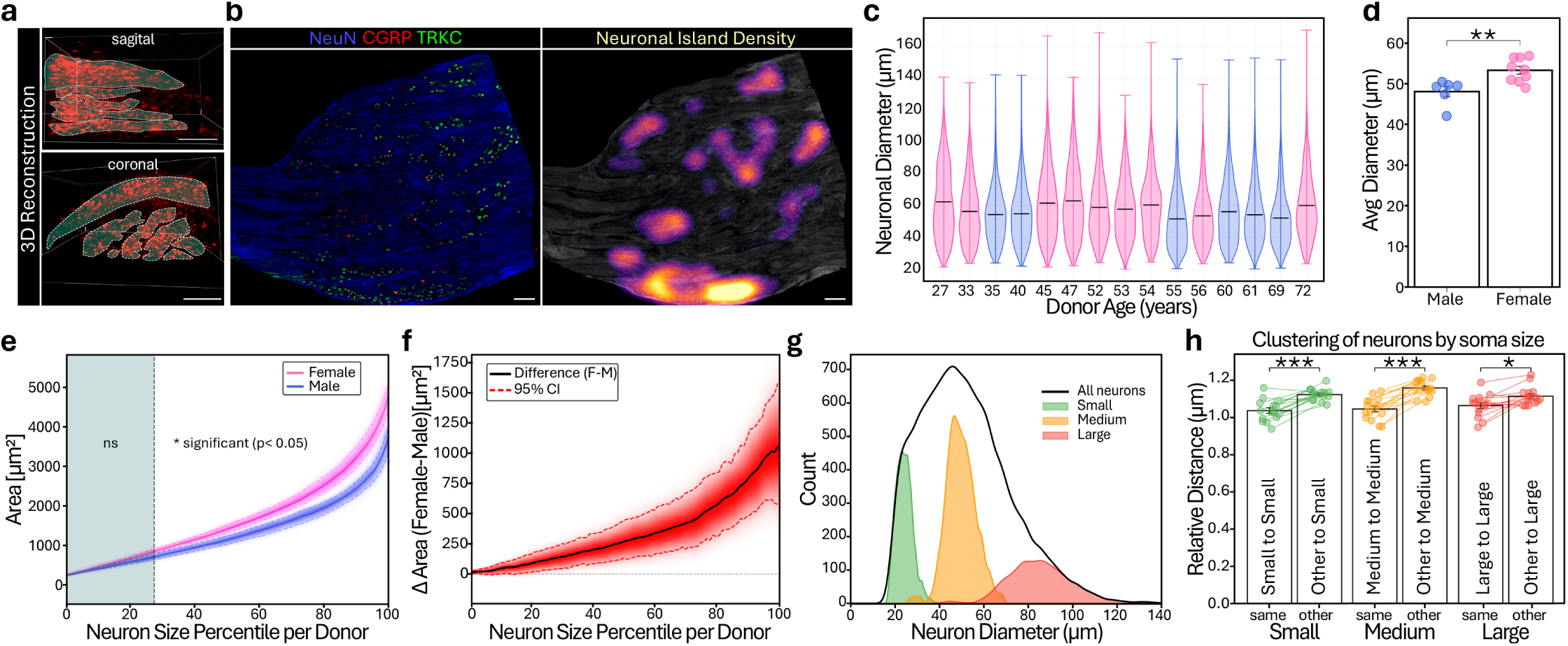
hDRG neuronal characterization. **a**, Neuron-rich regions separated by thick fiber regions from cleared hDRG with neuronal cell bodies marked by lipofuscin autofluorescence (red), where neuronal islands are highlighted by dotted lines. **b**, Immunostaining of NeuN (blue), CGRP (red), and TrkC (green) in an hDRG section showing neuronal clusters, followed by a heatmap of neuronal density highlighting neuron-islands. Scale bar = 500 µm. **c**, Violin plot of neuronal size (35,721 neurons) across all donors (n = 15), ordered by age and colored by sex (female: pink; male: blue). **d**, Bar plot of donors’ means neuronal size (female: pink; male: blue). ** indicates p < 0.01, Mann–Whitney U test. **e**, Average neuron size (cross-sectional area) for in-donor percentile averages for females (magenta) and males (blue). The area to the right of the black dashed line shows a significant difference (*P* < 0.05) between percentile averages (Benjamini–Hochberg FDR-corrected Mann–Whitney U test). Density clouds represent bootstrap distribution probability (white = low; saturated color = high), with colored dashed lines representing the bootstrap-estimated 95% confidence interval. **f**, The female–male difference in average neuron size (cross-sectional area) for in-donor percentile averages (black). Density clouds represent bootstrap distribution probability (white = low; saturated red = high), with red dashed lines representing the bootstrap-estimated 95% confidence interval. **g**, Size distribution of small (green), medium (orange), and large (red) neurons by Feret diameter. Neurons are classified per donor as: small (≤12.5th percentile), medium (37.5th–62.5th percentile), and large (≥87.5th percentile), with 25% padding between classes to ensure distinct populations. **h**, Spatial clustering of neurons by size class. For each size class, two distances are compared: (1) from each neuron of that class to its three nearest same-class neighbors, and (2) from each neuron outside that class to its three nearest neighbors within that class. Distances were averaged per donor and normalized to the median of the between-class distances. Bars show donor means, paired by donor (Wilcoxon signed-rank test: *P < 0.05, **P < 0.01, ***P < 0.001).

### hDRG neurons exhibit sex dependent differences in size

We developed ensemble deep learning models trained to identify neuronal soma (Methods and Extended Data Fig. 3a-d). Using this, we analyzed the morphology and spatial organization of 35,721 neurons in 4 multi-marker antibody panels across 15 donors (Fig. 3c). Showcasing this high density in neuron-rich islands, the average nearest-neighbor distance was 33.7 μm (Extended Data Fig. 4a), less than the average neuronal diameter of 51.2 μm inferred from maximum Ferret diameter, a measure invariant to shrinkage artifacts. Neuronal soma size varied significantly between donors. Surprisingly, sex explained 50.4% of the between-donor variance, with females having significantly larger neurons. (Fig. 3d). This sex difference scaled exponentially with neuron size percentiles, with no significant difference among small neurons but significant differences in both mid-sized and large neurons (Fig. 3e-f). This remained significant even when accounting for donor age, weight and height, as well as fixation protocol, indicating that sex dependent differences were not driven by confounders. While examining the size of neurons, we noticed that similar sized neurons seemingly clustered together. Quantitative analysis confirmed significant clustering of donor-normalized small, medium, and large neurons across all donors (Fig. 3g-h).

Consistent with our initial analyses, pan-neuronal markers TUBB3, NF200 and peripherin were detectable in most neurons, while NeuN labeled a small subset (Extended Data Fig. 4b). Additionally, we observed that neuronal subset defining proteins CGRP, TrkC, TRPA1, TRPV1, and CHRNA3 were detectable broadly across neurons (Fig. 4). For each marker/panel, the proportion of neurons is shown as detectable and high expressors (++). For additional quantifications, neurons were classified as high expressors (++) or low expressors (low) defined as not a high expressor. For each panel, combinations of high vs low expressors were displayed in Venn diagrams, bar plots displaying donor-normalized neuron diameter, and relative distance plots.

**Fig. 4:**
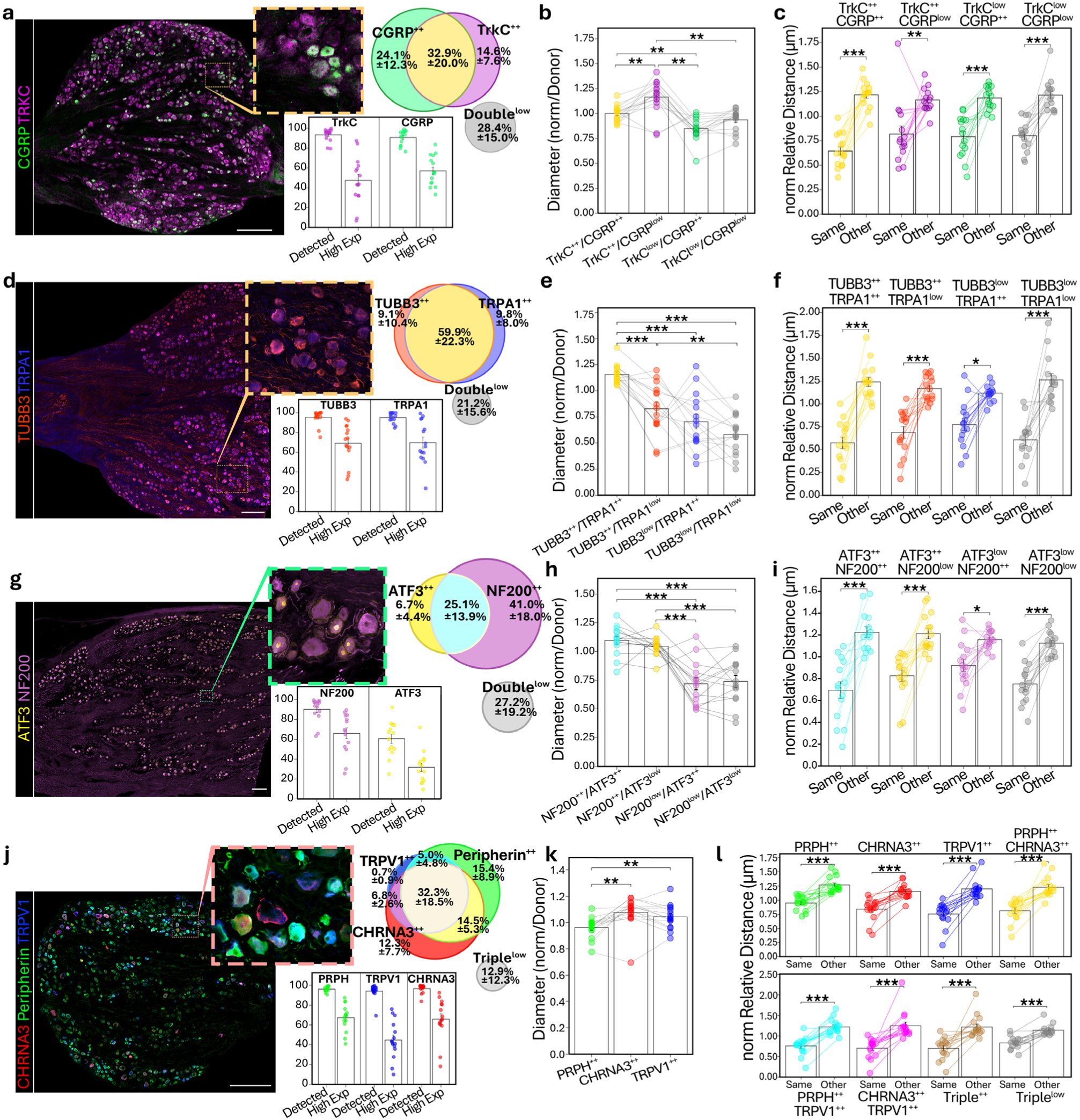
hDRG neuronal characterization. **a-c**, Panel 1, CGRP (Green), TrkC (Magenta) 15 donors, 9,223 neurons. **d-f**, Panel 2, TUBB3 (Red), TRPA1 (Blue) 14 donors, 10,408 neurons. **g-i**, Panel 3 ATF3 (yellow), Nf200 (Magenta), 15 donors, 8,298 neurons. **j-l**, Panel 4, Peripherin (Green), TRPV1 (blue), CHRNA3 (red) 15 donors, 7,972 neurons. **a, d, g, j**, Representative whole-section Immunostaining of panel specific markers, Scale bar = 500 um. Bar plot of donors means prevalence of detectable and high expressor. Venn diagram of high expressor (++) with sd. **b, e, h, k** Normalized somata size by donor across combinations in the different panels. (Statistics: \**P* < 0.05, \*\**P* < 0.01, \*\*\**P* < 0.001, Wilcoxon signed-rank, Benjamini-Hochberg FDR correction). **c, f, i, l,** Spatial clustering of neurons by marker-defined subpopulation. For each subpopulation, two distances are compared: (1) from each neuron of that subpopulation to its three nearest same-subpopulation neighbors, and (2) from each neuron outside that subpopulation to its three nearest neighbors within that subpopulation. Distances were averaged per donor and normalized to the median of the between-subpopulation distances. Bars show donor means, paired by donor (Wilcoxon signed-rank test: *P < 0.05, **P < 0.01, ***P < 0.001).

### Relationship between neuronal subtypes and soma size

TrkC and CGRP (Fig. 4a-c), enriched in A-fibers and peptidergic nociceptors, revealed TrkC⁺⁺ neurons were larger than CGRP⁺⁺ neurons and TrkC^++^ CGRP^Low^ neurons were significantly larger than CGRP⁺⁺ neurons. TrkC^++^ CGRP^++^, putative A-fiber nociceptors, had a soma size close to the population mean for each donor.

TUBB3 and TRPA1 were co-expressed in ∼70% of high-expressing neurons (Fig. 4d-f). TUBB3^low^ and TRPA1^low^ neurons were smaller, consistent with reduced *TUBB3* expression reported in C-LTMR neurons^47^. In our integrated sequencing data, TUBB3 displayed low expression in C-LTMR as well as other small-soma populations such as C-NP populations (Extended Data Fig. 2). However, in contrast to sequencing data, TRPA1 very high expressors (+++) were significantly larger, highlighting the importance of protein expression characterization in neuronal subsets (Extended Data Fig. 4c).

While ATF3^++^ neurons were more prevalent than previously reported^8–10,48^, NF200^++^ neurons were significantly larger than NF200^low^neurons, as previously reported^10,44,49^, which was independent of ATF3 expression (Fig. 4g-i).

Nociceptor-enriched markers showed co-expression in peripherin and TRPV1, and between TRPV1 and CHRNA3, the nicotinic acetylcholine receptor subunit associated with “silent” nociceptors ^50^ (Fig. 4j). Triple high expressors PRPH^++^TRPV1^++^CHRNA3^++^ accounted for 32.3% of quantified neurons. Donor-normalized analyses revealed systematic soma size biases. Peripherin⁺⁺ neurons were significantly smaller than both TRPV1⁺⁺ and CHRNA3⁺⁺ neurons. Across single-marker comparisons, peripherin⁺⁺ neurons were smaller than peripherin^low^ neurons, whereas CHRNA3⁺⁺ neurons were substantially larger than CHRNA3^low^ neurons (Fig. 4j-l). Further classification of peripherin into very high expressors strengthened the relationship between peripherin^+++^ and smaller soma (Extended Data Fig. 4d). While the association of peripherin and TRPV1 with smaller neurons is consistent with prior human and rodent studies^14,44,49,51–54^, the enrichment of CHRNA3 protein in larger neurons contrasts with previous publications^9,50^.

Noteworthy, differences in neuronal size populations were only apparent after donor normalization and there was not a clear cut for neuronal size for the neuronal populations analyzed (Extended Data Fig. 4e-i). Together, these data demonstrate that neuronal populations exhibit most of the generalized size associations typically observed in animal work, but only when normalizing size to each hDRG.In addition, individual neuronal subtypes also exhibited significant sex-dependent differences in size and abundance (Extended Data Fig. 5).

### Molecularly defined neuronal subpopulations exhibit robust spatial clustering

Further clustering evaluation of molecularly defined populations revealed a significant clustering in every subpopulation examined across the four marker panels. Whether defined by low, high, or very high expression levels; whether comprising abundant or sparse populations; and regardless of combinatorial complexity (Fig. 4l). This universal clustering, observed across all classifications tested, demonstrates that spatial organization is a fundamental architectural principle of the hDRG. This is to our knowledge, the first donor level proof of widespread, organized self-clustering of neuron subtypes within the neuron rich islands in the hDRG.

### Cell populations form a complex niche in the neuronal islands of the human DRG

The neuron-islands of the hDRG contain a diverse population of non-neuronal cells. While previous studies have shown the presence of glial, immune, vascular, and stromal cells, their spatial organization and proximity to neuronal somata have not been systematically mapped. To address this, we performed a marker-based analysis of major non-neuronal cell populations in the hDRG. Antibody specificity was validated based on expected staining patterns and concordance with published scRNAseq datasets (Fig. 5a, Extended Data Fig. 6). Out of the 68 antibodies tested, 46 displayed their expected specific patterns (Supplementary Table 3).

**Fig. 5:**
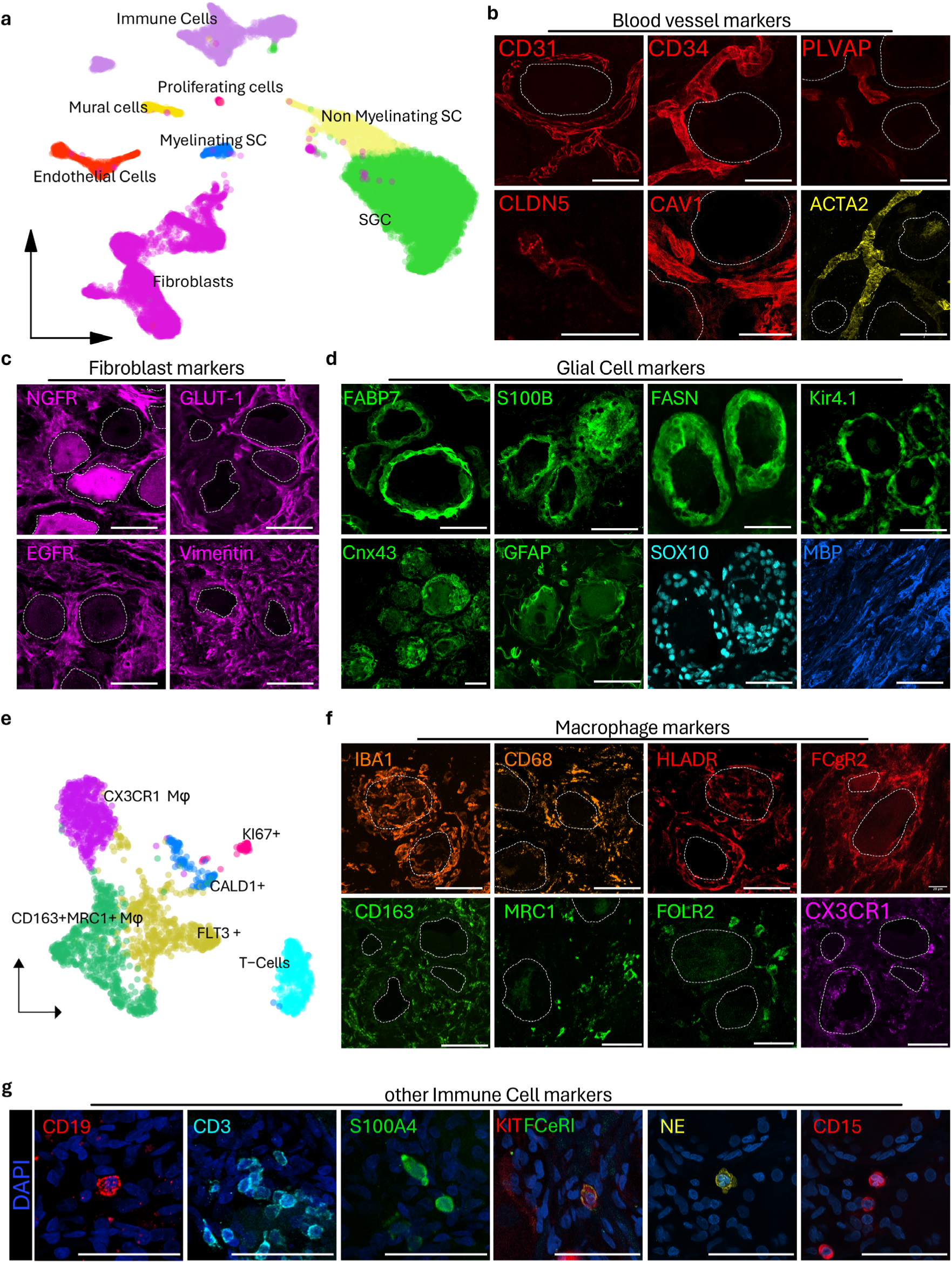
Non-neuronal markers in hDRG. **a**, UMAP showing main populations of non-neuronal cells found in the hDRG. **b**, Markers that stain endothelial and mural cells. **c**, Fibroblasts markers. **d**, Markers expressed SGCs and Schwann cells. **e**, UMAP showing main subpopulations of immune cells found in the hDRG. **f**, Pan macrophage markers (top row) and subset specific macrophage markers (bottom row). **g**, Additional immune cell markers with DAPI (blue). Scale = 50 μm. Dotted line represents neuronal soma coverage.

The hDRG is extensively vascularized as shown with the pan endothelial marker CD31(Fig. 5b). CLDN5^+^ arterioles were primarily observed in the inter-neuron-island regions that transitioned into permeable PLVAP^+^ capillaries and venules in the neuron-island regions^18,55^. PLVAP^+^ vessels were frequently positioned close to neuronal somata, coinciding with absence of CLDN5 expression, indicating spatial organization of vascular permeability. CD34 is present in both endothelial cells and fibroblasts but showed stronger endothelial staining in the hDRG. Mural vascular smooth muscle cells were identified around larger venous and arterial vessels with ACTA2.

Fibroblast markers showed variable specificity across cell types (Fig. 5c). GLUT-1 showed high fibroblasts specificity, while NGFR was found as well in a subpopulation of neurons. Vimentin also detected glial cells (Extended Data Fig. 7a), and EGFR was broadly expressed across fibroblast populations, but with occasional low-level neuronal signal.

We validated 11 antibodies, labelling the following 8 glial cell markers (Fig. 5d). SGCs: FASN, FABP7, Connexin 43, and Kir4.1; myelinating Schwann cells: myelin basic protein (MBP); Schwann cells and SGCs: S100beta and SOX10. Additionally, GFAP labeled a subset of SGCs, but also stained SOX10⁻ fibroblast-like cells (Extended Data Fig. 7a).

Macrophages were the most abundant immune cell type in the hDRG, identified by broad-macrophage markers, IBA1, CD68, FCGR2A, and HvLA-DR (Fig. 5f, Extended Data Fig. 6f). Subset specific markers showed CX3CR1^+^ macrophages intertwined with SGCs and closely associated with neuronal somata, while CD163^+^ macrophages distributed more broadly (Fig. 5f). CD163^High^ macrophages further subdivided into CD163^high^MRC1^-^ and CD163^high^MRC1^+^ populations with the latter further containing a subset of FOLR2^+^ macrophages. Finally, an additional population of hDRG macrophages expressing KI67 were observed, indicating recent division, with elevated CX3CR1 expression (Extended Data Fig. 6e-f).

CD3^+^ T-cells were the 2^nd^ largest immune cell population and were dispersed throughout the hDRG, with occasional perineuronal clusters (Fig. 5g). Neutrophils identified by neutrophil elastase (NE) and CD15 antibodies were primarily localized to vascular compartments but were abundant in the DRG parenchyma in some samples (Extended Data Fig. 7b). Additional, less abundant, immune cell populations were identified with markers: CD19 (B cells), S100A4 (enriched in hDRG monocytes), and FcER1/KIT (mast cells). Noteworthy, clusters of immune cells were occasionally observed near the ganglion capsule (Extended Data Fig. 7c). Together, these data provide a protein-level map of non-neuronal cell populations in the hDRG.

### Rich vascular architecture in neuron-rich regions

To characterize the cellular microenvironment of the hDRG, we first focused on the vasculature. The density of vascularization in hDRG was drastically increased in the neuron-island regions, to the point where it appeared as every neuron has its own associated blood vessel. In iFr-F samples we used CD31 to identify all blood vessels and PLVAP to identify fenestrated blood vessels (Fig. 6a). The volume of PLVAP^+^ vessels in the neuron-island rich regions was 7 times greater than PLVAP^-^ vessels (Fig. 6b). Identifying all neurons and quantifying each neuron’s distance to the nearest PLVAP+ or PLVAP-vessel revealed the average distance from any neuron to a PLVAP^+^ vessel was only 2.23 µm, which was significantly less than the average distance to PLVAP^-^ vessels or even the average neuron to nearest neuron distance (Fig. 6c).

**Fig. 6:**
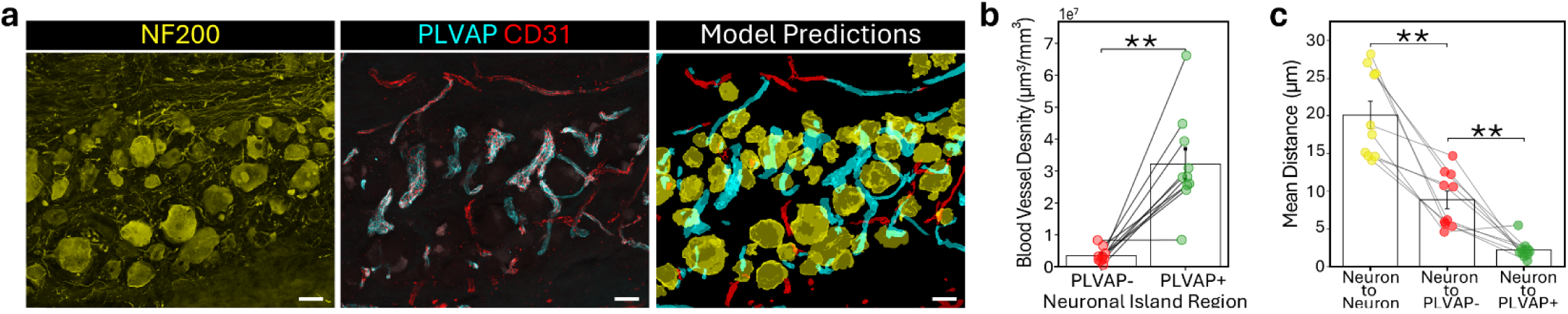
hDRG non-neuronal cells characterization. **a**, Representative images of neuron island stained with NF200 (yellow), followed by the colocalization of general blood vessel marker CD31 (red), and fenestrated vessels PLVAP (cyan) followed by the prediction output from the prediction model. **b**, Bar plot donor means showing blood predicted vessels volume in neuron-island regions. **c**, Bar plot from showing donor mean distance from neurons to each of other neurons, blood vessels, and fenestrated blood vessels respectively (Statistics: (iFr-F samples, n=10, \*\**p* < 0.01, Wilcoxon signed-rank, Benjamini-Hochberg FDR correction).**c**, Bar plot from showing donor mean distance from neurons to each of other neurons, blood vessels, and fenestrated blood vessels respectively (Statistics: (iFr-F samples, n=10, \*\**P* < 0.01, Wilcoxon signed-rank, Benjamini-Hochberg FDR correction).

### The onion layer algorithm, a novel method to classify perineuronal cells

When looking at hDRG neurons and the surrounding cells, it was readily apparent that the size and morphology of nuclei surrounding the neurons appeared concentrically stratified. Thus, we developed the “onion layer” algorithm, which automatically classifies objects in concentric zones radiating outward from neuronal surfaces (see Methods, Fig. 7a, Extended Data Fig. 8). While layer identification is independent of distance, they nonetheless correlate in overlapping distributions (Fig. 7b).

**Fig. 7:**
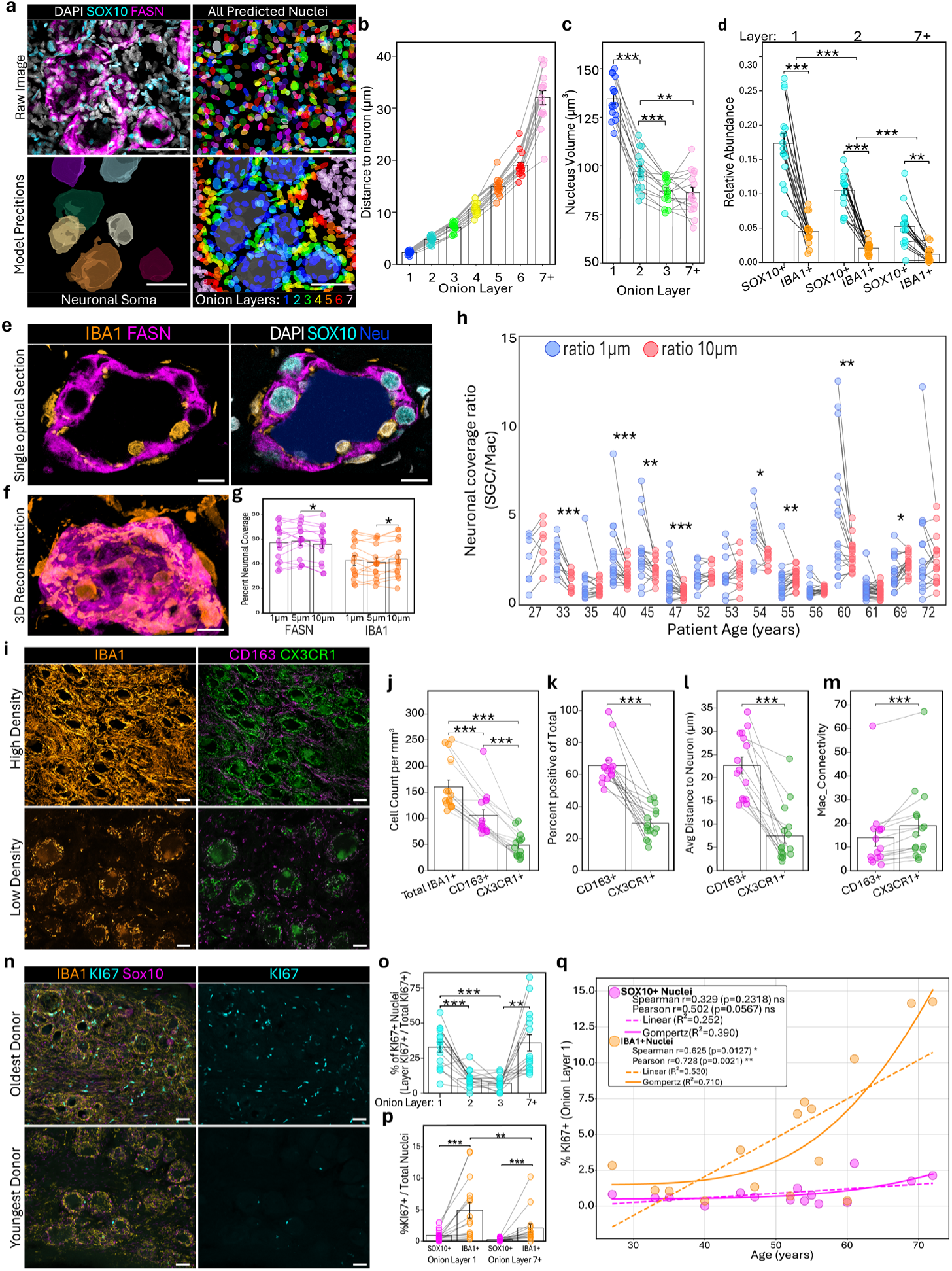
7hDRG non-neuronal cells characterization. **a,** Cellular layers around neurons in the hDRG. Glial marker staining in neuronal rich region (top left) and prediction models output for nuclei identification (top right), predicted neuronal location (bottom left), and onion layer classification (bottom right). **b**, Distance of layers. Bar plot showing the distance to closest neuron for cells on each category. **c**, Bar plot showing average nuclear volume for cells in layers 1,2,3, and 7+. **d**, Relative abundance of cell types with glial marker Sox10, and macrophage marker IBA1 on layers 1, 2, and 7+. **e-f**, Representative close-up image of neuronal surrounding region with SGC marker FASN, macrophage marker Iba1, glial marker SOX10, and model output for neuronal soma. **g**, Percentage of FASN and Iba1 coverage in the first 1, 5, and 10 mm. **h**, Ratio of SGC and macrophage coverage in the first 1, and 10 mm for the different donors analyzed. **i**, Macrophage analysis Representative image of general macrophage marker IBA1 (orange, left column) and marker for close neuronal contact macrophages, CX3CR1 (green), and other macrophages (CD163, magenta) in the hDRG for a sample with high (top) and low macrophage density (bottom). **j**, Macrophage density bar plot of number of macrophages per mm^3^. **k**, Macrophage CD163 and CX3CR1 relative class abundance within macrophages. **l**, CD163 and CX3CR1 distance to neuron bar plot. **m**, Quantification of macrophage connectivity. **n**, Representative images showing macrophage marker Iba1 (yellow), glial marker Sox10 (magenta), and dividing cell marker KI67 (cyan) for the oldest (top), and youngest (bottom) donor analyzed. **o**, Percentage of dividing cells on each nuclear layer. **p**, Percentage of Sox10 and Iba1 dividing cells for the first layer, and layer 7+. **q**, Scatter plot showing positive nuclei percentage in cell layer 1 for SOX10+ and Iba1+ nuclei and their respective linear and Gompertz R-squared adjusted values. N= 15 donors. *P* < 0.05, \*\**P* < 0.01, \*\*\**P* < 0.001.

### Expanded nuclear volume of supporting cells close to neuronal soma

Nuclear volume showed progressive expansion approaching neurons, with layer 1 nuclei measuring 134.5 µm³ compared to 86.2 µm³ in layer 7+ (Fig. 7c). This may reflect increased transcriptional activity in cells directly supporting neurons, as larger nuclei typically correlate with decondensed chromatin and higher gene expression^56–58^. To delve further into the perineuronal niche, we stained hDRGs for SOX10, FASN, IBA1 and DAPI. Interestingly, the size gradient was observed in the nuclei of both IBA1^+^ and SOX10^+^ cells and each contained a “large nuclei” subset in layer 1 (Extended Data Fig. 9a). IBA1^+^ nuclei, located closer to neurons (Extended Data Fig. 9b), were significantly smaller than same layer SOX10^+^ nuclei and more elongated (Extended Data Fig. 9c). SOX10^+^ and Iba1^+^ nuclei were significantly enriched in layers closest to the neuronal soma and displayed a remarkably consistent ratio of 4-fold greater SOX10^+^ cells per layer. This suggests active regulation of cell type composition in the perineuronal space, rather than stochastic distribution (Fig. 7d).

### Macrophages play dynamic role in neuronal soma coverage

Macrophage displayed complex cellular morphologies and were highly entangled with SGCs near the neuronal soma (Fig. 7e-f). To investigate neuronal coverage, we combined models trained to identify and segment the 3D cellular morphology of FASN^+^ SGCs, and IBA1^+^ macrophages, with models trained to identify neuronal somata. We quantified coverage of neurons 1, 5 and 10µm from the neuronal soma as it correlated to onion layers 1, 2, and 3-4 respectively. At 1, 5 and 10 µm SGC coverage was 57.2%, 58.6%, and 56.0% respectively, while macrophages covered the remaining 42.8%, 41.3%, and 44.0% (Fig. 7g). Since the ratio of SGCs/macrophage nuclei remained constant, shifts in neuronal-coverage ratios would thus imply morphological remodeling likely driven by macrophages.

When comparing the coverage ratio of 1μm to 10μm for individual neurons, almost all within-donor neurons showed consistent increasing or decreasing coverage directionality (Fig. 7h). Within donor-neurons were sampled across multiple neuron-islands, suggesting that peri-neuronal macrophage organization is either driven by hDRG parenchymal/systemic cues rather than neuron specific signals, or reflects a common intrinsic neuronal state throughout the entire hDRG.

### CX3CR1^+^ macrophages form highly interconnected networks surrounding neuronal somata

To further investigate macrophage heterogeneity in the hDRG we stained IBA1, CD163, CX3CR1 and DAPI and observed pronounced inter-donor differences in macrophage quantity and morphology (Fig. 7i). On average, 66% of hDRG macrophages were CD163^+^, which closely resembled data we previously published in mouse^18^, and an average of 30% of macrophages were CX3CR1^+^ (Fig. 7j-k). The average distance of CX3CR1^+^ macrophages to neuronal somata was significantly lower than CD163^+^ macrophages, indicating they are the subset intertwined with SGCs surrounding neurons (Fig. 7l). Finally, we noticed that the macrophages on the surface of neurons often formed a net-like structure and connected with one another. To quantify this, we created a connectivity index, where all Iba1 positive signal was segmented by connected components and the number of nuclei for each object became the connectivity number. Individual macrophages were then awarded the connectivity number of the object cluster they came from. CX3CR1^+^ macrophages showed significantly higher connectivity than CD163^+^ macrophages (Fig. 7m).

### Proliferating cells in the hDRG are predominantly immediately adjacent to neurons

Ki67 was stained to identify dividing cells along with SOX10, Iba1 and DAPI to indicate cell type. Ki67^+^ cells showed strong spatial clustering near neurons (Fig. 7n-o), with proliferation significantly enriched in the first layer. Although this is partially explained by higher cell abundance in inner layers, the ratio of dividing cells is significantly higher in layer 1 compared to layer 2 and 7+ respectively (Extended Data Fig. 9d). The majority of Ki67^+^ cells were predominantly IBA1^+^ and layer 1 macrophages constitute 84% of classified proliferative activity, which is striking as SOX10^+^ cells are 4-fold more abundant. There were SOX10^+^Ki67^+^ cells as well, which were also significantly enriched in layer 1 (Fig. 7p).

### Perineuronal macrophage proliferation increases with age

Ki67⁺ cell abundance was strongly correlated with donor’s age in layer 1 macrophages, but not in SGCs. This relationship in layer 1 macrophages was well described by a Gompertz function (R²=0.71 which models cumulative, saturable processes common in aging biology^59,60^ (Fig. 7q). The exponential nature of the response is consistent with age-associated changes in macrophage turnover that may reflect altered functional states^61^.

## Discussion

Human DRGs exhibited substantial donor-to-donor variability across multiple structural and molecular parameters, reflecting a combination of biological heterogeneity and technical factors related to tissue collection, handling, and processing that are inherent to post-mortem human tissue. By systematically evaluating these variables, we identified cryoprotection as a critical determinant of neuronal ultrastructural preservation. On this basis, we propose the Tissue Quality Index as a practical quality-control metric for future studies, with samples scoring below 0.9 warranting cautious interpretation. In addition, our results highlight the need to consider signal diffusion in snap frozen tissues, potentially driven by protein redistribution^33,36^ as well as epitope loss and fixation induced artifacts when designing and interpreting immunohistochemical analyses of hDRG.

After accounting for technical sources of variability through donor-normalized analyses, the observation of larger neuronal somata in women provides a potential explanation for previously reported sex-dependent differences in human pain processing and may contribute to documented differences in nerve conduction velocities^62–64^. While other anatomical factors may contribute to inter-individual variability; our results highlight the importance of donor-normalized analyses when categorizing hDRG neuronal populations.

Neurons within the ganglion displayed robust spatial clustering based on shared protein-expression profiles and soma size, suggesting that hDRG organization is highly structured rather than stochastic. This organization is consistent with developmental patterning principles previously proposed in transcriptomic studies, suggesting that molecular identity and spatial positioning remain linked in the mature hDRG^7,10,65^.

At a higher organizational level, hDRGs are arranged into discrete neuronal islands characterized by dense vascularization, reminiscent of the vascular architecture observed across entire rodent DRGs, including enrichment of permeable microvessels near neurons. This arrangement likely reflects the high metabolic demands of the neuron-SGC-macrophage niche may also create a permissive interface for circulating factors, with potential implications for therapeutic access to the DRG.

Previous studies have identified distinct transcriptional profiles of glial cell and macrophage subpopulations in the hDRG, based on their proximity to neuronal somata^21,23^. Cells in the innermost layer exhibited enlarged nuclei, consistent with increased transcriptional activity associated with direct neuronal support. Macrophages occupying this layer were enriched for CX3CR1, formed interconnected structures and exhibited DRG-wide, coordinated, remodeling/occupancy of the perineuronal space. This suggests the presence of dynamic perineuronal macrophage networks that collectively respond to systemic or tissue-level cues. Such intimate neuron-macrophage interactions have been implicated in pain regulation through mechanisms such as mitochondrial transfer^28,66^. Finally, we observed a strong age-associated increase in proliferation of layer 1 macrophages, indicating increased cellular turnover rather than numerical expansion. This finding may suggest increased metabolic demands and/or decreased perineuronal macrophage function with age. Additionally, macrophage turnover may serve as a quantitative indicator of relative hDRG biological age. Whether chronic pain states or disease-associated conditions further modulate the dynamics of CX3CR1+ perineuronal macrophages remains an important question for future studies.

While derived from a limited number of donors, the organizational principles described here, neuronal clustering, layered perineuronal niches, and vascular specialization, were consistently observed across all samples, supporting their robustness despite inter-individual variability.

By integrating optimized tissue-processing workflows, large-scale antibody benchmarking, quantitative spatial metrics, and deep learning–assisted image analysis, this study establishes a robust and scalable framework for protein-level and spatial interrogation of human DRGs. All analytical pipelines are marker-agnostic and readily adaptable to additional datasets, facilitating reuse across laboratories and future hDRG studies. This resource complements existing transcriptomic atlases by providing spatially resolved protein context and uncovering organizational principles such as neuronal clustering, layered perineuronal niches, and vascular specialization that are not readily accessible through RNA-based approaches alone and provide a scalable reference for future human DRG studies.

## Methods

### Tissue processing of human dorsal root ganglia

Human lumbar DRGs (L3-L5) were collected from 16 different organ donors in Canada following consent from the next of kin (Supplementary Table 5, McGill University Health Centre REB2019-4896). At the time of collection whole DRGs were either immediately frozen on dry ice (iFr) or immediately fixed (iFix) in 4% paraformaldehyde (PFA). iFr DRGs were shipped on dry ice and stored at −80°C. When ready for processing, iFr DRGs were quartered, using a 3D printed blade guide (see supplemental material) on dry ice, taking care to not let the samples thaw. Quartered frozen DRGs were either incubated for 3 hours in 4% PFA (iFr-F) or blocked in O.C.T. (Histolab, ref: 45830), cryosectioned and then fixed on the slide for 30 minutes (iFr-Fos). Whole, iFix DRGs were incubated in 4% PFA for 8 hours and shipped in PBS + 0.02% sodium azide at 4°C. Fixation time for iFix hDRGs experienced higher variability due to tissue collection hour. iFix and iFr-F DRGs were cryoprotected in 30% sucrose with 0.02% sodium azide until tissue sunk (3-5 days) at 4°C. Samples were blocked in O.C.T or NEG50 (Epredia, ref: 6502) and stored at −80°C until cryosectioned. We note that NEG50 resulted in superior section quality and consistency. DRGs were sectioned using a CryoStar NX70 cryostat (20-50 μm). Thinner slides (20 μm) were used for initial protein identification and antibody optimization, while 50 mm sections were utilized during analysis of whole tissue fixed samples across donors 1-15. For donors with both iFix and iFr-F samples, both samples were cryoprotected at the same time and blocked/sectioned together on the same slides.

### Immunohistochemistry of hDRG

We modified IHC treatment for the hDRG using solutions from the iDISCO ^64^ protocol to improve antibody penetration for 50μm thick sections. Additionally, we add a high salt prewash using 0.1M PBS, which decreases tissue autofluorescence. ^64^Tissue sections were thawed at room temperature for 1 hour, then washed twice with 0.1M phosphate-buffered saline (PBS; pH 7.35–7.45 for all PBS solutions (Sigma-Aldrich, ref: P38135) and once with 0.01M PBS for 5 minutes each. Next, sections were incubated at room temperature with a permeabilization solution containing 0.01M PBS, 0.16% Triton X-100 (Sigma-Aldrich cat: X100-100ML), 20% dimethyl sulfoxide (DMSO) (Sigma-Aldrich, cat: D4540-100ML), and 23 g/L glycine (Sigma, cat: G8898-500G) for 30 minutes. Following permeabilization, a blocking solution consisting of 0.01M PBS, 0.2% Triton X-100, 6% normal donkey serum (Millipore, cat: S30-100ML) and 10% DMSO was applied for 1 hour.

Primary antibodies (details provided in Supplementary Table 2) were applied overnight at room temperature in an antibody solution composed of 0.01M PBS, 0.2% Tween-20 (Sigma-Aldrich, cat: P9416-100ML), 10 mg/L heparin (Leo Pharma, ATC-code: B01AB01), 0.02% sodium azide, 5% DMSO, and 3% normal donkey serum. After incubation, slides were washed with a washing solution (0.01M PBS, 0.2% Tween-20, 10 mg/L heparin, and 0.02% sodium azide): a brief 1-minute wash followed by two 5-minute washes.

Tissue sections were then incubated with the appropriate donkey secondary antibodies (Supplementary Table 2). After secondary antibody incubation, sections were washed three times using the washing solution (5 minutes each), followed by incubation with DAPI (Invitrogen, ref: D21490) (1:10,000) for 10 minutes. Additional washes were performed: once with the washing solution for 5 minutes, followed by 0.1M PBS and 0.01M PBS washes. Slides were dried in the dark for 5 minutes and mounted with ProLong Gold Antifade Mountant (Invitrogen, cat: P36934).

### Lipofuscin quenching protocols

Two different lipofuscin quenchers were tested. Sudan Black B (Sigma-Aldrich, ref: 199664-25G) was incubated for 1 hour used at 0.2%, diluted in 70% EtOH, as a pretreatment. True Black Plus (Biotium, ref: 23014) was used as a post-treatment before mounting, applied for 30 min diluted 1:40 in PBS, after which 3 five minutes PBS washes were applied to the samples. All neuronal markers were tested and analyzed with True Black Plus to decrease false positives.

### Tyramide signal amplification (TSA)

When the regular protocol did not generate a clear signal, the effect of Tyramide Signal Amplification (TSA) (Akoya Biosciences, Ref: NEL741001KT) was assessed. Slides were allowed to equilibrate at room temperature for 3 hours before being incubated overnight with the primary antibody diluted in 0.01M PBS containing 0.3% Triton X-100. Following primary antibody incubation, slides were washed in TNT buffer (0.05% Tween-20 in 0.1M Tris and 0.15M NaCl, pH 7.4–7.45) for 30 minutes. They were then incubated in TNB buffer (prepared with 0.05 g blocking reagent from the PerkinElmer Life Sciences kit, #NEL741) for 30 minutes, followed by a 30-minute incubation with horseradish peroxidase (HRP)-conjugated secondary antibody (DAKO, HRP Rabbit, 1:200) diluted in TNB buffer. Afterward, slides were washed in TNT buffer for 30 minutes and incubated for 15 minutes with Tyramide-Fluorescein solution (1:100 in Amplification Diluent) that had been centrifuged at 14,000 rpm for 10 minutes at 4°C. This was followed by another 30-minute wash in TNT buffer and three additional 5-minute washes in 0.01M PBS. Slides were then incubated for 1 hour in a blocking solution composed of 0.01M PBS, 0.3% Triton X-100, and 3% normal donkey serum (NDS). For non-amplified targets, primary antibodies were added and incubated overnight in a solution containing 0.1% Triton X-100 and 1% NDS. Primary antibody washes consisted of two 5-minute rinses in 0.01M PBS. Secondary antibodies (1:300 dilution in antibody solution) were applied and incubated for 1.5 hours. Slides were briefly rinsed in PBS, followed by a 10-minute incubation with DAPI (1:10,000). Finally, slides underwent three 5-minute washes in 0.01M PBS before being mounted with ProLong Gold Antifade Mountant.

### Data analysis

#### Tissue Quality Index (TQI)

The Tissue Quality Index was defined as the percentage of observed somatic tissue volume relative to the full predicted neuron volume. For iFr-F and iFix samples, full soma volume was estimated using the pan-cytosolic marker NF200 and the DRG-quant pipeline^67^. Due to performance limitations of iFr-Fos predictions, full soma was estimated by manual annotation looking while observing NF200 and DAPI channels. Observed somatic tissue volume was calculated by subtracting shrunken volume from the full volume. A pixel was classified as shrunken if it met two criteria: (1) NF200 signal intensity below 50% of the non-neuron background mean for that sample, and (2) location within a 5-pixel diameter sphere where >50% of pixels met criterion 1.

### Comparison of whole tissue fixation immediately fixed or frozen

To compare fixation protocols, IHC was performed on DRGs from the same donor. One DRG was processed using tissue fixed immediately after DRG collection while the matched DRG was frozen at time of collection. Sections from both samples were mounted on the same slide to ensure comparable conditions. Paired DRGs from 6 donors were used. Representative images of each marker and DAPI were captured with optimized settings for each donor, ensuring identical settings were applied to each pair. The images were exported as TIFF format and blinded for scoring. Three independent researchers evaluated the staining quality based on signal intensity, specificity, and background noise on a 0-100% scale. Representative images shown were selected based on proximity to the average score for each group/ Representative images of paired donor samples were selected to showcase the expected differences for each marker. Statistical significance was assessed with a paired T-test and adjusted for False Discovery Rate using two-stage step-up method of Benjamini, Krieger and Yekutieli.

### Manual quantification of neuronal markers

After confirming antibody human specificity and a visual confirmation of positive staining, we quantify single sections stained for the 20 neuronal proteins. Whole section images of the stained samples were taken with confocal microscopy with an open pinhole (80μm). Neurons were then manually identified with help of the selected markers, background noise, and DAPI signal by an experienced researcher. The neurons were classified in three different categories based on protein expression: Negative neurons had no visual difference with unspecific signal; detectable neurons had enough signal to distinguish them from background staining; and lastly, high expressors were identified after an overview of general neuronal expression, as those with a clear higher expression than the rest of the neurons. For nuclear markers, only neurons with a visible nucleus were quantified, while neuronal selection criteria for non-nuclear proteins included all neurons with clear somata without large coverage of non-neuronal cells.

### RNA sequencing data analysis

Due to the low number of neurons with a single scRNAseq study, and to have a good representation of the different human neuronal subsets, we integrated neuronal scRNAseq studies from all the current hDRG studies on RStudio (R version 4.4.1). Neurons proceeding from single soma RNA sequencing^10^ filtered into a Seurat object for genes expressed in at least 3 cells, with a minimum of 200 features per cell. The cells were later filtered for cells with between 4500 and 14000 RNA features, and RNA count of 600000. After normalization, this dataset was then merged with three previously filtered and integrated^8^ scRNAseq human studies. To integrate these datasets, previously integrated datasets were re-transformed into human gene symbols using biomaRt in a similar manner as for the original transformation into mouse symbols. All data was normalized based on study, with 2000 variable genes as anchors for integration. The Seurat object was then scaled, and the first 20 PCs were selected (based on elbow Plot) to run the UMAP and find neighbors. The selection of cluster resolution was based on the clustree plugin. The clusters were later compared to the original publications and manually labeled, mapped based on common genes with described populations in the large-scale RNA atlas study of hDRGs^9^.

To facilitate comparison to protein data, the percentage of neurons expressing more than 0.5 RNA counts was retrieved for the proteins that were manually evaluated for protein expression.

For non-neuronal cells, a single dataset was used^20^, filtered for RNA features between 600 and 5000, with a count lower than 15000, and mitochondrial percentage under 5%. Data was normalized, integrated between samples, scaled, and the first 24 PCs were used for the UMAP generation. After finding neighbors and identifying cluster resolution, the clusters were identified based on marker genes for each population. Immune cells were then separated, re-normalized, and scaled, with a subplot using the first 15 PCs for UMAP generation and finding neighbors. Cluster resolution was informed with clustree, with marker genes allowing the identification of the main populations of immune cells in the hDRG.

For data visualization, the generated UMAPS were plotted using scCustomize package on RStudio.

### Image Acquisition

Images were acquired on a Zeiss LSM800 laser-scanning confocal microscope equipped with four lasers (405, 488, 561, and 640 nm). Tissue overviews were taken with a 10x air objective. Quantification images were either taken with a 20x air objective, 40x water immersion objective, or 63x oil immersion objective.

### iDISCO clearing of hDRG

One hDRG that was glutaraldehyde fixed was quartered as described above. Since the sample was glutaraldehyde fixed it was not stained and followed the strndard iDISCO protocol^68^. In brief, tissue was serially dehydrated with methanol in H2O, 20%, 40%, 60%, 80%,100%,100% for 1 hour each. The sample was then incubated for 3 hours with Dichloromethane (DCM) at room temperature. Followed by two 15-minute washes in 100% DCM. The DRG was then placed in DiBenzyl Ether and imaged with with the LaVision Light Sheet microscope (Biotech, UltraMicroscope II). The extremely high level of autofluorescence from lipofuscin in glutaraldehyde fixed tissue was used to identify neuronal soma and observe the neuron-islands in the hDRG.

### Image Analysis

All raw CZI images were processed and run through the DRGquant pipeline^67^ to organize, run model prediction, and set up analysis. Any Unet models not described below were trained using yapic, as described in DRGquant. Image analysis was performed on workstations running Ubuntu (linux), with >128GB ram and NVIDIA GPUs (RTX3090 or RTX6000).

### Analysis of neuron-stained panels

Neuron locations were predicted using a two-stage meta-learning ensemble architecture implemented in the MONAI framework (MONAI 1.3.0, PyTorch 2.0.1, Python 3.9.15)^69^. In Stage 1, three channel-specific Attention gated U-Nets were trained independently, with two input channels, each having nuclei stain (DAPI) to ensure nuclei-based information, and one of the 3 IHC based stains to capture cell type specific information. In Stage 2, a meta-learner Attention U-Net received the 4 original channels concatenated with the 3 Stage 1 probability maps (7 channels total) to generate refined predictions by learning optimal integration of the base models.

The Attention U-Net architecture comprised encoder channels (32→64→128 filters) with 2*strided convolution down sampling and 3×3 convolutional kernels, a 256-filter bottleneck, and a symmetric decoder with attention-gated skip connections. Models were trained using the Dice Focal loss function with AdamW optimization (learning rate 2*10⁻⁴, weight decay 1*10⁻⁵) on 352*352-pixel patches with cosine annealing scheduling. Data augmentation included, horizontal/vertical flips, 90° rotations, additive Gaussian noise (σ=0.05), and contrast adjustment (γ ∈ [0.7, 1.3]).

Five-fold cross-validation was performed with independent training on each fold. For inference, images were processed with all 5-fold models at 9 spatial offsets (stride/3 grid), generating 45 predictions per pixel. Final masks were created by consensus voting with optimal thresholds determined via cross-validation resulting in a Raw prediction. All Stage 1 models for all panels had a Dice Focal better than 90% while all stage 2 had a Dice Focal better than 95%.

Raw AI predictions were post-processed using ImageJ/Fiji (v1.54p) with CLIJ2 package (v2.5.3.1). Binary predictions underwent morphological refinement and non-disjunct objects separation was done by a size and shape adaptive algorithm based under the assumption of high compactness objects.

Inter-soma hole regions were identified by thresholding each channel at a channel consensus of signal weaker than background-mean/4 (where background was defined as tissue regions outside neurons). Defined hole regions were included in size calculations, while excluded in signal ratio calculations. Neurons with hole ratios ≥0.5 were retained.

Neuron subpopulation classification was done using the same logic as in DRG-quant^67^, where expression was related to statistics of the surrounding tissue region. Detectable signal was defined by ∼4 times a minimum expected random signal in relation to background (based on neuron size, >10^-15^% chance to happen randomly), high expressors was classified by ∼30 times higher and very high expressors ∼180 times higher the expected random signal. Data collection from labelled images was done using CLIJ2_statisticsOfLabelledPixels and morphometry metrics was done by 3D ImageJ Suite’s 3D Manager.

Inter-neuronal distances were calculated by estimated edge-to-edge measurements. For each neuron pair, edge-to-edge distance was computed as: d_edge_ = d_center_ - r₁ - r₂ where d_center_ is the Euclidean distance between centroids, and r₁, r₂ are approximated neuronal radii.

### Distance analysis by Cross-type nearest neighbor distance comparison

To assess whether annotated subpopulations exhibit spatial clustering within tissue volumes, we implemented an edge-based cross-type nearest-neighbor comparison approach. For each subpopulation (e.g., A+) within each donor sample, we calculated two metrics: (i) the within-group distance, defined as the mean edge-to-edge Euclidean distance from each A+ object to its three nearest A+ neighbors, averaged across all A+ objects; and (ii) the cross-group distance, defined as the mean edge-to-edge Euclidean distance from each A-object to its three nearest A+ neighbors, averaged across all A-objects. Edge-to-edge distances were calculated by closest Euclidian distance from the estimated object edges. This yielded a single paired measurement per donor: a within-group distance and a cross-group distance for each subpopulation. Spatial clustering is indicated by systematically lower within-group distances compared to cross-group distances, reflecting that objects of the same subpopulation are positioned closer to each other than objects from the complementary population.

### Onion layers algorithm for perineuronal nuclear categorization

Nuclei were categorized into concentric layers around individual neurons using the “onion layers algorithm” (Extended Data Fig. 8). For each nucleus, spatial territories were defined via Voronoi tessellation. Layer assignment proceeded iteratively by testing if a nucleus with any Voronoi volume fell within an iteration specific boundary volume. The first boundary was generated by eroding the neuron mask by an estimate of non-neuronal nuclei (r_nucleus_), with subsequent boundaries created by dilating the first boundary volume by 2*r_nucleus_ per iteration. Assigned nuclei were excluded from subsequent iterations, and remaining nuclei formed the outermost layer. To reduce unnecessary computational time, we limited our number of onion layers to 6, where all additional nuclei/cells were classified as 7+.

Onion layers algorithm properties. This approach ensures: mutually exclusive layer assignment (each nucleus assigned to exactly one layer), preservation of spatial relationships through Voronoi territories rather than point distances alone, shape-adaptive boundaries that follow the neuron morphology rather than assuming circular symmetry.

## Data availability

The scRNAseq data used in this study are publicly available from previous studies in the Gene Expression Omnibus (GEO), for single soma dataset was available under accession number GSE273557 and previously integrated human dorsal root ganglia datasets can be accessed at https://painseq.shinyapps.io/harmonized_painseq_v1/. Non-neuronal scRNAseq used can be accessed under accession number GSE169301.

## Code availability

Code recourse for the hDRG protein atlas workflow, that is easily adapted with complete documentation is available at: https://github.com/CamillaSvensson/Human-DRG-Protein-Atlas-Workflow. Raw code regarding RNAseq analysis and image analysis for all the panels used is available at https://github.com/CamillaSvensson/Molecular_pain_research.

## Supporting information

hDRG Cutter

## Acknowledgements

We would thank the McGill Scoliosis and Spine team for providing the hDRGs.

## Author information

**Authors affiliations**

**Department of Physiology and Pharmacology, Karolinska Institutet, Stockholm, Sweden**

Matthew Adam Hunt, Sven David Arvidsson, Gustaf Wängberg, Zhening Zhang, Zerina Kurtovic, Camilla I Svensson

**Faculty of Dental Medicine and Oral Health Sciences, Alan Edwards Centre for Research on Pain, McGill University, Montreal, QC, Canada**

Emerson Krock

**Division of Orthopaedic Surgery, Department of Surgery, McGill University, Montreal, QC, Canada**

Lisbet Haglund

## Contributions

M.H, S.D.A, and C.I.S. conceived the project and designed the experiments. C.I.S. supervised the work. E.K. and L.H. provided the human dorsal root ganglia and controlled for the initial collection protocol. M.H. processed the samples and prepared tissue slides. S.D.A and Z.Z. compiled antibody information and performed immunofluorescence stainings. S.D.A performed confocal imaging, initial evaluation of antibody performance, and neuronal protein manual quantification. Z.Z., Z.K., and E.K. scored the antibody signal. S.D.A. and Z.K. contributed to RNAseq analysis of publicly available data. G.W. developed the novel algorithms and neural network architecture used in this study, performed image analysis, and generated deep learning models for neuronal panels. M.H. and G.W. jointly developed deep learning models and conducted image analysis for non-neuronal panels. G.W. prepared the online analysis resource. M.H., S.D.A., G.W., and C.I.S. wrote the paper with assistance from all authors.

## Corresponding author

Correspondence to Camilla I. Svensson

## Ethics declaration

Human lumbar DRGs were obtained from organ donors through a collaboration with Transplant Quebec. All procedures were approved by and performed in accordance with the ethical review board at McGill University (McGill University Health Centre REB 2019-4896) and the Swedish Ethics Review Board (ethics permit number 2025-05433). Familial consent was obtained for each subject.

---

**Supplementary Fig. 1.**
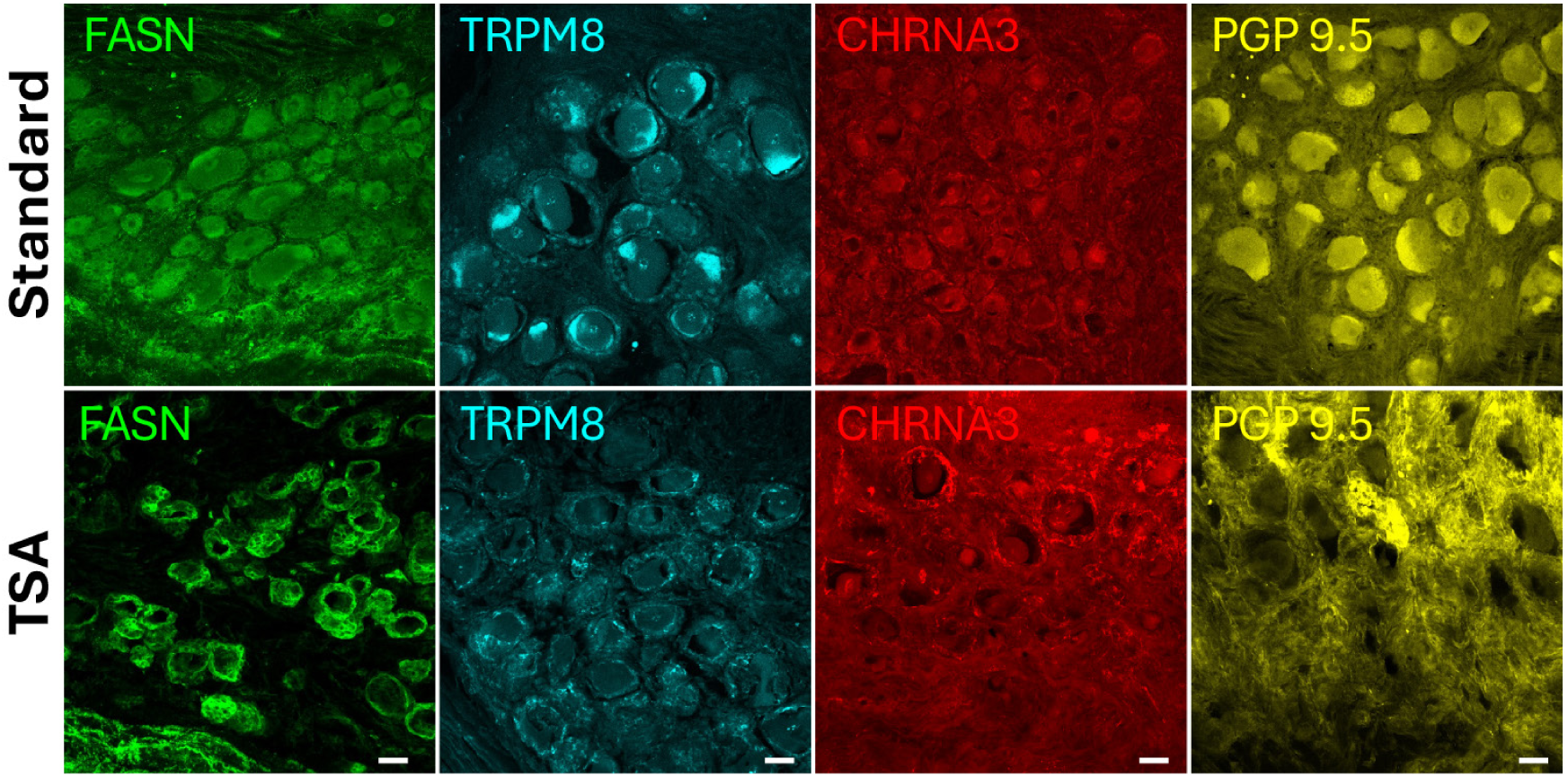
TSA effect on sensitive antibodies. Representative images of neuron rich regions with FASN (green), TRPM8 (cyan), CHRNA3 (red) and PGP 9.5 (yellow) using a regular protocol (top row) or TSA (bottom row). Scale bar = 50 μm

**Extended Data Fig. 1.**
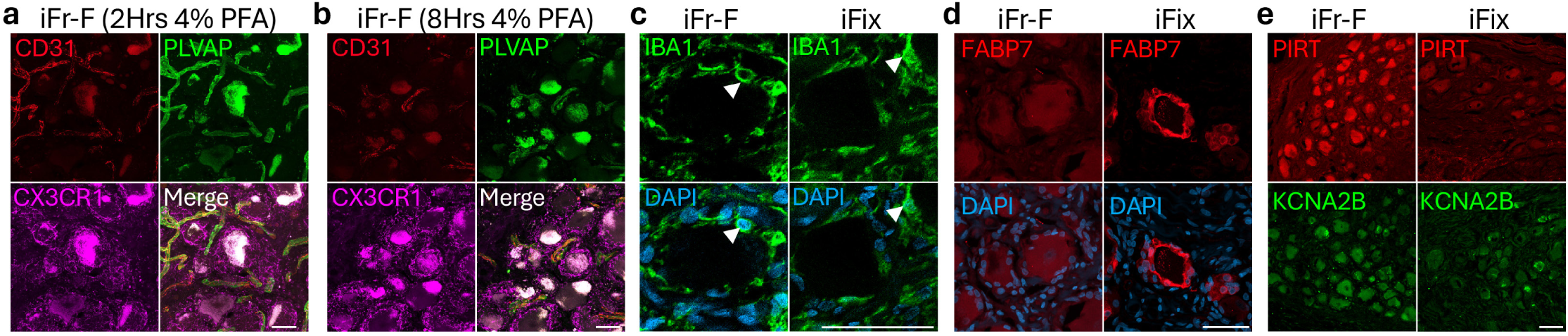
Fixation time affects antibody specificity. a, Staining pattern for CD31 (red), PLVAP (green), CX3CR1 (magenta) after 2 hours of whole tissue fixation of iFr tissue. b, Paired donor staining for CD31, PLVAP, CX3CR1 after 8 hours of whole tissue fixation. Scale bar = 50 μm. c, Staining pattern for IBA1 (green) and DAPI (blue) in iFr-F and iFix samples from the same donor. Arrows highlight the change in macrophage nuclear region. d, FABP7 (rabbit, cat: PA5-24949) and DAPI (blue) staining showing perineuronal staining in paired donor iFr-F and iFix samples showing loss of SGC staining in iFr-F samples. e, Paired donor images of iFr-F and iFix samples for PIRT (red) and KCNA2B (green) taken with equal laser settings highlighting a weak staining in iFix samples. Scale bars = 50 μm.

**Extended Data Fig. 2.**
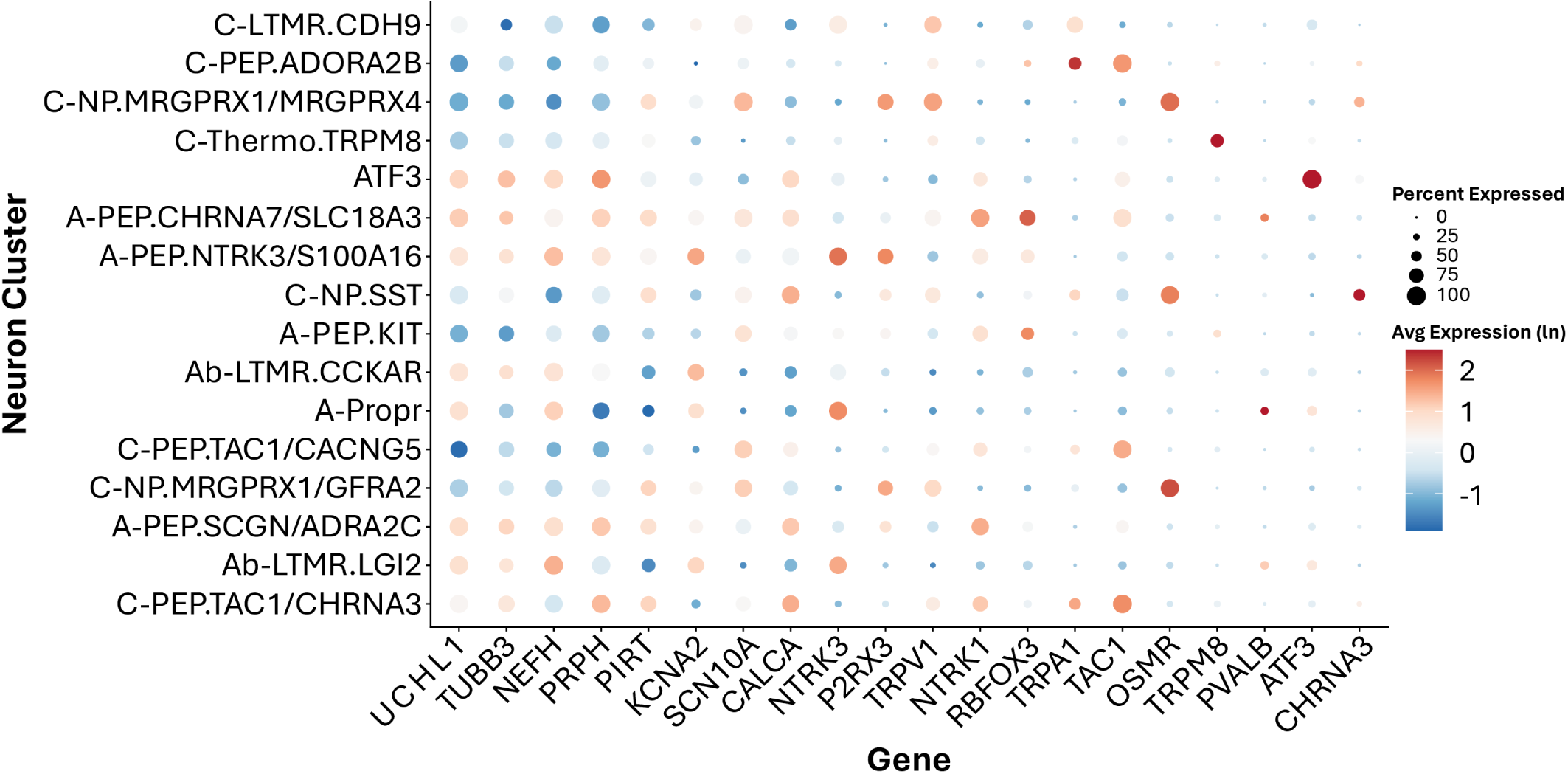
Neuronal RNA differential expression of selected markers. Dotplot showing average log normalized RNA expression for each neuronal subpopulation. The selected genes correspond to RNA transcripts for manually quantified neuronal markers.

**Extended Data Fig. 3.**
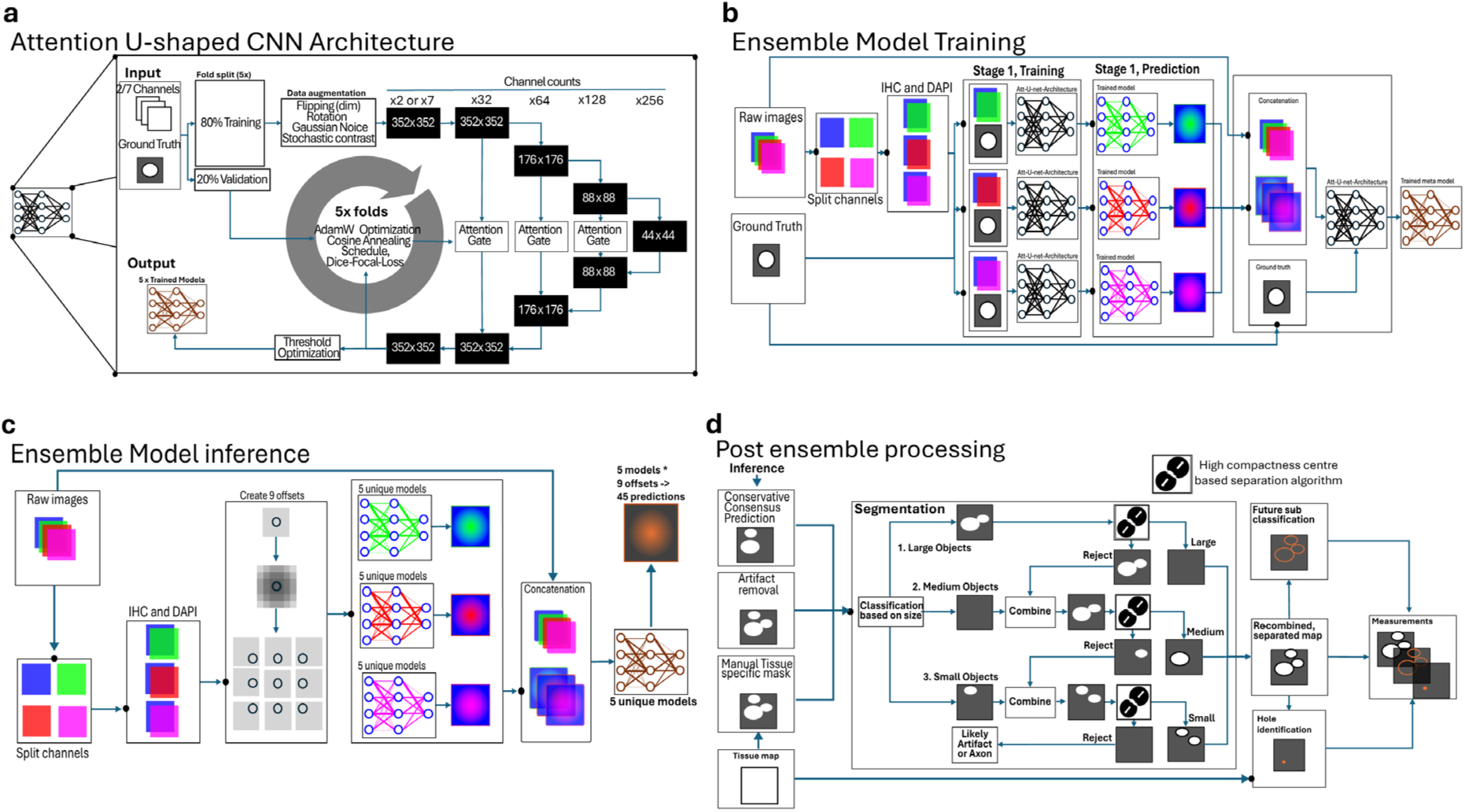
Neural network-based image segmentation pipeline. a, Neural network architecture. Input consists of either 2-or 7-channel images and binary ground truth masks. Data are split 80/20 for 5-fold cross-validation. Each fold is trained independently on augmented data using an attention-gated U-Net architecture with fold-specific threshold optimization. b, Two-stage ensemble model training. All images were acquired with DAPI and three different IHC stains. Stage 1: Models were trained on paired combinations of DAPI with each IHC channel using binary ground truth labels, generating 15 models total (3 IHC-DAPI pairs 5 folds). Each model outputs a prediction map. Stage 2: An ensemble network is trained using the three prediction maps from each Stage 1-fold plus the four original image channels (7 channels total). c, Ensemble prediction strategy. Each of the five ensemble models predicts independently using a tiled approach with nine different shifted windows, yielding 45 predictions per pixel (5 models * 9 windows) to improve contextual information and robustness. d, Post-processing and mask refinement. The 45-layer prediction stack is thresholded to generate a binary mask, then filtered using a manually drawn tissue boundary outlining the dorsal root ganglion (DRG). Sequential filters are applied, including edge-preserving operations. Masks are categorized by size (pixel count). Assuming high object compactness within each size category, objects are eroded to achieve separation, followed by acceptance or rejection based on morphological criteria. Finally, region-specific annotations are applied (e.g., tissue holes, membrane regions).

**Extended Data Fig. 4.**
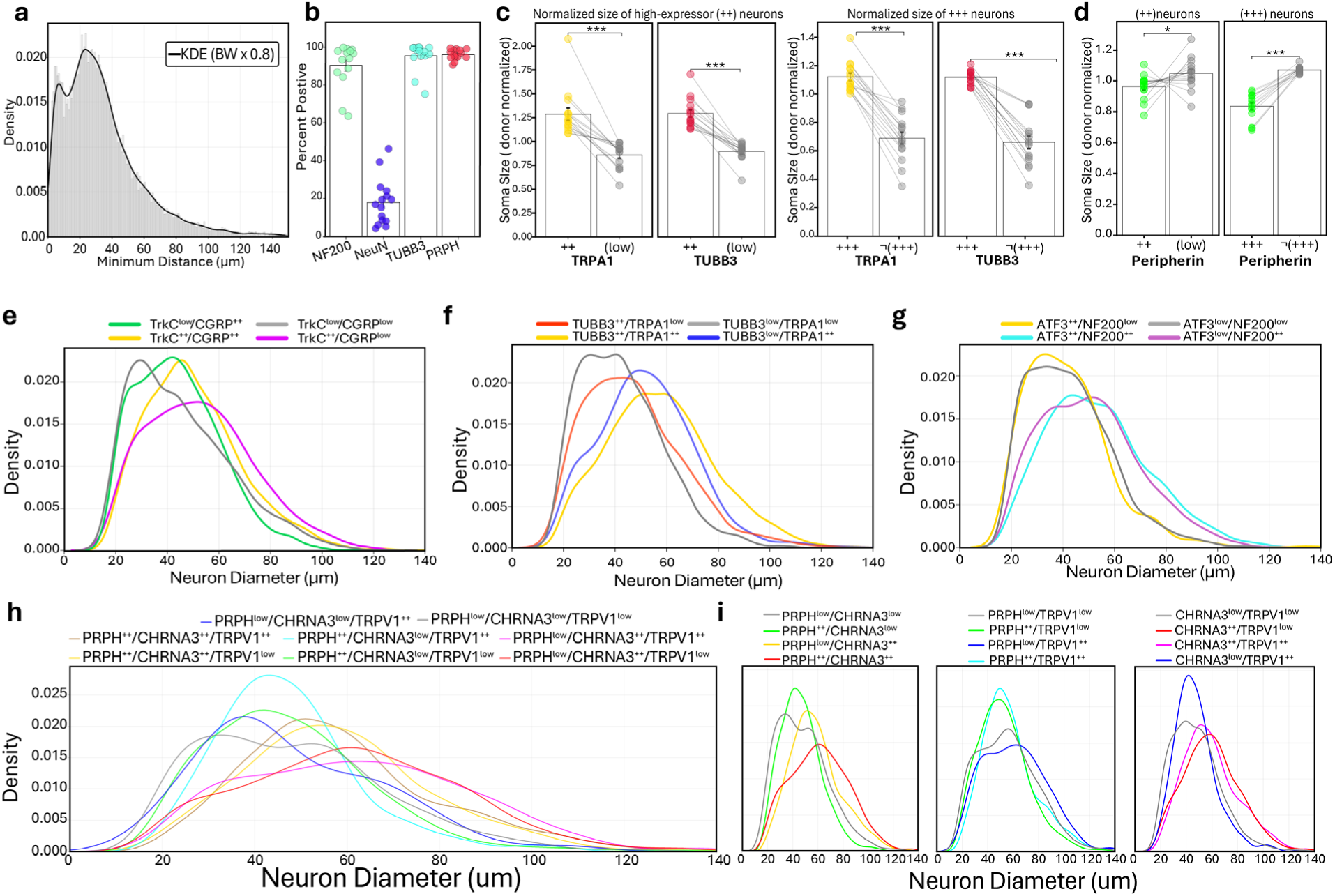
Additional morphometric and spatial analyses of human DRG neurons. a, Histogram (gray) and kernel density estimate (black line) showing the distribution of minimum distances between Neurons in the hDRG. b, Percentage of higher than detectable signal at neurons for 4 putative pan-neuronal markers across 15 donors. c, Normalized neuron size for TRPA1 and β3-tubulin high expressors (++) and very high expressors (+++) versus the rest low/not very high ¬(+++); Wilcoxon signed-rank test (two-sided); \**P* < 0.05, \*\**P* < 0.01, \*\*\**P* < 0.001. d, Normalized neuron size for peripherin high expressors (++) and very high expressors (+++) versus the rest low/ ¬(+++); Wilcoxon signed-rank test (two-sided); \**P* < 0.05, \*\**P* < 0.01, \*\*\**P* < 0.001. e-h KDE curves of neuronal soma diameter for high expressor subpopulations, e, TRKC and CGRP n=9223; f, TUBB3 and TRPA1 n=10408; g, ATF3 NF200 n=8298; h, PRPH, CHRNA3 and TRPV1 n=7972.e-h KDE curves of neuronal soma diameter for high expressor subpopulations, e, TRKC and CGRP n=9,223; f, TUBB3 and TRPA1 n=10,408; g, ATF3 NF200 n=8,298; h, PRPH, CHRNA3 and TRPV1 n=7,972.

**Extended data Fig. 5.**
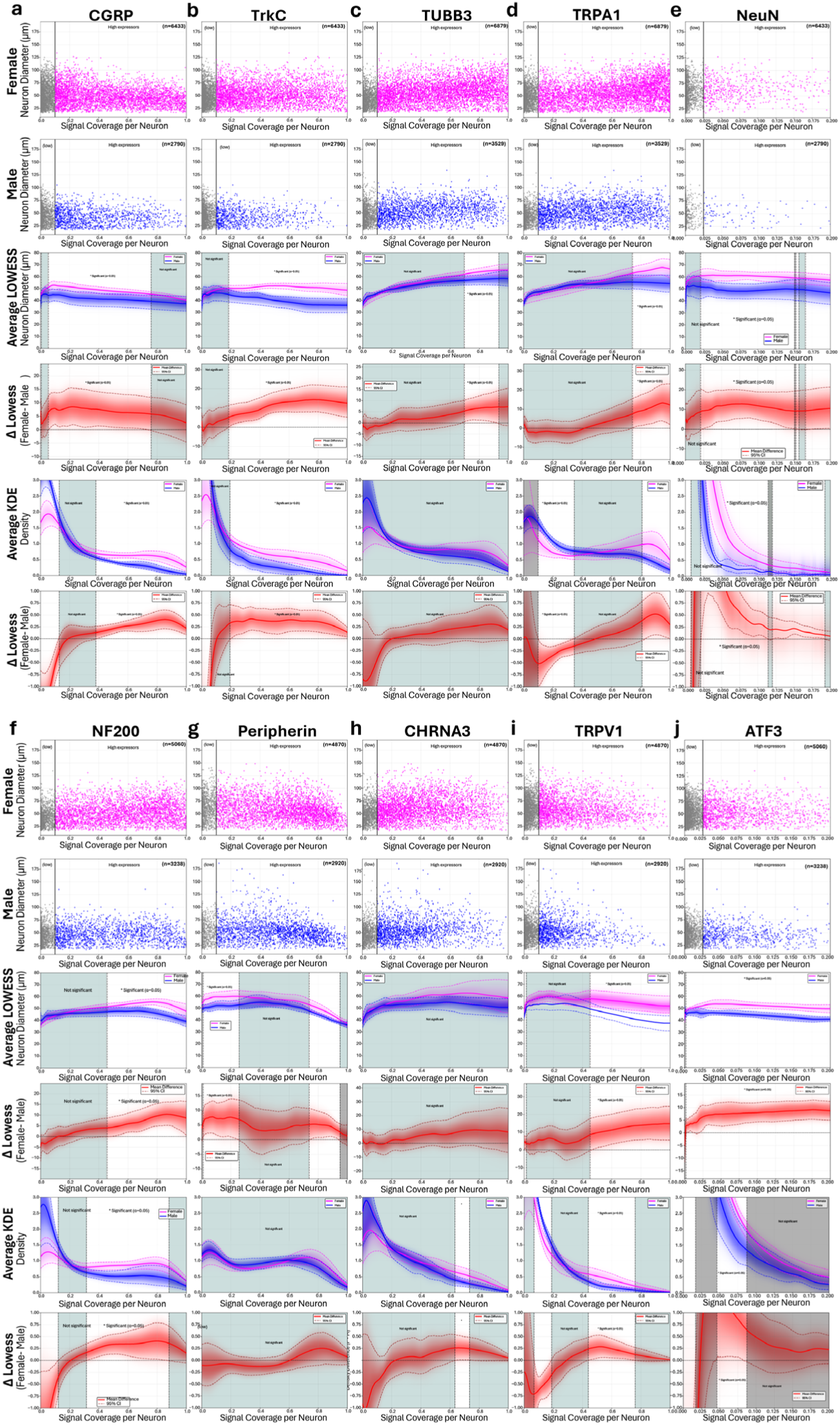
Marker specific neuronal sex-dependent differences in the hDRGa. CGRP, b, TrkC, c, TUBB3, d, TRPA1, e, NeuN, f, NF200, g, Peripherin, h, CHRNA3, i, TRPV1, j, ATF3. Row 1-2, Raw data distributions showing individual neurons from male (blue) and female (red) donors. Row 3, Locally weighted scatterplot smoothing (LOWESS) regression curves (solid lines) with 95% confidence intervals (dashed lines) derived from bootstrap resampling of donor-level LOWESS fits (n=1,000 iterations). Density clouds represent bootstrap density distributions. Gray background shading indicates regions where sex differences are not statistically significant (Percentile Bootstrap Method, 95% CI of the difference includes zero). Vertical dashed lines mark transitions between significant and non-significant regions. Row 4, Difference donor-level LOWESS fits (Female-Male) with 95% bootstrap confidence intervals (red=mean difference, dark red dashed=CI bounds). Red density cloud shows bootstrap distribution of the difference Kernel density estimation (KDE) of marker expression distributions with 95% bootstrap confidence intervals. Density clouds and significance annotations as in the left figure. Row 5, Kernel density estimation (KDE) of marker expression distributions with 95% bootstrap confidence intervals. Density clouds and significance annotations as in the third row. Row 6, Difference in KDE distributions (Female minus Male) with 95% confidence intervals. The red density cloud shows bootstrap distribution. Horizontal dashed line at zero indicates no difference. Significance regions annotated as in previous panels. All statistical comparisons account for donor-level variability through bootstrap resampling at the donor level rather than individual neuron level.

**Extended Data Fig. 6.**
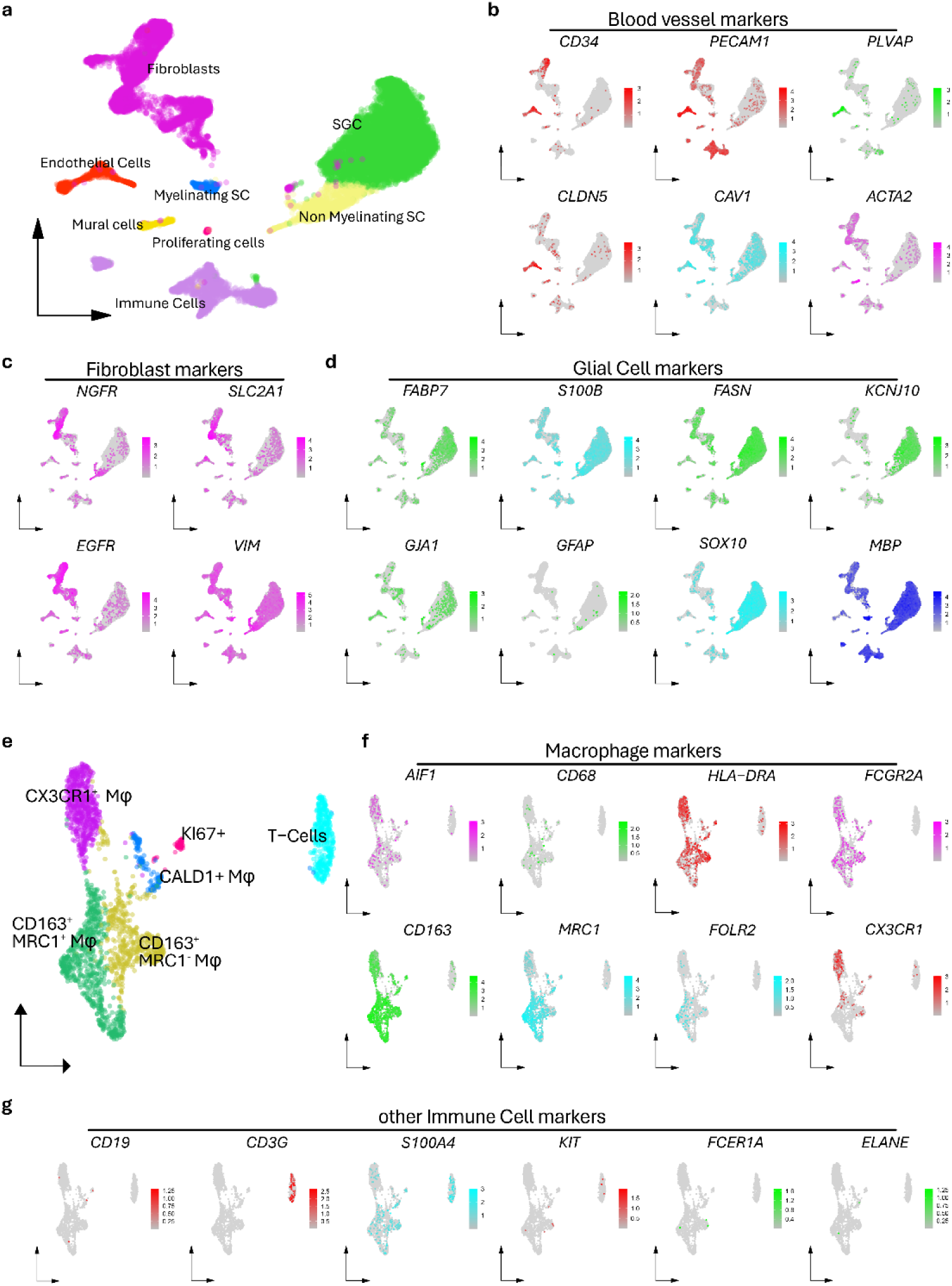
Non neuronal cells in the hDRG. a,UMAP showing cell type clustering in the hDRG. b-d, Feature plots showing RNA expression of markers for blood vessels (b), fibroblasts (c), and glial cells (d). e, UMAP subsetting immune cells showing different immune cell clusters. f-g, Feature plots of immune cell subset showing RNA expression of markers for macrophages (f) and other immune cells (g).

**Extended Data Fig. 7.**
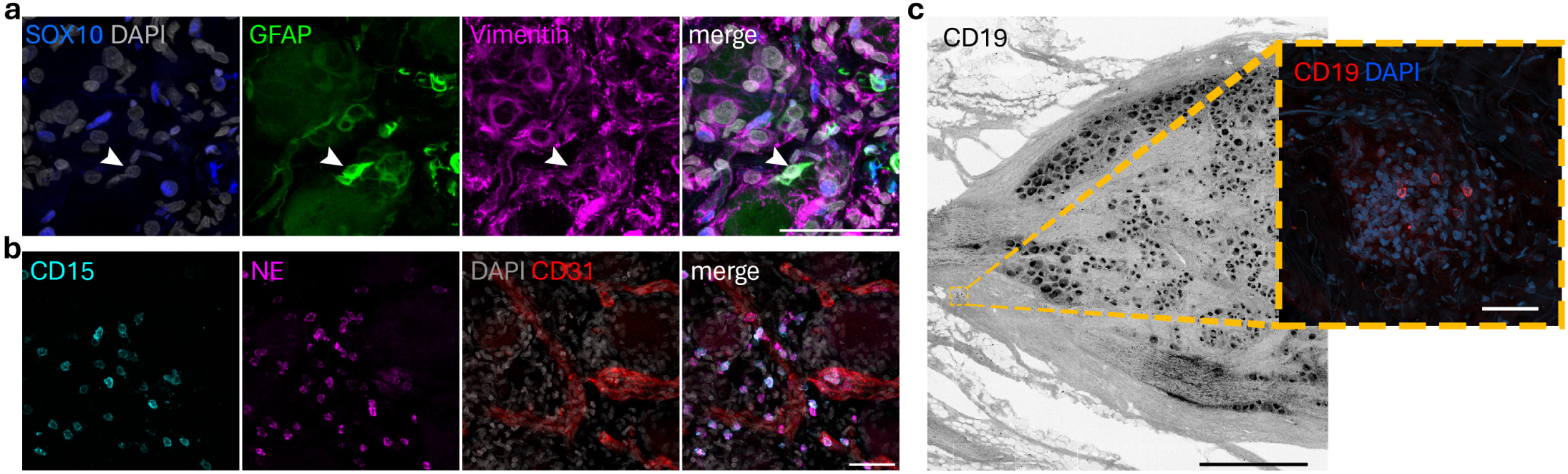
Unexpected marker location. a, Costaining of classical glial and fibroblast markers showing events of Sox10-GFAP+ cells (white arrow), and broad expression of vimentin across cell types. b, Sample with high neutrophil number in a neuron-island. To the left, overview image of neuron-island for nuclear marker (DAPI, Blue), and neutrophil markers CD15 (cyan), and NE (magenta), along with CD31-based masking of blood vessels (red), showing parenchymal localization of neutrophils as well as vascular c, Overview image of a DRG section where lipofuscin highlights neurons, and closeup image of a capsule border region showing clustering of immune cells, including B Cells (CD19, red) among others black scale bar: 500 μm, white scale bars: 50 μm.

**Extended Data Fig. 8.**
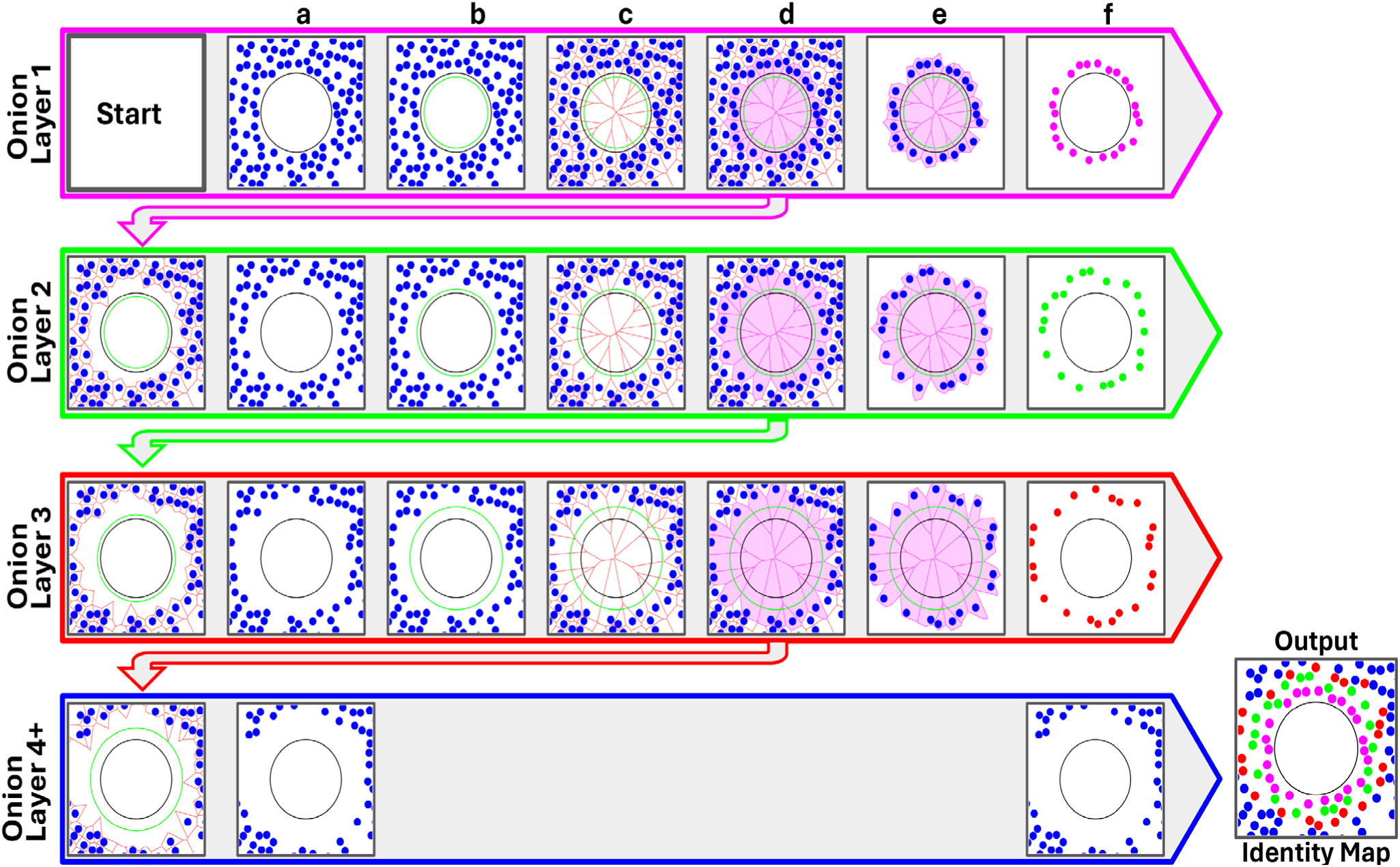
Cyclical onion-layer algorithm for spatial nuclei classification. The onion-layer algorithm classifies nuclei surrounding individual neurons into concentric n-spheres. Algorithm inputs are a nuclei label mask and a neuron label mask. The algorithm iterates for a specified number of layers (4 in this example). a, Input maps: nuclei label mask (blue dots) and neuron label mask (black circle) are defined for the current layer. b, The competitive zone for the current layer (green circle) is defined by dilating the neuron label mask by (LC − 1.5) * Dn, where LC is the layer count and Dn is the estimated average nucleus diameter. c, Current nuclei map used for Voronoi tessellation (red borders). d, The map is partitioned into two subsets: nuclei with Voronoi regions overlapping the competitive zone, and nuclei outside this zone. The latter serves as input for the subsequent layer. e, Overlapping nuclei are reset to their original unlabeled state. f, The classified nuclei for the current layer are saved as an independent label map.

**Extended Data Fig. 9.**
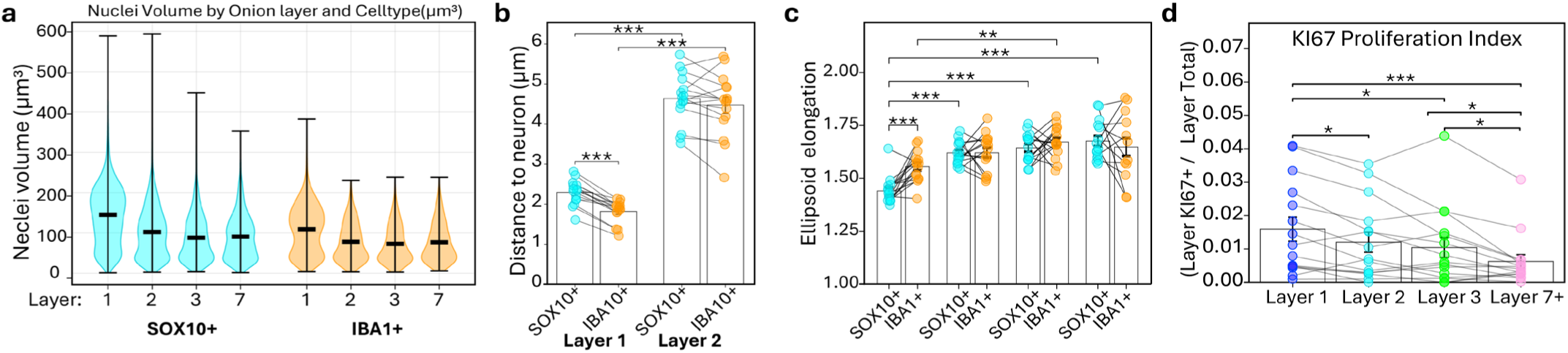
SOX10+ and IBA1+ cell morphology, distance, and proliferation across perineuronal layers. a, Nuclei volume distribution of SOX10+ and IBA1+ cells by onion layer. Violins show full distribution; lines indicate means. b, Distance from SOX10+ and IBA1+ nuclei to nearest neuron surface by perineuronal layer; donor means paired. Wilcoxon signed-rank test with FDR correction; \**P* < 0.05, \*\**P* < 0.01, \*\*\**P* < 0.001. c, Nuclei elongation and compactness of SOX10+ and IBA1+ cells across onion layers. Points represent donor means; bars show mean ± standard error mean; lines connect paired samples. Wilcoxon signed-rank test (two-sided) with FDR correction; \**P* < 0.05, \*\**P* < 0.01, \*\*\**P* < 0.001. d, Proportion of proliferating (KI67+) cells by onion ring, normalized to total KI67+ cells per donor; all cell types pooled. Points show individual donors; bars show mean ± standard error mean. Wilcoxon signed-rank test; \**P* < 0.05, \*\**P* < 0.01, \*\*\**P* < 0.001.

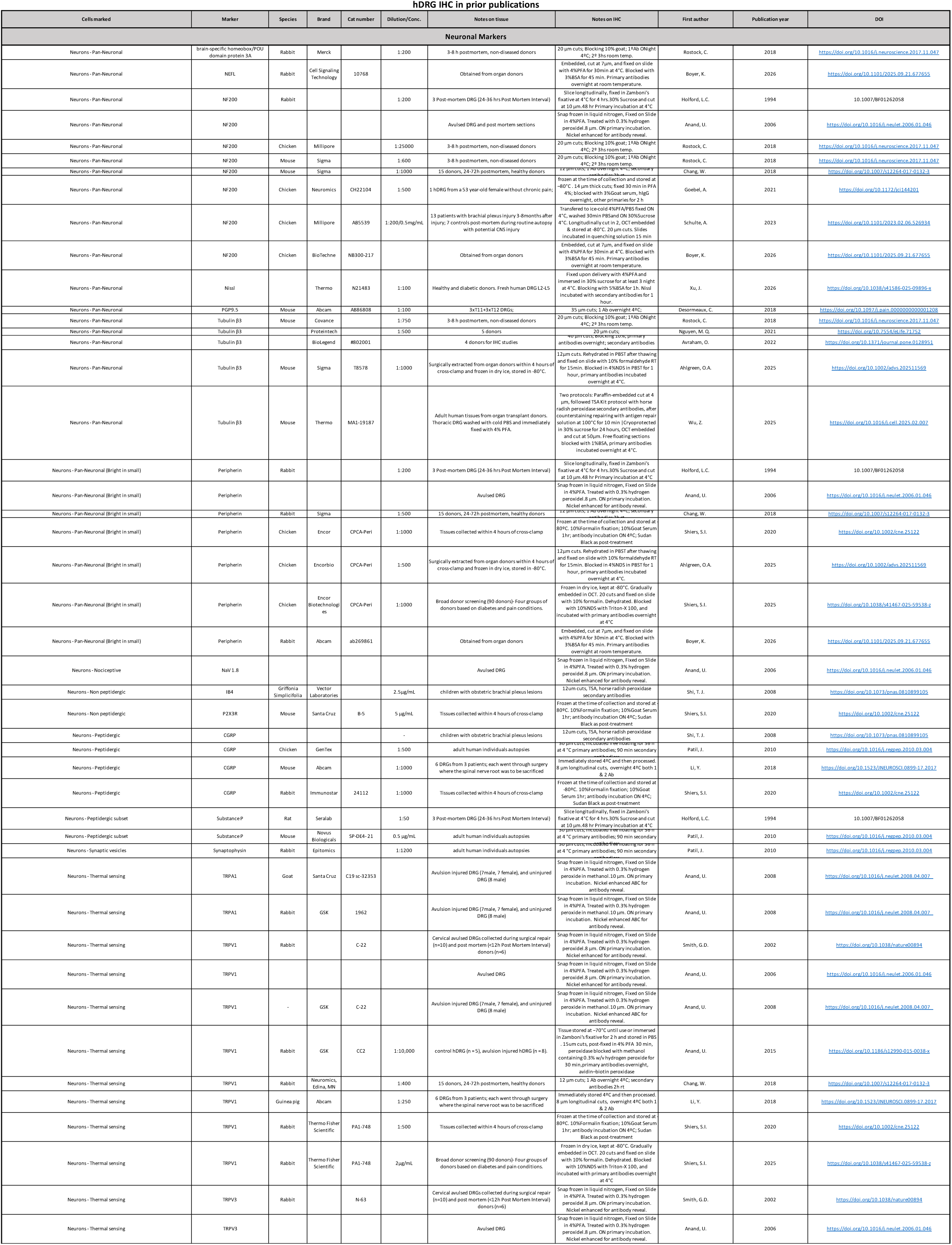
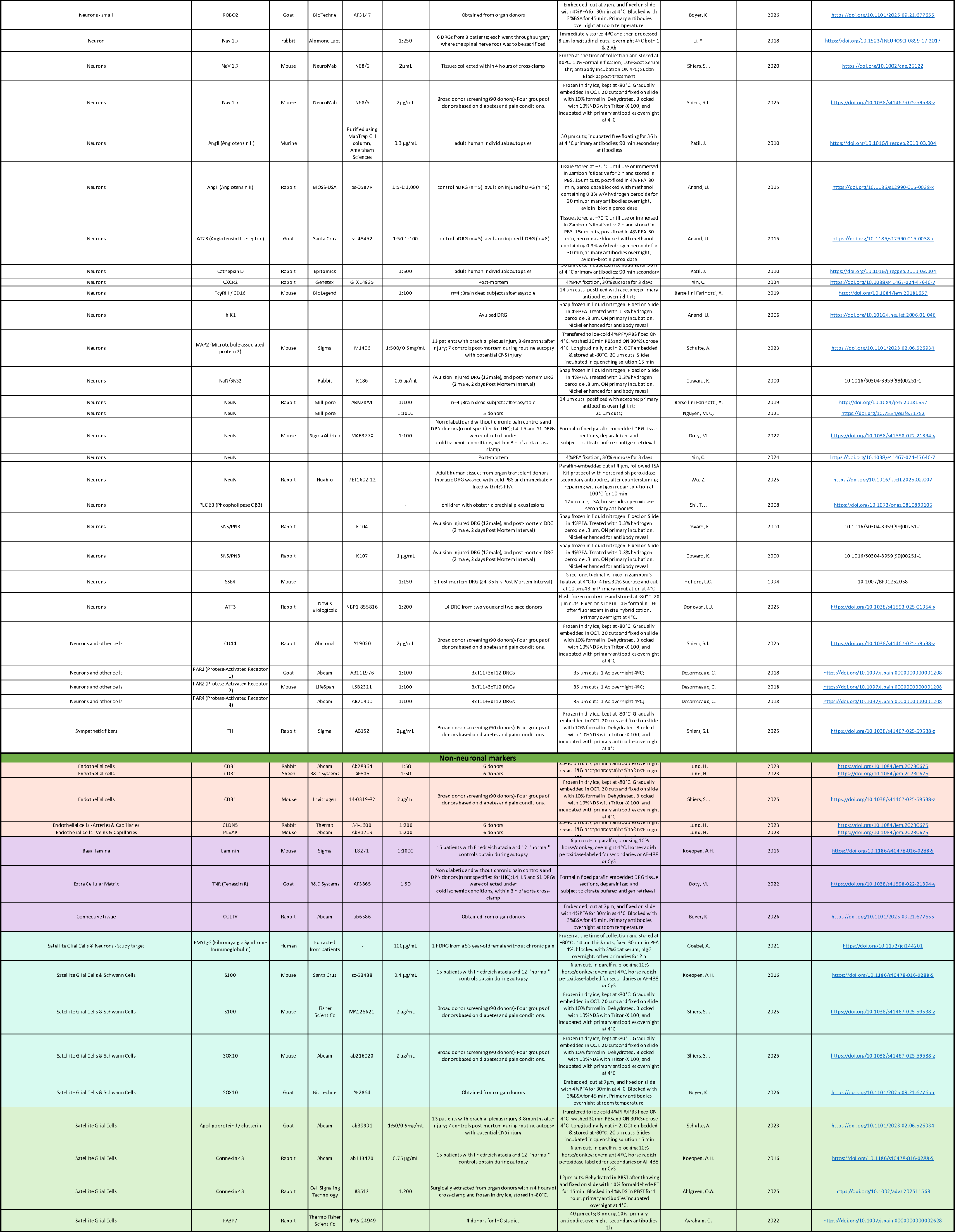
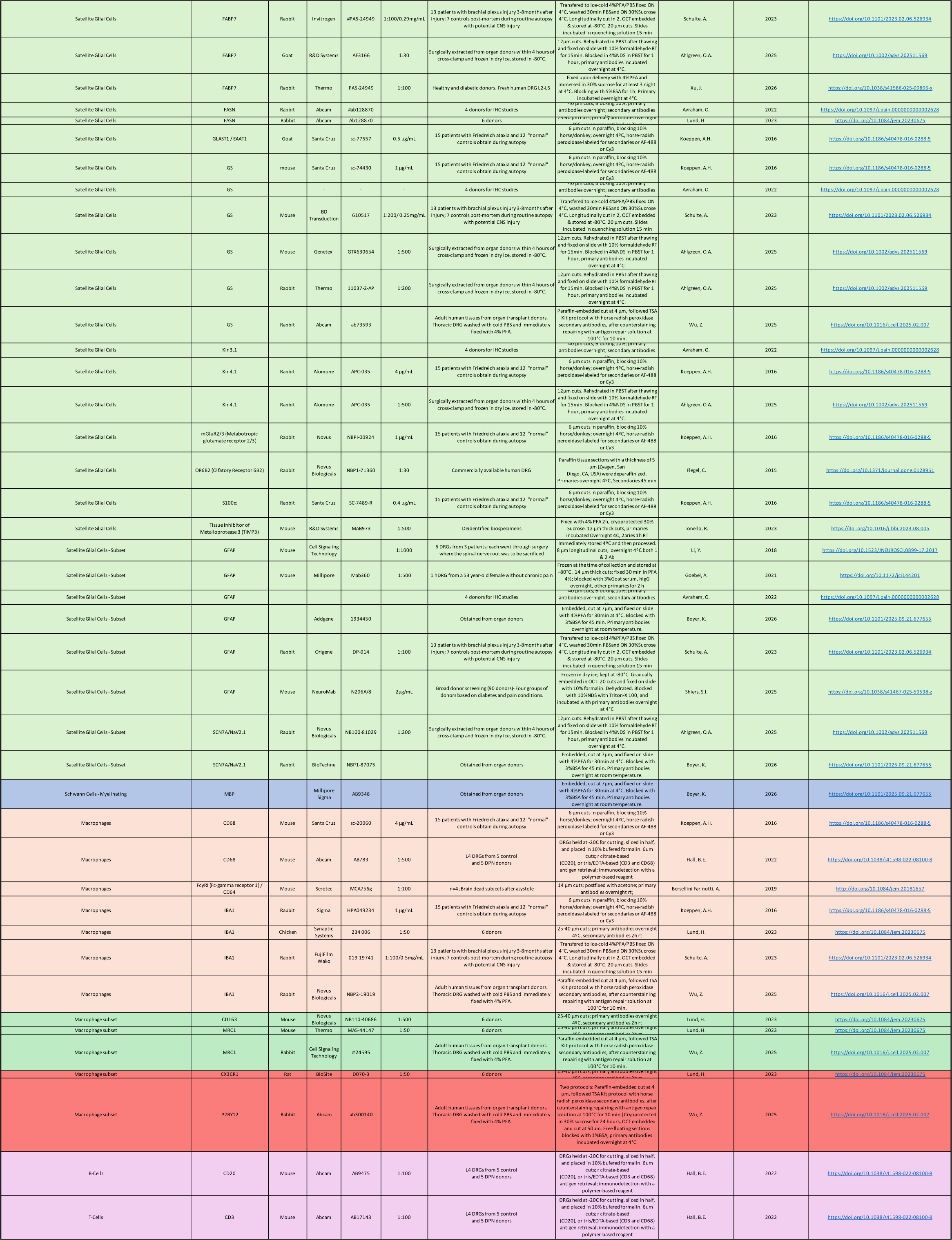
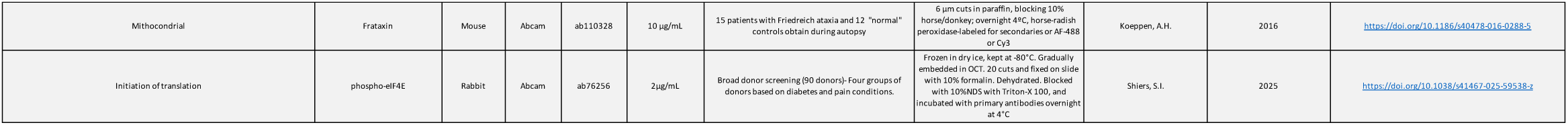

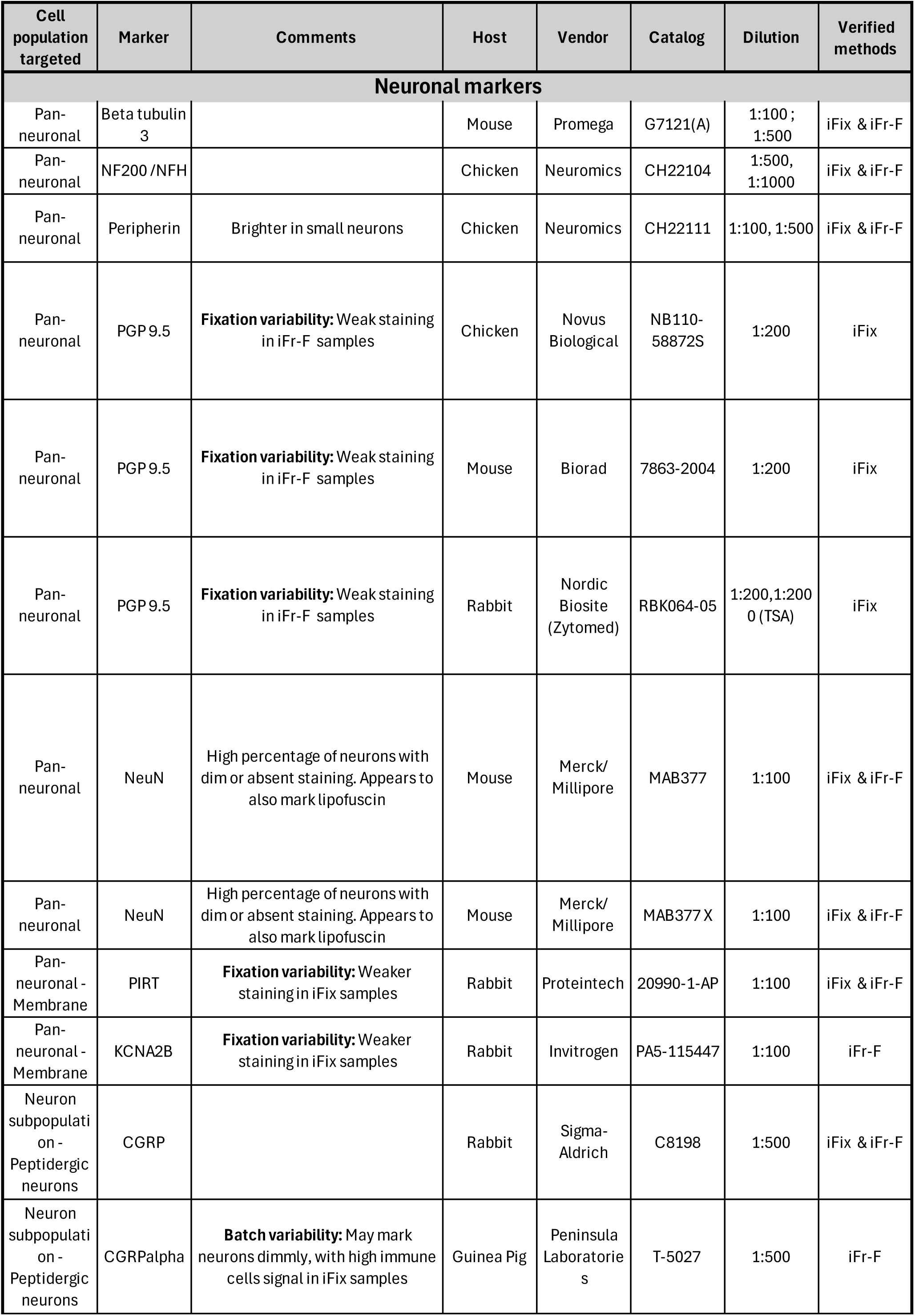
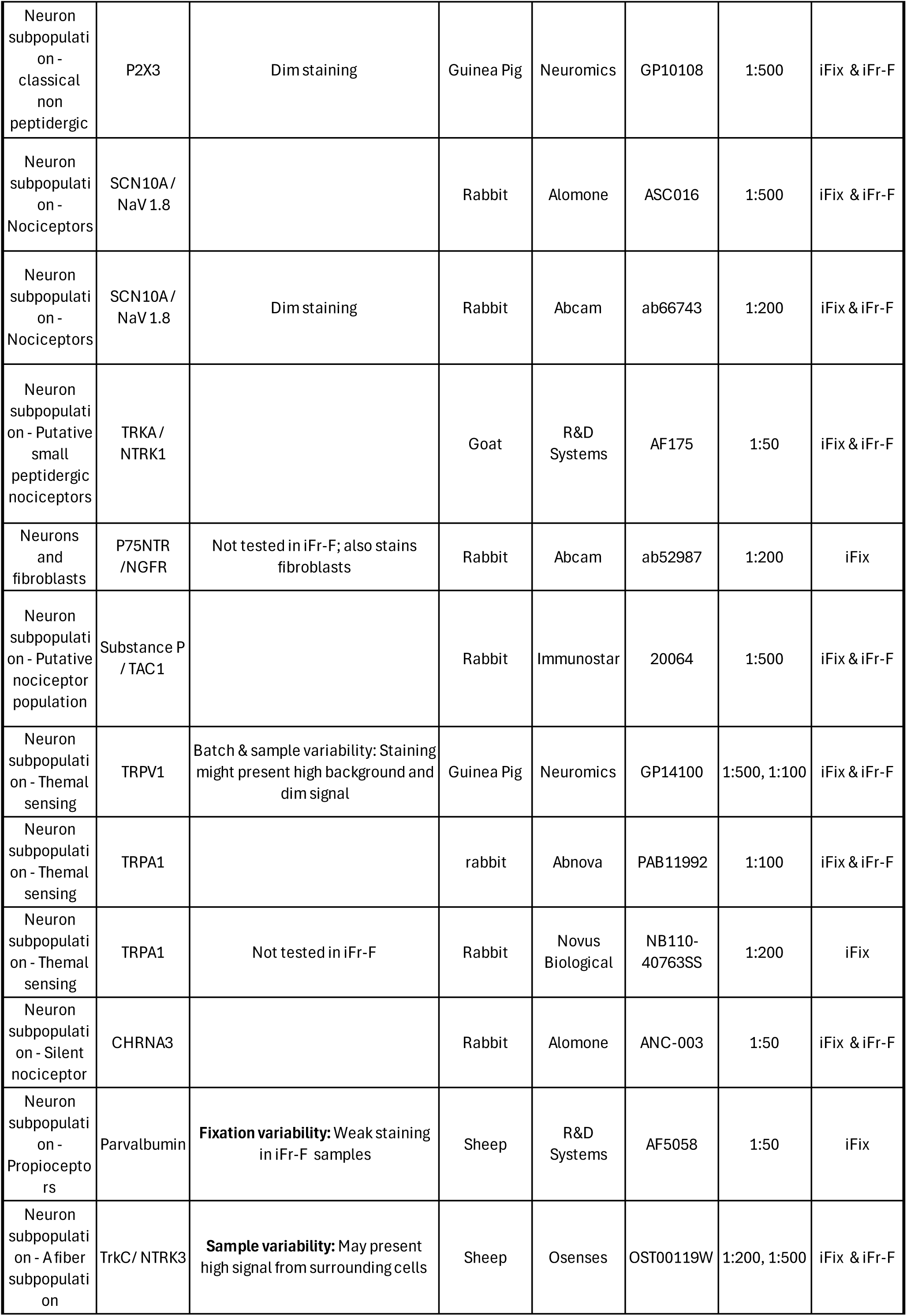
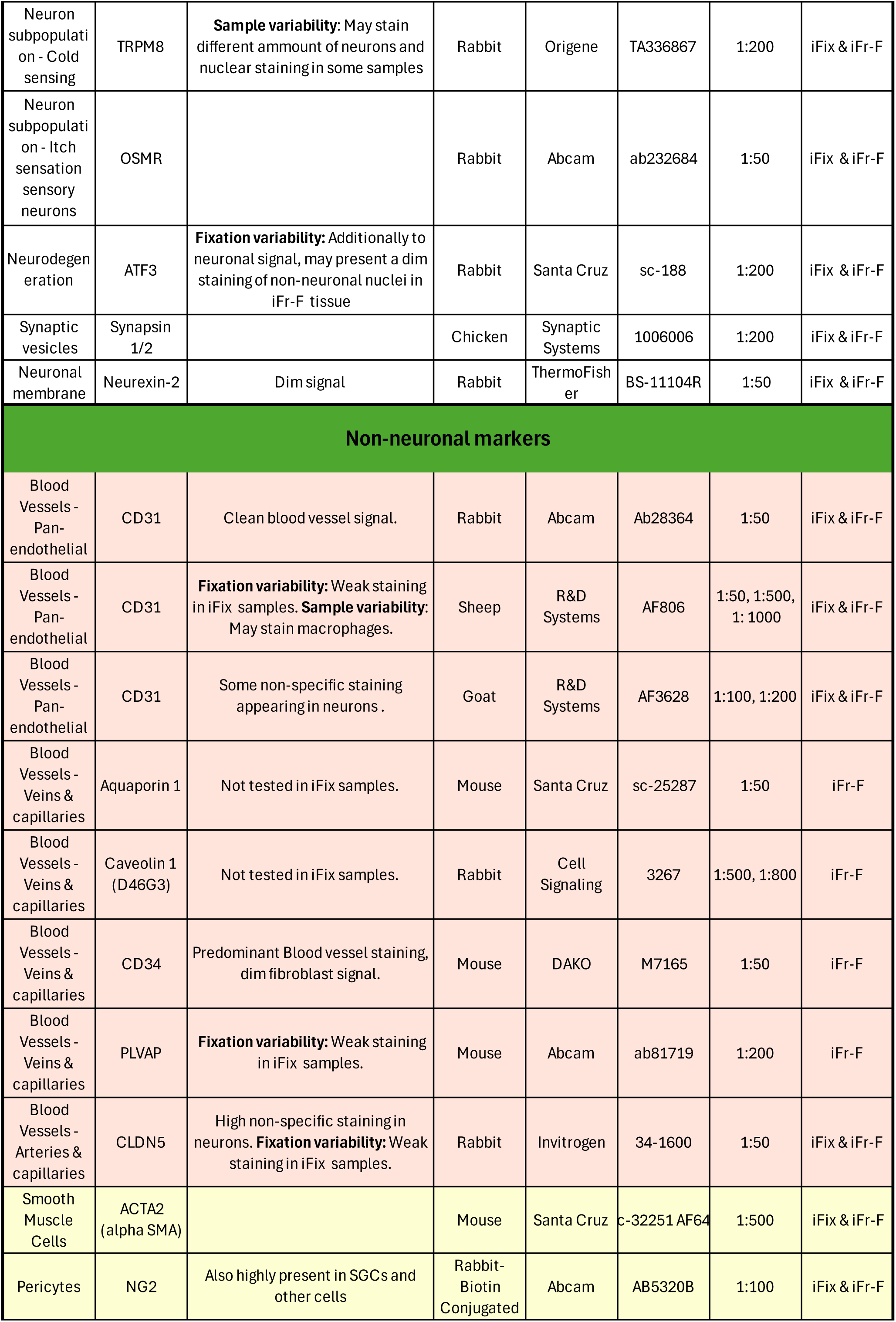
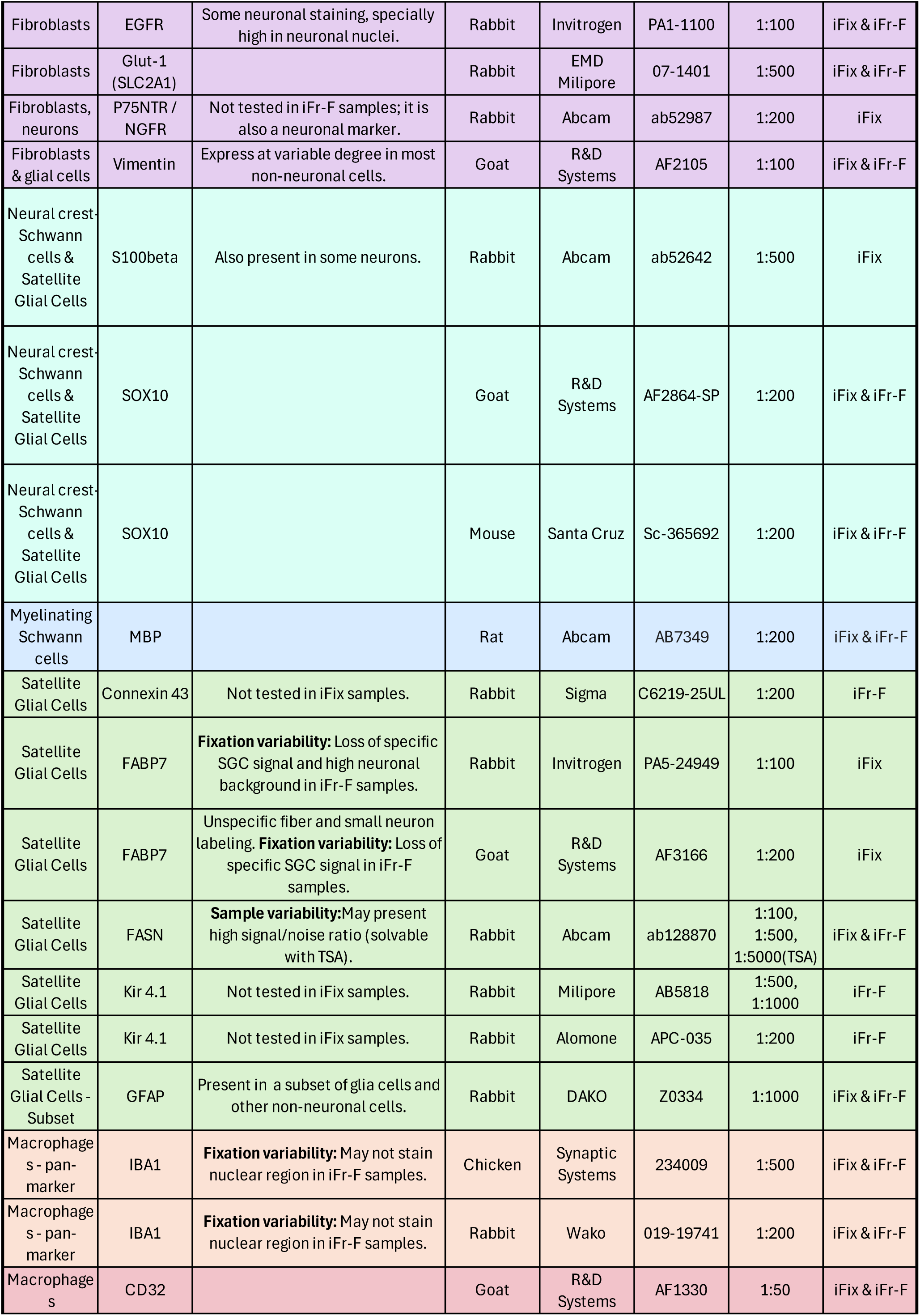
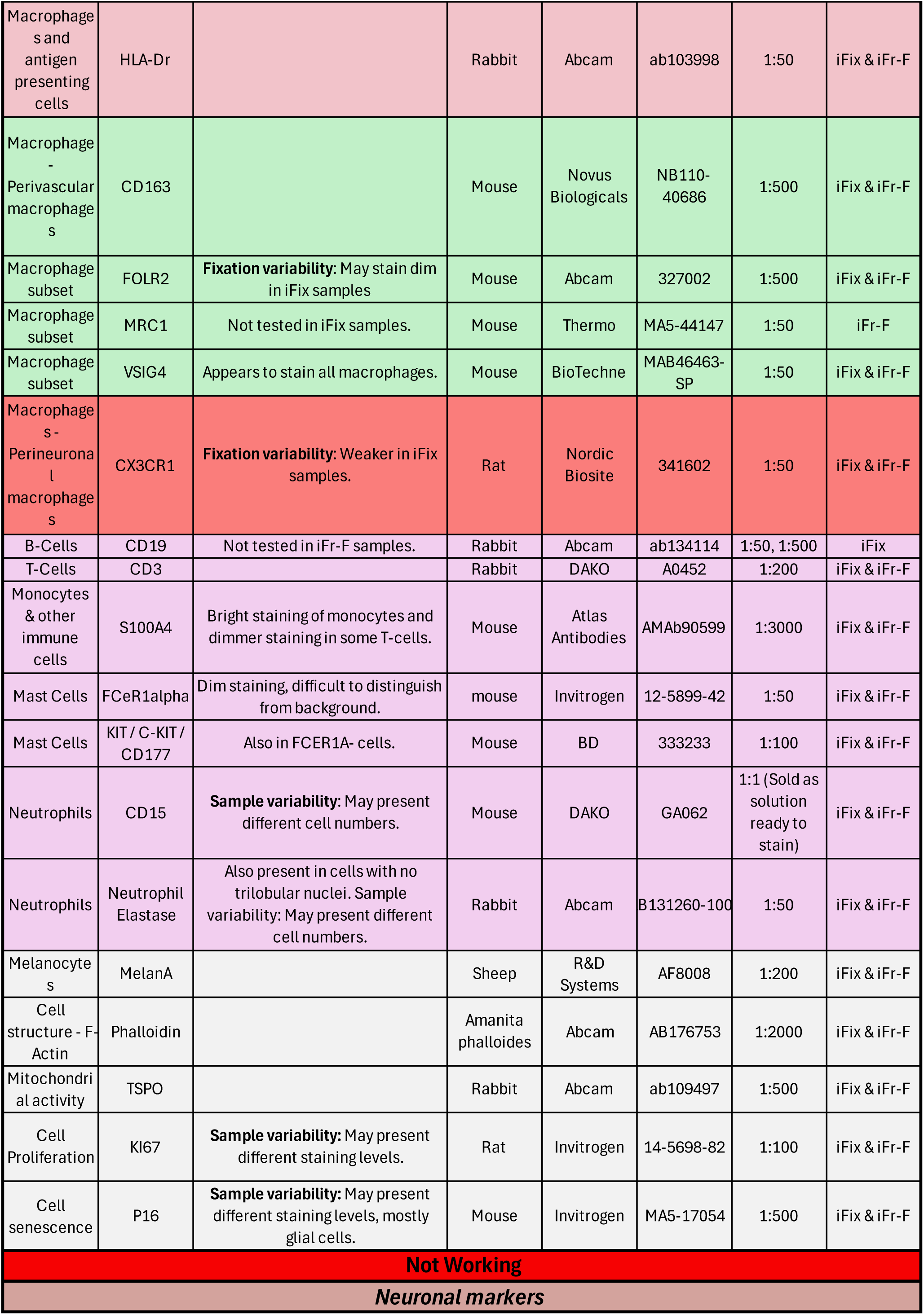
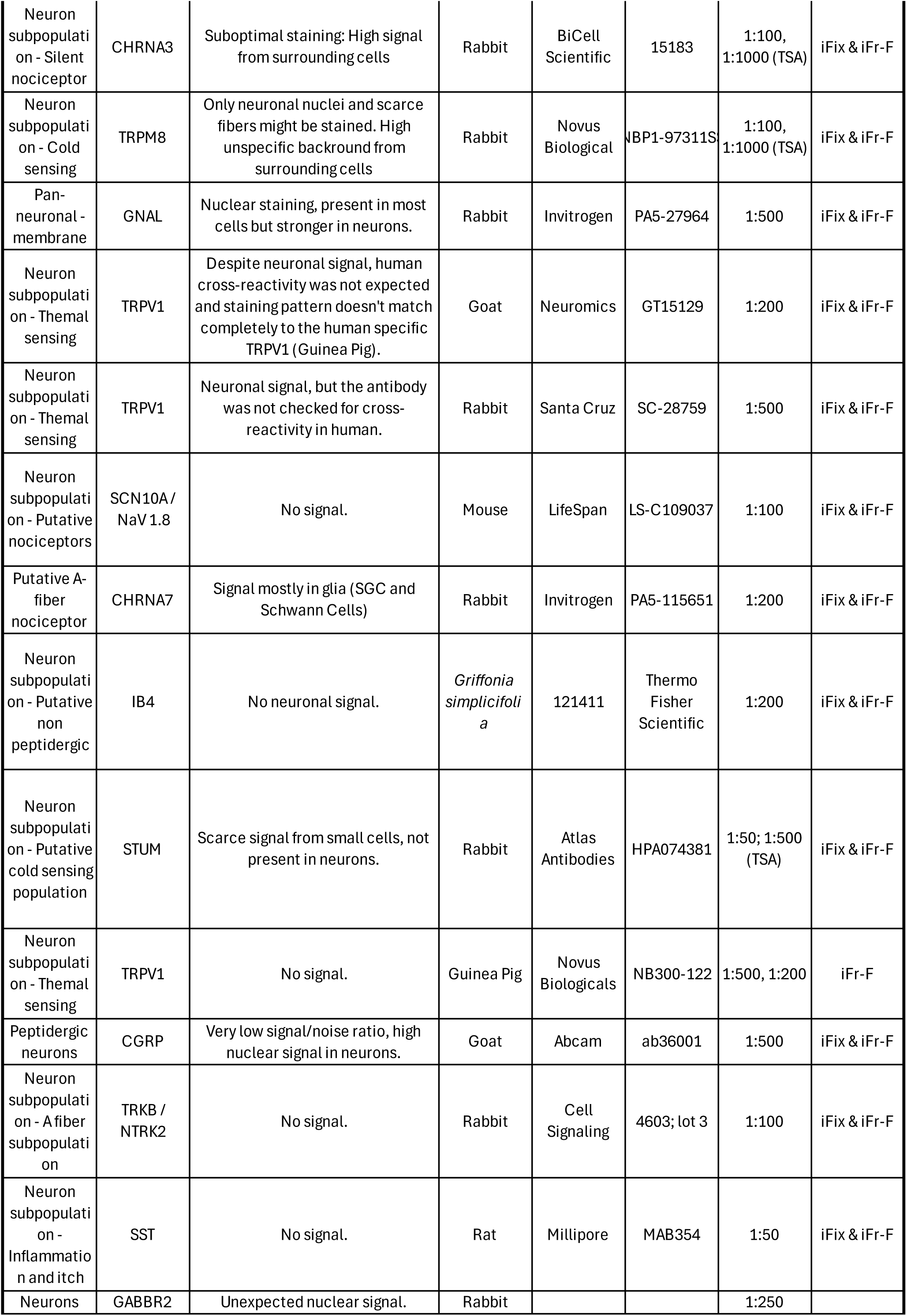
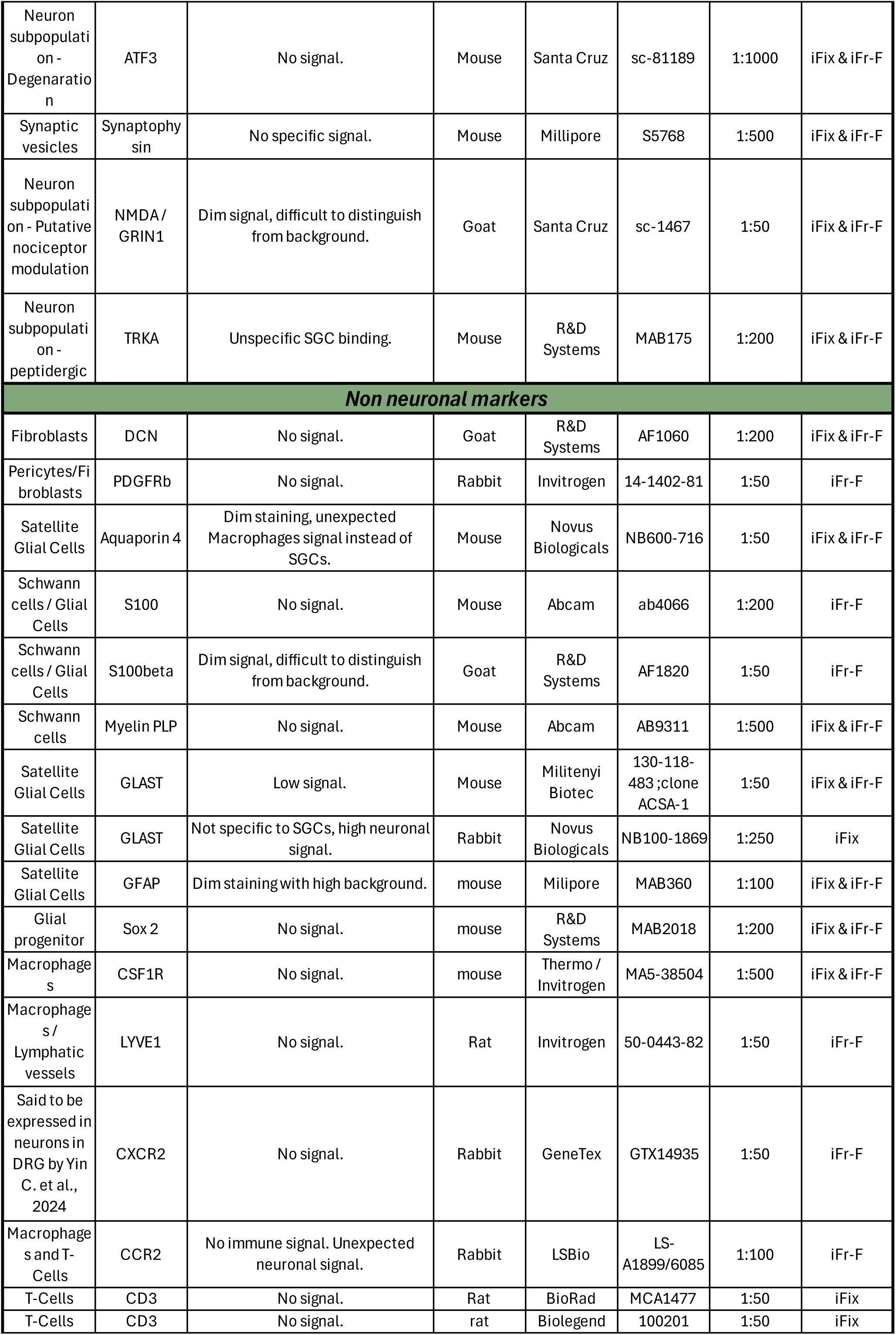
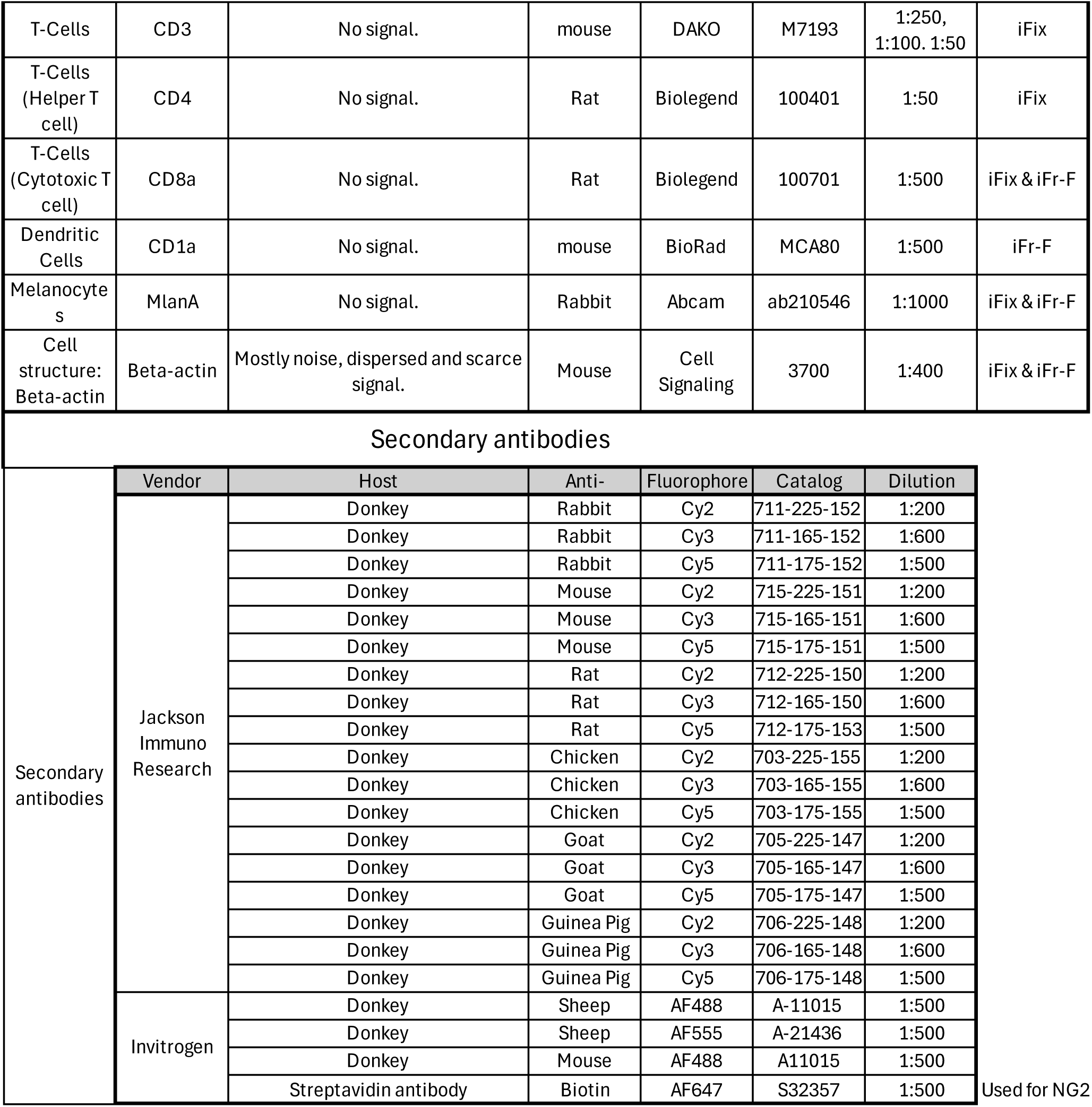

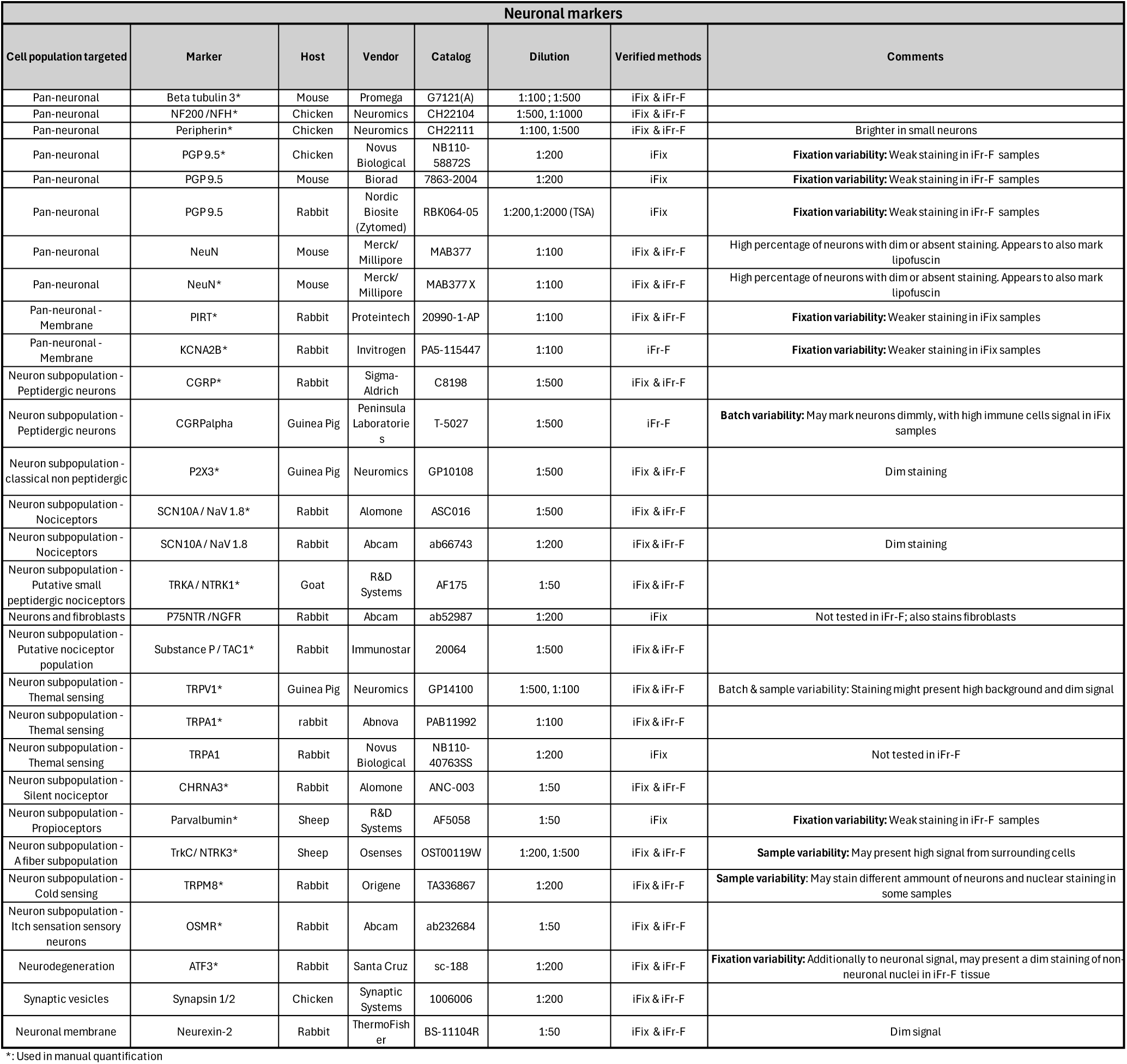

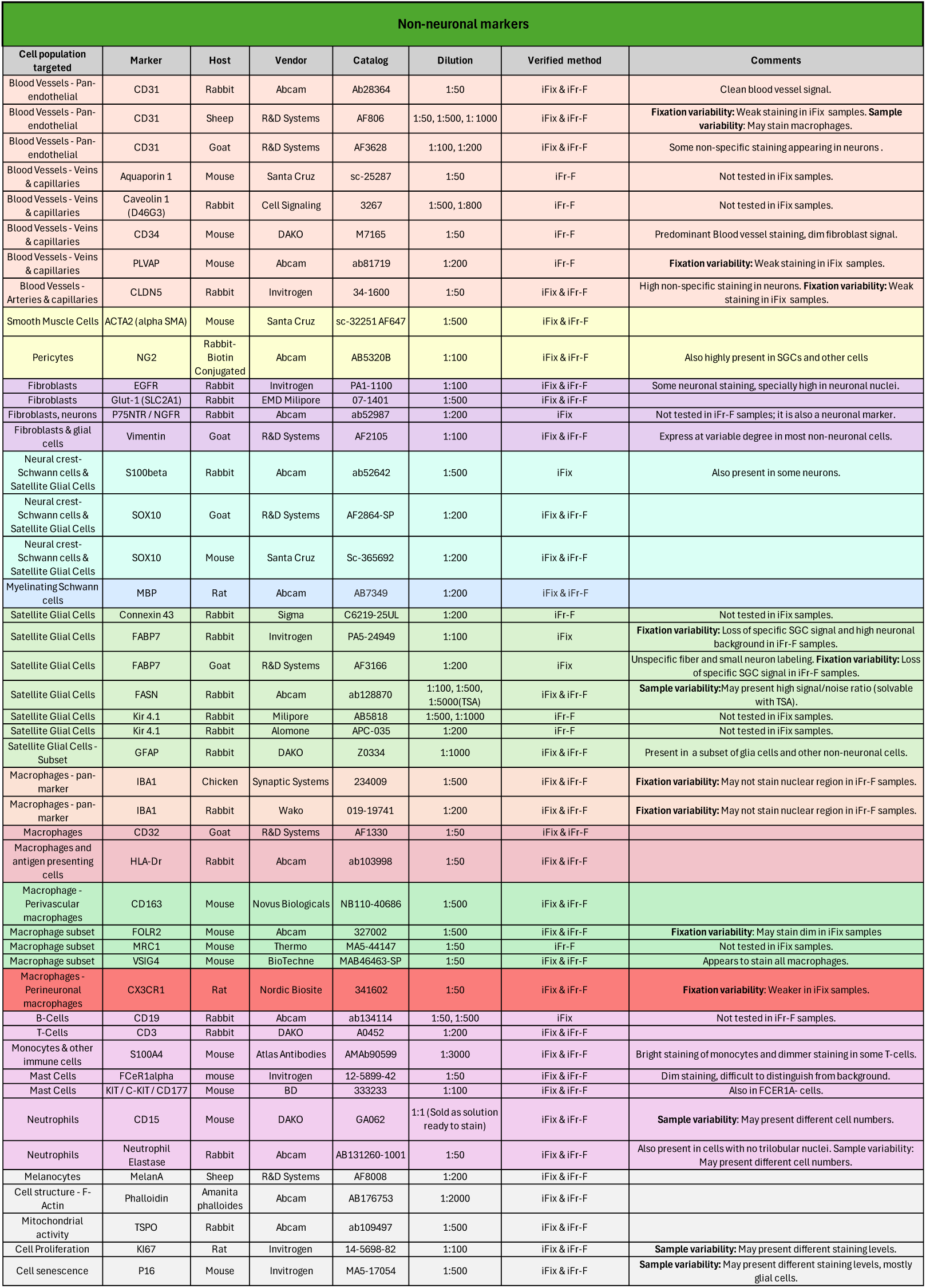

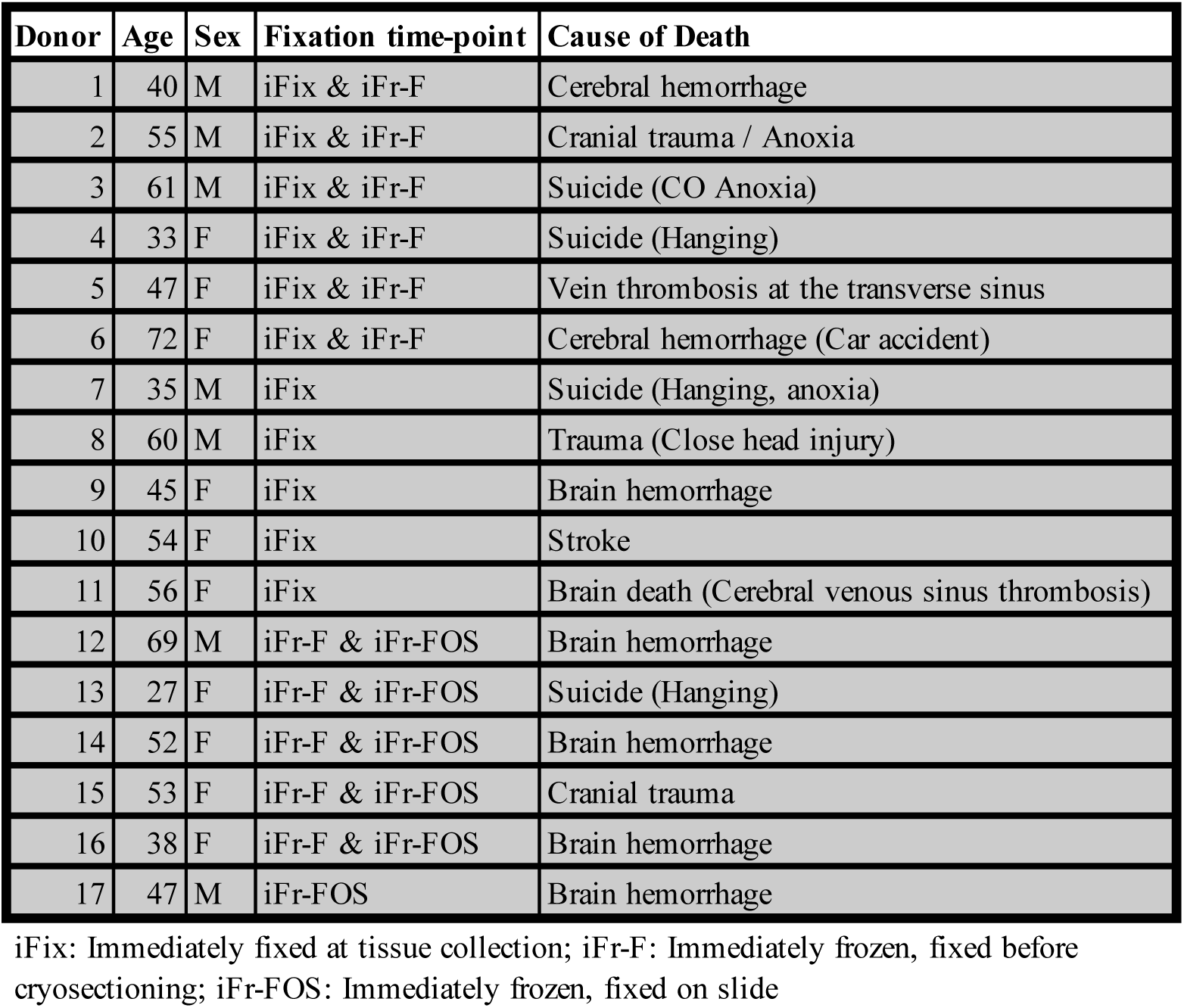

