## Supplementary Table 1 for "Tissue-to-Analysis Framework Enables Multiscale Mapping of the Architectural and Cellular Organization in the Human Dorsal Root Ganglion"

### hDRG IHC in prior publications

| Cells marked | Marker | Species | Brand | Cat number | Dilution/Conc. | Notes on tissue | Notes on IHC | First author | Publication year | DOI |
|---|---|---|---|---|---|---|---|---|---|---|
| **Neuronal Markers** | | | | | | | | | | |
| Neurons - Pan-Neuronal | brain-specific homeobox/POU domain protein 3A | Rabbit | Merck | | 1:200 | 3-8 h postmortem, non-diseased donors | 20 µm cuts; Blocking 10% goat; 1ºAb ONight 4ºC; 2º 3hs room temp. | Rostock, C. | 2018 | https://doi.org/10.1016/j.neuroscience.2017.11.047 |
| Neurons - Pan-Neuronal | NEFL | Rabbit | Cell Signaling Technology | 10768 | | Obtained from organ donors | Embedded, cut at 7µm, and fixed on slide with 4%PFA for 30min at 4°C. Blocked with 3%BSA for 45 min. Primary antibodies overnight at room temperature. | Boyer, K. | 2026 | https://doi.org/10.1101/2025.09.21.677655 |
| Neurons - Pan-Neuronal | NF200 | Rabbit | | | 1:200 | 3 Post-mortem DRG (24-36 hrs Post Mortem Interval) | Slice longitudinally, fixed in Zamboni's fixative at 4°C for 4 hrs.30% Sucrose and cut at 10 µm.48 hr Primary incubation at 4°C | Holford, L.C. | 1994 | 10.1007/BF01262058 |
| Neurons - Pan-Neuronal | NF200 | | | | | Avulsed DRG and post mortem sections | Snap frozen in liquid nitrogen, Fixed on Slide in 4%PFA. Treated with 0.3% hydrogen peroxidel.8 µm. ON primary incubation. Nickel enhanced for antibody reveal. | Anand, U. | 2006 | https://doi.org/10.1016/j.neulet.2006.01.046 |
| Neurons - Pan-Neuronal | NF200 | Chicken | Millipore | | 1:25000 | 3-8 h postmortem, non-diseased donors | 20 µm cuts; Blocking 10% goat; 1ºAb ONight 4ºC; 2º 3hs room temp. | Rostock, C. | 2018 | https://doi.org/10.1016/j.neuroscience.2017.11.047 |
| Neurons - Pan-Neuronal | NF200 | Mouse | Sigma | | 1:600 | 3-8 h postmortem, non-diseased donors | 20 µm cuts; Blocking 10% goat; 1ºAb ONight 4ºC; 2º 3hs room temp. | Rostock, C. | 2018 | https://doi.org/10.1016/j.neuroscience.2017.11.047 |
| Neurons - Pan-Neuronal | NF200 | Mouse | Sigma | | 1:1000 | 15 donors, 24-72h postmortem, healthy donors | 12 µm cuts; 1 Ab overnight 4ºC; secondary antibodies 2h rt | Chang, W. | 2018 | https://doi.org/10.1007/s12264-017-0132-3 |
| Neurons - Pan-Neuronal | NF200 | Chicken | Neuromics | CH22104 | 1:500 | 1 hDRG from a 53 year-old female without chronic pain; | frozen at the time of collection and stored at −80°C . 14 µm thick cuts; fixed 30 min in PFA 4%; blocked with 3%Goat serum, hIgG overnight, other primaries for 2 h | Goebel, A. | 2021 | https://doi.org/10.1172/jci144201 |
| Neurons - Pan-Neuronal | NF200 | Chicken | Millipore | AB5539 | 1:200/0.5mg/mL | 13 patients with brachial plexus injury 3-8months after injury; 7 controls post-mortem during routine autopsy with potential CNS injury | Transfered to ice-cold 4%PFA/PBS fixed ON 4°C, washed 30min PBSand ON 30%Sucrose 4°C. Longitudinally cut in 2, OCT embedded & stored at -80°C. 20 µm cuts. Slides incubated in quenching solution 15 min | Schulte, A. | 2023 | https://doi.org/10.1101/2023.02.06.526934 |
| Neurons - Pan-Neuronal | NF200 | Chicken | BioTechne | NB300-217 | | Obtained from organ donors | Embedded, cut at 7µm, and fixed on slide with 4%PFA for 30min at 4°C. Blocked with 3%BSA for 45 min. Primary antibodies overnight at room temperature. | Boyer, K. | 2026 | https://doi.org/10.1101/2025.09.21.677655 |
| Neurons - Pan-Neuronal | Nissl | | Thermo | N21483 | 1:100 | Healthy and diabetic donors. Fresh human DRG L2-L5 | Fixed upon delivery with 4%PFA and immersed in 30% sucrose for at least 3 night at 4°C. Blocking with 5%BSA for 1h. Nissl incubated with secondary antibodies for 1 hour. | Xu, J. | 2026 | https://doi.org/10.1038/s41586-025-09896-x |
| Neurons - Pan-Neuronal | PGP9.5 | Mouse | Abcam | AB86808 | 1:100 | 3xT11+3xT12 DRGs; | 35 µm cuts; 1 Ab overnight 4ºC; | Desormeaux, C. | 2018 | https://doi.org/10.1097/j.pain.0000000000001208 |
| Neurons - Pan-Neuronal | Tubulin β3 | Mouse | Covance | | 1:750 | 3-8 h postmortem, non-diseased donors | 20 µm cuts; Blocking 10% goat; 1ºAb ONight 4ºC; 2º 3hs room temp. | Rostock, C. | 2018 | https://doi.org/10.1016/j.neuroscience.2017.11.047 |
| Neurons - Pan-Neuronal | Tubulin β3 | | Proteintech | | 1:500 | 5 donors | 20 µm cuts; | Nguyen, M. Q. | 2021 | https://doi.org/10.7554/eLife.71752 |
| Neurons - Pan-Neuronal | Tubulin β3 | | BioLegend | #802001 | | 4 donors for IHC studies | 40 µm cuts; Blocking 10%; primary antibodies overnight; secondary antibodies 1h | Avraham, O. | 2022 | https://doi.org/10.1371/journal.pone.0128951 |
| Neurons - Pan-Neuronal | Tubulin β3 | Mouse | Sigma | T8578 | 1:1000 | Surgically extracted from organ donors within 4 hours of cross-clamp and frozen in dry ice, stored in -80°C. | 12µm cuts. Rehydrated in PBST after thawing and fixed on slide with 10% formaldehyde RT for 15min. Blocked in 4%NDS in PBST for 1 hour, primary antibodies incubated overnight at 4°C. | Ahlgreen, O.A. | 2025 | https://doi.org/10.1002/advs.202511569 |
| Neurons - Pan-Neuronal | Tubulin β3 | Mouse | Thermo | MA1-19187 | | Adult human tissues from organ transplant donors. Thoracic DRG washed with cold PBS and immediately fixed with 4% PFA. | Two protocols: Paraffin-embedded cut at 4 µm, followed TSA Kit protocol with horse radish peroxidase secondary antibodies, after counterstaining repairing with antigen repair solution at 100°C for 10 min \|Cryoprotected in 30% sucrose for 24 hours, OCT embedded and cut at 50µm. Free floating sections blocked with 1%BSA, primary antibodies incubated overnight at 4°C. | Wu, Z. | 2025 | https://doi.org/10.1016/j.cell.2025.02.007 |
| Neurons - Pan-Neuronal (Bright in small) | Peripherin | Rabbit | | | 1:200 | 3 Post-mortem DRG (24-36 hrs Post Mortem Interval) | Slice longitudinally, fixed in Zamboni's fixative at 4°C for 4 hrs.30% Sucrose and cut at 10 µm.48 hr Primary incubation at 4°C | Holford, L.C. | 1994 | 10.1007/BF01262058 |
| Neurons - Pan-Neuronal (Bright in small) | Peripherin | | | | | Avulsed DRG | Snap frozen in liquid nitrogen, Fixed on Slide in 4%PFA. Treated with 0.3% hydrogen peroxidel.8 µm. ON primary incubation. Nickel enhanced for antibody reveal. | Anand, U. | 2006 | https://doi.org/10.1016/j.neulet.2006.01.046 |
| Neurons - Pan-Neuronal (Bright in small) | Peripherin | Rabbit | Sigma | | 1:500 | 15 donors, 24-72h postmortem, healthy donors | 12 µm cuts; 1 Ab overnight 4ºC; secondary antibodies 2h rt | Chang, W. | 2018 | https://doi.org/10.1007/s12264-017-0132-3 |
| Neurons - Pan-Neuronal (Bright in small) | Peripherin | Chicken | Encor | CPCA-Peri | 1:1000 | Tissues collected within 4 hours of cross-clamp | Frozen at the time of collection and stored at -80ºC. 10%Formalin fixation; 10%Goat Serum 1hr; antibody incubation ON 4ºC; Sudan Black as post-treatment | Shiers, S.I. | 2020 | https://doi.org/10.1002/cne.25122 |
| Neurons - Pan-Neuronal (Bright in small) | Peripherin | Chicken | Encorbio | CPCA-Peri | 1:500 | Surgically extracted from organ donors within 4 hours of cross-clamp and frozen in dry ice, stored in -80°C. | 12µm cuts. Rehydrated in PBST after thawing and fixed on slide with 10% formaldehyde RT for 15min. Blocked in 4%NDS in PBST for 1 hour, primary antibodies incubated overnight at 4°C. | Ahlgreen, O.A. | 2025 | https://doi.org/10.1002/advs.202511569 |
| Neurons - Pan-Neuronal (Bright in small) | Peripherin | Chicken | Encor Biotechnologies | CPCA-Peri | 1:1000 | Broad donor screening (90 donors)- Four groups of donors based on diabetes and pain conditions. | Frozen in dry ice, kept at -80°C. Gradually embedded in OCT. 20 cuts and fixed on slide with 10% formalin. Dehydrated. Blocked with 10%NDS with Triton-X 100, and incubated with primary antibodies overnight at 4°C | Shiers, S.I. | 2025 | https://doi.org/10.1038/s41467-025-59538-z |
| Neurons - Pan-Neuronal (Bright in small) | Peripherin | Rabbit | Abcam | ab269861 | | Obtained from organ donors | Embedded, cut at 7µm, and fixed on slide with 4%PFA for 30min at 4°C. Blocked with 3%BSA for 45 min. Primary antibodies overnight at room temperature. | Boyer, K. | 2026 | https://doi.org/10.1101/2025.09.21.677655 |
| Neurons - Nociceptive | NaV 1.8 | | | | | Avulsed DRG | Snap frozen in liquid nitrogen, Fixed on Slide in 4%PFA. Treated with 0.3% hydrogen peroxidel.8 µm. ON primary incubation. Nickel enhanced for antibody reveal. | Anand, U. | 2006 | https://doi.org/10.1016/j.neulet.2006.01.046 |
| Neurons - Non peptidergic | IB4 | Griffonia Simplicifolia | Vector Laboratories | | 2.5µg/mL | children with obstetric brachial plexus lesions | 12um cuts, TSA, horse radish peroxidase secondary antibodies | Shi, T. J. | 2008 | https://doi.org/10.1073/pnas.0810899105 |
| Neurons - Non peptidergic | P2X3R | Mouse | Santa Cruz | B-5 | 5 µg/mL | Tissues collected within 4 hours of cross-clamp | Frozen at the time of collection and stored at -80ºC. 10%Formalin fixation; 10%Goat Serum 1hr; antibody incubation ON 4ºC; Sudan Black as post-treatment | Shiers, S.I. | 2020 | https://doi.org/10.1002/cne.25122 |
| Neurons - Peptidergic | CGRP | | | | - | children with obstetric brachial plexus lesions | 12um cuts, TSA, horse radish peroxidase secondary antibodies | Shi, T. J. | 2008 | https://doi.org/10.1073/pnas.0810899105 |
| Neurons - Peptidergic | CGRP | Chicken | GenTex | | 1:500 | adult human individuals autopsies | 30 µm cuts, incubated free floating for 36 h at 4 °C primary antibodies; 90 min secondary antibodies | Patil, J. | 2010 | https://doi.org/10.1016/j.regpep.2010.03.004 |
| Neurons - Peptidergic | CGRP | Mouse | Abcam | | 1:1000 | 6 DRGs from 3 patients; each went through surgery where the spinal nerve root was to be sacrificed | Immediately stored 4ºC and then processed. 8 µm longitudinal cuts, overnight 4ºC both 1 & 2 Ab | Li, Y. | 2018 | https://doi.org/10.1523/JNEUROSCI.0899-17.2017 |
| Neurons - Peptidergic | CGRP | Rabbit | Immunostar | 24112 | 1:1000 | Tissues collected within 4 hours of cross-clamp | Frozen at the time of collection and stored at -80ºC. 10%Formalin fixation; 10%Goat Serum 1hr; antibody incubation ON 4ºC; Sudan Black as post-treatment | Shiers, S.I. | 2020 | https://doi.org/10.1002/cne.25122 |
| Neurons - Peptidergic subset | Substance P | Rat | Seralab | | 1:50 | 3 Post-mortem DRG (24-36 hrs Post Mortem Interval) | Slice longitudinally, fixed in Zamboni's fixative at 4°C for 4 hrs.30% Sucrose and cut at 10 µm.48 hr Primary incubation at 4°C | Holford, L.C. | 1994 | 10.1007/BF01262058 |
| Neurons - Peptidergic subset | Substance P | Mouse | Novus Biologicals | SP-DE4- 21 | 0.5 µg/mL | adult human individuals autopsies | 30 µm cuts, incubated free floating for 36 h at 4 °C primary antibodies; 90 min secondary antibodies | Patil, J. | 2010 | https://doi.org/10.1016/j.regpep.2010.03.004 |
| Neurons - Synaptic vesicles | Synaptophysin | Rabbit | Epitomics | | 1:1200 | adult human individuals autopsies | 30 µm cuts, incubated free floating for 36 h at 4 °C primary antibodies; 90 min secondary antibodies | Patil, J. | 2010 | https://doi.org/10.1016/j.regpep.2010.03.004 |
| Neurons - Thermal sensing | TRPA1 | Goat | Santa Cruz | C19 sc-32353 | | Avulsion injured DRG (7male, 7 female), and uninjured DRG (8 male) | Snap frozen in liquid nitrogen, Fixed on Slide in 4%PFA. Treated with 0.3% hydrogen peroxide in methanol.10 µm. ON primary incubation. Nickel enhanced ABC for antibody reveal. | Anand, U. | 2008 | https://doi.org/10.1016/j.neulet.2008.04.007 |
| Neurons - Thermal sensing | TRPA1 | Rabbit | GSK | 1962 | | Avulsion injured DRG (7male, 7 female), and uninjured DRG (8 male) | Snap frozen in liquid nitrogen, Fixed on Slide in 4%PFA. Treated with 0.3% hydrogen peroxide in methanol.10 µm. ON primary incubation. Nickel enhanced ABC for antibody reveal. | Anand, U. | 2008 | https://doi.org/10.1016/j.neulet.2008.04.007 |
| Neurons - Thermal sensing | TRPV1 | Rabbit | | C-22 | | Cervical avulsed DRGs collected during surgical repair (n=10) and post mortem (<12h Post Mortem Interval) donors (n=6) | Snap frozen in liquid nitrogen, Fixed on Slide in 4%PFA. Treated with 0.3% hydrogen peroxidel.8 µm. ON primary incubation. Nickel enhanced for antibody reveal. | Smith, G.D. | 2002 | https://doi.org/10.1038/nature00894 |
| Neurons - Thermal sensing | TRPV1 | | | | | Avulsed DRG | Snap frozen in liquid nitrogen, Fixed on Slide in 4%PFA. Treated with 0.3% hydrogen peroxidel.8 µm. ON primary incubation. Nickel enhanced for antibody reveal. | Anand, U. | 2006 | https://doi.org/10.1016/j.neulet.2006.01.046 |
| Neurons - Thermal sensing | TRPV1 | - | GSK | C-22 | | Avulsion injured DRG (7male, 7 female), and uninjured DRG (8 male) | Snap frozen in liquid nitrogen, Fixed on Slide in 4%PFA. Treated with 0.3% hydrogen peroxide in methanol.10 µm. ON primary incubation. Nickel enhanced ABC for antibody reveal. | Anand, U. | 2008 | https://doi.org/10.1016/j.neulet.2008.04.007 |
| Neurons - Thermal sensing | TRPV1 | Rabbit | GSK | CC2 | 1:10,000 | control hDRG (n = 5), avulsion injured hDRG (n = 8). | Tissue stored at −70°C until use or immersed in Zamboni's fixative for 2 h and stored in PBS . 15um cuts, post-fixed in 4% PFA 30 min, peroxidase blocked with methanol containing 0.3% w/v hydrogen peroxide for 30 min,primary antibodies overnight, avidin–biotin peroxidase | Anand, U. | 2015 | https://doi.org/10.1186/s12990-015-0038-x |
| Neurons - Thermal sensing | TRPV1 | Rabbit | Neuromics, Edina, MN | | 1:400 | 15 donors, 24-72h postmortem, healthy donors | 12 µm cuts; 1 Ab overnight 4ºC; secondary antibodies 2h rt | Chang, W. | 2018 | https://doi.org/10.1007/s12264-017-0132-3 |
| Neurons - Thermal sensing | TRPV1 | Guinea pig | Abcam | | 1:250 | 6 DRGs from 3 patients; each went through surgery where the spinal nerve root was to be sacrificed | Immediately stored 4ºC and then processed. 8 µm longitudinal cuts, overnight 4ºC both 1 & 2 Ab | Li, Y. | 2018 | https://doi.org/10.1523/JNEUROSCI.0899-17.2017 |
| Neurons - Thermal sensing | TRPV1 | Rabbit | Thermo Fisher Scientific | PA1-748 | 1:500 | Tissues collected within 4 hours of cross-clamp | Frozen at the time of collection and stored at -80ºC. 10%Formalin fixation; 10%Goat Serum 1hr; antibody incubation ON 4ºC; Sudan Black as post-treatment | Shiers, S.I. | 2020 | https://doi.org/10.1002/cne.25122 |
| Neurons - Thermal sensing | TRPV1 | Rabbit | Thermo Fisher Scientific | PA1-748 | 2µg/mL | Broad donor screening (90 donors)- Four groups of donors based on diabetes and pain conditions. | Frozen in dry ice, kept at -80°C. Gradually embedded in OCT. 20 cuts and fixed on slide with 10% formalin. Dehydrated. Blocked with 10%NDS with Triton-X 100, and incubated with primary antibodies overnight at 4°C | Shiers, S.I. | 2025 | https://doi.org/10.1038/s41467-025-59538-z |
| Neurons - Thermal sensing | TRPV3 | Rabbit | | N-63 | | Cervical avulsed DRGs collected during surgical repair (n=10) and post mortem (<12h Post Mortem Interval) donors (n=6) | Snap frozen in liquid nitrogen, Fixed on Slide in 4%PFA. Treated with 0.3% hydrogen peroxidel.8 µm. ON primary incubation. Nickel enhanced for antibody reveal. | Smith, G.D. | 2002 | https://doi.org/10.1038/nature00894 |
| Neurons - Thermal sensing | TRPV3 | | | | | Avulsed DRG | Snap frozen in liquid nitrogen, Fixed on Slide in 4%PFA. Treated with 0.3% hydrogen peroxidel.8 µm. ON primary incubation. Nickel enhanced for antibody reveal. | Anand, U. | 2006 | https://doi.org/10.1016/j.neulet.2006.01.046 |

| | | | | | | | | | | |
|---|---|---|---|---|---|---|---|---|---|---|
| Neurons - small | ROBO2 | Goat | BioTechne | AF3147 | | Obtained from organ donors | Embedded, cut at 7µm, and fixed on slide with 4%PFA for 30min at 4°C. Blocked with 3%BSA for 45 min. Primary antibodies overnight at room temperature. | Boyer, K. | 2026 | https://doi.org/10.1101/2025.09.21.677655 |
| Neuron | Nav 1.7 | rabbit | Alomone Labs | | 1:250 | 6 DRGs from 3 patients; each went through surgery where the spinal nerve root was to be sacrificed | Immediately stored 4ºC and then processed. 8 µm longitudinal cuts, overnight 4ºC both 1 & 2 Ab | Li, Y. | 2018 | https://doi.org/10.1523/JNEUROSCI.0899-17.2017 |
| Neurons | NaV 1.7 | Mouse | NeuroMab | N68/6 | 2µmL | Tissues collected within 4 hours of cross-clamp | Frozen at the time of collection and stored at -80ºC. 10%Formalin fixation; 10%Goat Serum 1hr; antibody incubation ON 4ºC; Sudan Black as post-treatment | Shiers, S.I. | 2020 | https://doi.org/10.1002/cne.25122 |
| Neurons | Nav 1.7 | Mouse | NeuroMab | N68/6 | 2µg/mL | Broad donor screening (90 donors)- Four groups of donors based on diabetes and pain conditions. | Frozen in dry ice, kept at -80°C. Gradually embedded in OCT. 20 cuts and fixed on slide with 10% formalin. Dehydrated. Blocked with 10%NDS with Triton-X 100, and incubated with primary antibodies overnight at 4°C | Shiers, S.I. | 2025 | https://doi.org/10.1038/s41467-025-59538-z |
| Neurons | AngII (Angiotensin II) | Murine | | Purified using MabTrap G II column, Amersham Sciences | 0.3 µg/mL | adult human individuals autopsies | 30 µm cuts; incubated free floating for 36 h at 4 °C primary antibodies; 90 min secondary antibodiess | Patil, J. | 2010 | https://doi.org/10.1016/j.regpep.2010.03.004 |
| Neurons | AngII (Angiotensin II) | Rabbit | BIOSS-USA | bs-0587R | 1:5-1:1,000 | control hDRG (n = 5), avulsion injured hDRG (n = 8) | Tissue stored at −70°C until use or immersed in Zamboni's fixative for 2 h and stored in PBS. 15um cuts, post-fixed in 4% PFA 30 min, peroxidase blocked with methanol containing 0.3% w/v hydrogen peroxide for 30 min,primary antibodies overnight, avidin–biotin peroxidase | Anand, U. | 2015 | https://doi.org/10.1186/s12990-015-0038-x |
| Neurons | AT2R (Angiotensin II receptor ) | Goat | Santa Cruz | sc-48452 | 1:50-1:100 | control hDRG (n = 5), avulsion injured hDRG (n = 8) | Tissue stored at −70°C until use or immersed in Zamboni's fixative for 2 h and stored in PBS. 15um cuts, post-fixed in 4% PFA 30 min, peroxidase blocked with methanol containing 0.3% w/v hydrogen peroxide for 30 min,primary antibodies overnight, avidin–biotin peroxidase | Anand, U. | 2015 | https://doi.org/10.1186/s12990-015-0038-x |
| Neurons | Cathepsin D | Rabbit | Epitomics | | 1:500 | adult human individuals autopsies | 30 µm cuts; incubated free floating for 36 h at 4 °C primary antibodies; 90 min secondary [illegible] | Patil, J. | 2010 | https://doi.org/10.1016/j.regpep.2010.03.004 |
| Neurons | CXCR2 | Rabbit | Genetex | GTX14935 | | Post-mortem | 4%PFA fixation, 30% sucrose for 3 days | Yin, C. | 2024 | https://doi.org/10.1038/s41467-024-47640-7 |
| Neurons | FcγRIII / CD16 | Mouse | BioLegend | | 1:100 | n=4 ;Brain dead subjects after asystole | 14 µm cuts; postfixed with acetone; primary antibodies overnight rt; | Bersellini Farinotti, A. | 2019 | http://doi.org/10.1084/jem.20181657 |
| Neurons | hIK1 | | | | | Avulsed DRG | Snap frozen in liquid nitrogen, Fixed on Slide in 4%PFA. Treated with 0.3% hydrogen peroxidel.8 µm. ON primary incubation. Nickel enhanced for antibody reveal. | Anand, U. | 2006 | https://doi.org/10.1016/j.neulet.2006.01.046 |
| Neurons | MAP2 (Microtubule-associated protein 2) | Mouse | Sigma | M1406 | 1:500/ 0.5mg/mL | 13 patients with brachial plexus injury 3-8months after injury; 7 controls post-mortem during routine autopsy with potential CNS injury | Transfered to ice-cold 4%PFA/PBS fixed ON 4°C, washed 30min PBSand ON 30%Sucrose 4°C. Longitudinally cut in 2, OCT embedded & stored at -80°C. 20 µm cuts. Slides incubated in quenching solution 15 min | Schulte, A. | 2023 | https://doi.org/10.1101/2023.02.06.526934 |
| Neurons | NaN/SNS2 | | Rabbit | K186 | 0.6 µg/mL | Avulsion injured DRG (12male), and post-mortem DRG (2 male, 2 days Post Mortem Interval) | Snap frozen in liquid nitrogen, Fixed on Slide in 4%PFA. Treated with 0.3% hydrogen peroxidel.8 µm. ON primary incubation. Nickel enhanced for antibody reveal. | Coward, K. | 2000 | 10.1016/S0304-3959(99)00251-1 |
| Neurons | NeuN | Rabbit | Millipore | ABN78A4 | 1:100 | n=4 ;Brain dead subjects after asystole | 14 µm cuts; postfixed with acetone; primary antibodies overnight rt; | Bersellini Farinotti, A. | 2019 | http://doi.org/10.1084/jem.20181657 |
| Neurons | NeuN | | Millipore | | 1:1000 | 5 donors | 20 µm cuts; | Nguyen, M. Q. | 2021 | https://doi.org/10.7554/eLife.71752 |
| Neurons | NeuN | Mouse | Sigma Aldrich | MAB377X | 1:100 | Non diabetic and without chronic pain controls and DPN donors (n not specified for IHC); L4, L5 and S1 DRGs were collected under cold ischemic conditions, within 3 h of aorta cross-clamp | Formalin fixed parafin embedded DRG tissue sections, deparafnized and subject to citrate bufered antigen retrieval. | Doty, M. | 2022 | https://doi.org/10.1038/s41598-022-21394-y |
| Neurons | NeuN | | | | | Post-mortem | 4%PFA fixation, 30% sucrose for 3 days | Yin, C. | 2024 | https://doi.org/10.1038/s41467-024-47640-7 |
| Neurons | NeuN | Rabbit | Huabio | # ET1602-12 | | Adult human tissues from organ transplant donors. Thoracic DRG washed with cold PBS and immediately fixed with 4% PFA. | Paraffin-embedded cut at 4 µm, followed TSA Kit protocol with horse radish peroxidase secondary antibodies, after counterstaining repairing with antigen repair solution at 100°C for 10 min. | Wu, Z. | 2025 | https://doi.org/10.1016/j.cell.2025.02.007 |
| Neurons | PLC β3 (Phospholipase C β3) | | | | - | children with obstetric brachial plexus lesions | 12um cuts, TSA, horse radish peroxidase secondary antibodies | Shi, T. J. | 2008 | https://doi.org/10.1073/pnas.0810899105 |
| Neurons | SNS/PN3 | Rabbit | | K104 | | Avulsion injured DRG (12male), and post-mortem DRG (2 male, 2 days Post Mortem Interval) | Snap frozen in liquid nitrogen, Fixed on Slide in 4%PFA. Treated with 0.3% hydrogen peroxidel.8 µm. ON primary incubation. Nickel enhanced for antibody reveal. | Coward, K. | 2000 | 10.1016/S0304-3959(99)00251-1 |
| Neurons | SNS/PN3 | Rabbit | | K107 | 1 µg/mL | Avulsion injured DRG (12male), and post-mortem DRG (2 male, 2 days Post Mortem Interval) | Snap frozen in liquid nitrogen, Fixed on Slide in 4%PFA. Treated with 0.3% hydrogen peroxidel.8 µm. ON primary incubation. Nickel enhanced for antibody reveal. | Coward, K. | 2000 | 10.1016/S0304-3959(99)00251-1 |
| Neurons | SSE4 | Mouse | | | 1:150 | 3 Post-mortem DRG (24-36 hrs Post Mortem Interval) | Slice longitudinally, fixed in Zamboni's fixative at 4°C for 4 hrs.30% Sucrose and cut at 10 µm.48 hr Primary incubation at 4°C | Holford, L.C. | 1994 | 10.1007/BF01262058 |
| Neurons | ATF3 | Rabbit | Novus Biologicals | NBP1-855816 | 1:200 | L4 DRG from two youg and two aged donors | Flash frozen on dry ice and stored at -80°C. 20 µm cuts. Fixed on slide in 10% formalin. IHC after fluorescent in situ hybridization. Primary overnight at 4°C. | Donovan, L.J. | 2025 | https://doi.org/10.1038/s41593-025-01954-x |
| Neurons and other cells | CD44 | Rabbit | Abclonal | A19020 | 2µg/mL | Broad donor screening (90 donors)- Four groups of donors based on diabetes and pain conditions. | Frozen in dry ice, kept at -80°C. Gradually embedded in OCT. 20 cuts and fixed on slide with 10% formalin. Dehydrated. Blocked with 10%NDS with Triton-X 100, and incubated with primary antibodies overnight at 4°C | Shiers, S.I. | 2025 | https://doi.org/10.1038/s41467-025-59538-z |
| Neurons and other cells | PAR1 (Protese-Activated Receptor 1) | Goat | Abcam | AB111976 | 1:100 | 3xT11+3xT12 DRGs | 35 µm cuts; 1 Ab overnight 4ºC; | Desormeaux, C. | 2018 | https://doi.org/10.1097/j.pain.0000000000001208 |
| Neurons and other cells | PAR2 (Protese-Activated Receptor 2) | Mouse | LifeSpan | LSB2321 | 1:100 | 3xT11+3xT12 DRGs | 35 µm cuts; 1 Ab overnight 4ºC; | Desormeaux, C. | 2018 | https://doi.org/10.1097/j.pain.0000000000001208 |
| Neurons and other cells | PAR4 (Protese-Activated Receptor 4) | - | Abcam | AB70400 | 1:100 | 3xT11+3xT12 DRGs | 35 µm cuts; 1 Ab overnight 4ºC; | Desormeaux, C. | 2018 | https://doi.org/10.1097/j.pain.0000000000001208 |
| Sympathetic fibers | TH | Rabbit | Sigma | AB152 | 2µg/mL | Broad donor screening (90 donors)- Four groups of donors based on diabetes and pain conditions. | Frozen in dry ice, kept at -80°C. Gradually embedded in OCT. 20 cuts and fixed on slide with 10% formalin. Dehydrated. Blocked with 10%NDS with Triton-X 100, and incubated with primary antibodies overnight at 4°C | Shiers, S.I. | 2025 | https://doi.org/10.1038/s41467-025-59538-z |
| **Non-neuronal markers** | | | | | | | | | | |
| Endothelial cells | CD31 | Rabbit | Abcam | Ab28364 | 1:50 | 6 donors | 25-40 µm cuts; primary antibodies overnight [illegible] | Lund, H. | 2023 | https://doi.org/10.1084/jem.20230675 |
| Endothelial cells | CD31 | Sheep | R&D Systems | AF806 | 1:50 | 6 donors | 25-40 µm cuts; primary antibodies overnight [illegible] | Lund, H. | 2023 | https://doi.org/10.1084/jem.20230675 |
| Endothelial cells | CD31 | Mouse | Invitrogen | 14-0319-82 | 2µg/mL | Broad donor screening (90 donors)- Four groups of donors based on diabetes and pain conditions. | Frozen in dry ice, kept at -80°C. Gradually embedded in OCT. 20 cuts and fixed on slide with 10% formalin. Dehydrated. Blocked with 10%NDS with Triton-X 100, and incubated with primary antibodies overnight at 4°C | Shiers, S.I. | 2025 | https://doi.org/10.1038/s41467-025-59538-z |
| Endothelial cells - Arteries & Capillaries | CLDN5 | Rabbit | Thermo | 34-1600 | 1:200 | 6 donors | 25-40 µm cuts; primary antibodies overnight [illegible] | Lund, H. | 2023 | https://doi.org/10.1084/jem.20230675 |
| Endothelial cells - Veins & Capillaries | PLVAP | Mouse | Abcam | Ab81719 | 1:200 | 6 donors | 25-40 µm cuts; primary antibodies overnight [illegible] | Lund, H. | 2023 | https://doi.org/10.1084/jem.20230675 |
| Basal lamina | Laminin | Mouse | Sigma | L8271 | 1:1000 | 15 patients with Friedreich ataxia and 12 "normal" controls obtain during autopsy | 6 µm cuts in paraffin, blocking 10% horse/donkey; overnight 4ºC, horse-radish peroxidase-labeled for secondaries or AF-488 or Cy3 | Koeppen, A.H. | 2016 | https://doi.org/10.1186/s40478-016-0288-5 |
| Extra Cellular Matrix | TNR (Tenascin R) | Goat | R&D Systems | AF3865 | 1:50 | Non diabetic and without chronic pain controls and DPN donors (n not specified for IHC); L4, L5 and S1 DRGs were collected under cold ischemic conditions, within 3 h of aorta cross-clamp | Formalin fixed parafin embedded DRG tissue sections, deparafnized and subject to citrate bufered antigen retrieval. | Doty, M. | 2022 | https://doi.org/10.1038/s41598-022-21394-y |
| Connective tissue | COL IV | Rabbit | Abcam | ab6586 | | Obtained from organ donors | Embedded, cut at 7µm, and fixed on slide with 4%PFA for 30min at 4°C. Blocked with 3%BSA for 45 min. Primary antibodies overnight at room temperature. | Boyer, K. | 2026 | https://doi.org/10.1101/2025.09.21.677655 |
| Satellite Glial Cells & Neurons - Study target | FMS IgG (Fibromyalgia Syndrome Immunoglobulin) | Human | Extracted from patients | - | 100µg/mL | 1 hDRG from a 53 year-old female without chronic pain | Frozen at the time of collection and stored at −80°C . 14 µm thick cuts; fixed 30 min in PFA 4%; blocked with 3%Goat serum, hIgG overnight, other primaries for 2 h | Goebel, A. | 2021 | https://doi.org/10.1172/jci144201 |
| Satellite Glial Cells & Schwann Cells | S100 | Mouse | Santa Cruz | sc-53438 | 0.4 µg/mL | 15 patients with Friedreich ataxia and 12 "normal" controls obtain during autopsy | 6 µm cuts in paraffin, blocking 10% horse/donkey; overnight 4ºC, horse-radish peroxidase-labeled for secondaries or AF-488 or Cy3 | Koeppen, A.H. | 2016 | https://doi.org/10.1186/s40478-016-0288-5 |
| Satellite Glial Cells & Schwann Cells | S100 | Mouse | Fisher Scientific | MA126621 | 2 µg/mL | Broad donor screening (90 donors)- Four groups of donors based on diabetes and pain conditions. | Frozen in dry ice, kept at -80°C. Gradually embedded in OCT. 20 cuts and fixed on slide with 10% formalin. Dehydrated. Blocked with 10%NDS with Triton-X 100, and incubated with primary antibodies overnight at 4°C | Shiers, S.I. | 2025 | https://doi.org/10.1038/s41467-025-59538-z |
| Satellite Glial Cells & Schwann Cells | SOX10 | Mouse | Abcam | ab216020 | 2 µg/mL | Broad donor screening (90 donors)- Four groups of donors based on diabetes and pain conditions. | Frozen in dry ice, kept at -80°C. Gradually embedded in OCT. 20 cuts and fixed on slide with 10% formalin. Dehydrated. Blocked with 10%NDS with Triton-X 100, and incubated with primary antibodies overnight at 4°C | Shiers, S.I. | 2025 | https://doi.org/10.1038/s41467-025-59538-z |
| Satellite Glial Cells & Schwann Cells | SOX10 | Goat | BioTechne | AF2864 | | Obtained from organ donors | Embedded, cut at 7µm, and fixed on slide with 4%PFA for 30min at 4°C. Blocked with 3%BSA for 45 min. Primary antibodies overnight at room temperature. | Boyer, K. | 2026 | https://doi.org/10.1101/2025.09.21.677655 |
| Satellite Glial Cells | Apolipoprotein J / clusterin | Goat | Abcam | ab39991 | 1:50/0.5mg/mL | 13 patients with brachial plexus injury 3-8months after injury; 7 controls post-mortem during routine autopsy with potential CNS injury | Transfered to ice-cold 4%PFA/PBS fixed ON 4°C, washed 30min PBSand ON 30%Sucrose 4°C. Longitudinally cut in 2, OCT embedded & stored at -80°C. 20 µm cuts. Slides incubated in quenching solution 15 min | Schulte, A. | 2023 | https://doi.org/10.1101/2023.02.06.526934 |
| Satellite Glial Cells | Connexin 43 | Rabbit | Abcam | ab113470 | 0.75 µg/mL | 15 patients with Friedreich ataxia and 12 "normal" controls obtain during autopsy | 6 µm cuts in paraffin, blocking 10% horse/donkey; overnight 4ºC, horse-radish peroxidase-labeled for secondaries or AF-488 or Cy3 | Koeppen, A.H. | 2016 | https://doi.org/10.1186/s40478-016-0288-5 |
| Satellite Glial Cells | Connexin 43 | Rabbit | Cell Signaling Technology | #3512 | 1:200 | Surgically extracted from organ donors within 4 hours of cross-clamp and frozen in dry ice, stored in -80°C. | 12µm cuts. Rehydrated in PBST after thawing and fixed on slide with 10% formaldehyde RT for 15min. Blocked in 4%NDS in PBST for 1 hour, primary antibodies incubated overnight at 4°C. | Ahlgreen, O.A. | 2025 | https://doi.org/10.1002/advs.202511569 |
| Satellite Glial Cells | FABP7 | Rabbit | Thermo Fisher Scientific | #PA5-24949 | | 4 donors for IHC studies | 40 µm cuts; Blocking 10%; primary antibodies overnight; secondary antibodies 1h | Avraham, O. | 2022 | https://doi.org/10.1097/j.pain.0000000000002628 |

| | | | | | | | | | | |
|---|---|---|---|---|---|---|---|---|---|---|
| Satellite Glial Cells | FABP7 | Rabbit | Invitrogen | #PA5-24949 | 1:100/0.29mg/mL | 13 patients with brachial plexus injury 3-8months after injury; 7 controls post-mortem during routine autopsy with potential CNS injury | Transfered to ice-cold 4%PFA/PBS fixed ON 4°C, washed 30min PBSand ON 30%Sucrose 4°C. Longitudinally cut in 2, OCT embedded & stored at -80°C. 20 µm cuts. Slides incubated in quenching solution 15 min | Schulte, A. | 2023 | https://doi.org/10.1101/2023.02.06.526934 |
| Satellite Glial Cells | FABP7 | Goat | R&D Systems | AF3166 | 1:30 | Surgically extracted from organ donors within 4 hours of cross-clamp and frozen in dry ice, stored in -80°C. | 12µm cuts. Rehydrated in PBST after thawing and fixed on slide with 10% formaldehyde RT for 15min. Blocked in 4%NDS in PBST for 1 hour, primary antibodies incubated overnight at 4°C. | Ahlgreen, O.A. | 2025 | https://doi.org/10.1002/advs.202511569 |
| Satellite Glial Cells | FABP7 | Rabbit | Thermo | PA5-24949 | 1:100 | Healthy and diabetic donors. Fresh human DRG L2-L5 | Fixed upon delivery with 4%PFA and immersed in 30% sucrose for at least 3 night at 4°C. Blocking with 5%BSA for 1h. Primary incubated overnight at 4°C | Xu, J. | 2026 | https://doi.org/10.1038/s41586-025-09896-x |
| Satellite Glial Cells | FASN | Rabbit | Abcam | #ab128870 | | 4 donors for IHC studies | 40 µm cuts, Blocking 10%, primary antibodies overnight; secondary antibodies 1h | Avraham, O. | 2022 | https://doi.org/10.1097/j.pain.0000000000002628 |
| Satellite Glial Cells | FASN | Rabbit | Abcam | Ab128870 | | 6 donors | 25-40 µm cuts, primary antibodies overnight 4ºC, secondary antibodies 2h rt | Lund, H. | 2023 | https://doi.org/10.1084/jem.20230675 |
| Satellite Glial Cells | GLAST1 / EAAT1 | Goat | Santa Cruz | sc-77557 | 0.5 µg/mL | 15 patients with Friedreich ataxia and 12 "normal" controls obtain during autopsy | 6 µm cuts in paraffin, blocking 10% horse/donkey; overnight 4ºC, horse-radish peroxidase-labeled for secondaries or AF-488 or Cy3 | Koeppen, A.H. | 2016 | https://doi.org/10.1186/s40478-016-0288-5 |
| Satellite Glial Cells | GS | mouse | Santa Cruz | sc-74430 | 1 µg/mL | 15 patients with Friedreich ataxia and 12 "normal" controls obtain during autopsy | 6 µm cuts in paraffin, blocking 10% horse/donkey; overnight 4ºC, horse-radish peroxidase-labeled for secondaries or AF-488 or Cy3 | Koeppen, A.H. | 2016 | https://doi.org/10.1186/s40478-016-0288-5 |
| Satellite Glial Cells | GS | | - | - | - | 4 donors for IHC studies | 40 µm cuts, Blocking 10%, primary antibodies overnight; secondary antibodies 1h | Avraham, O. | 2022 | https://doi.org/10.1097/j.pain.0000000000002628 |
| Satellite Glial Cells | GS | Mouse | BD Transduction | 610517 | 1:200/ 0.25mg/mL | 13 patients with brachial plexus injury 3-8months after injury; 7 controls post-mortem during routine autopsy with potential CNS injury | Transfered to ice-cold 4%PFA/PBS fixed ON 4°C, washed 30min PBSand ON 30%Sucrose 4°C. Longitudinally cut in 2, OCT embedded & stored at -80°C. 20 µm cuts. Slides incubated in quenching solution 15 min | Schulte, A. | 2023 | https://doi.org/10.1101/2023.02.06.526934 |
| Satellite Glial Cells | GS | Mouse | Genetex | GTX630654 | 1:500 | Surgically extracted from organ donors within 4 hours of cross-clamp and frozen in dry ice, stored in -80°C. | 12µm cuts. Rehydrated in PBST after thawing and fixed on slide with 10% formaldehyde RT for 15min. Blocked in 4%NDS in PBST for 1 hour, primary antibodies incubated overnight at 4°C. | Ahlgreen, O.A. | 2025 | https://doi.org/10.1002/advs.202511569 |
| Satellite Glial Cells | GS | Rabbit | Thermo | 11037-2-AP | 1:200 | Surgically extracted from organ donors within 4 hours of cross-clamp and frozen in dry ice, stored in -80°C. | 12µm cuts. Rehydrated in PBST after thawing and fixed on slide with 10% formaldehyde RT for 15min. Blocked in 4%NDS in PBST for 1 hour, primary antibodies incubated overnight at 4°C. | Ahlgreen, O.A. | 2025 | https://doi.org/10.1002/advs.202511569 |
| Satellite Glial Cells | GS | Rabbit | Abcam | ab73593 | | Adult human tissues from organ transplant donors. Thoracic DRG washed with cold PBS and immediately fixed with 4% PFA. | Paraffin-embedded cut at 4 µm, followed TSA Kit protocol with horse radish peroxidase secondary antibodies, after counterstaining repairing with antigen repair solution at 100°C for 10 min. | Wu, Z. | 2025 | https://doi.org/10.1016/j.cell.2025.02.007 |
| Satellite Glial Cells | Kir 3.1 | | | | | 4 donors for IHC studies | 40 µm cuts, Blocking 10%, primary antibodies overnight; secondary antibodies 1h | Avraham, O. | 2022 | https://doi.org/10.1097/j.pain.0000000000002628 |
| Satellite Glial Cells | Kir 4.1 | Rabbit | Alomone | APC-035 | 4 µg/mL | 15 patients with Friedreich ataxia and 12 "normal" controls obtain during autopsy | 6 µm cuts in paraffin, blocking 10% horse/donkey; overnight 4ºC, horse-radish peroxidase-labeled for secondaries or AF-488 or Cy3 | Koeppen, A.H. | 2016 | https://doi.org/10.1186/s40478-016-0288-5 |
| Satellite Glial Cells | Kir 4.1 | Rabbit | Alomone | APC-035 | 1:500 | Surgically extracted from organ donors within 4 hours of cross-clamp and frozen in dry ice, stored in -80°C. | 12µm cuts. Rehydrated in PBST after thawing and fixed on slide with 10% formaldehyde RT for 15min. Blocked in 4%NDS in PBST for 1 hour, primary antibodies incubated overnight at 4°C. | Ahlgreen, O.A. | 2025 | https://doi.org/10.1002/advs.202511569 |
| Satellite Glial Cells | mGluR2/3 (Metabotropic glutamate receptor 2/3) | Rabbit | Novus | NBPI-00924 | 1 µg/mL | 15 patients with Friedreich ataxia and 12 "normal" controls obtain during autopsy | 6 µm cuts in paraffin, blocking 10% horse/donkey; overnight 4ºC, horse-radish peroxidase-labeled for secondaries or AF-488 or Cy3 | Koeppen, A.H. | 2016 | https://doi.org/10.1186/s40478-016-0288-5 |
| Satellite Glial Cells | OR6B2 (Olfatory Receptor 6B2) | Rabbit | Novus Biologicals | NBP1-71360 | 1:30 | Commercially available human DRG | Paraffin tissue sections with a thickness of 5 µm (Zyagen, San Diego, CA, USA) were deparaffinized . Primaries overnight 4ºC, Secondaries 45 min | Flegel, C. | 2015 | https://doi.org/10.1371/journal.pone.0128951 |
| Satellite Glial Cells | S100α | Rabbit | Santa Cruz | SC-7489-R | 0.4 µg/mL | 15 patients with Friedreich ataxia and 12 "normal" controls obtain during autopsy | 6 µm cuts in paraffin, blocking 10% horse/donkey; overnight 4ºC, horse-radish peroxidase-labeled for secondaries or AF-488 or Cy3 | Koeppen, A.H. | 2016 | https://doi.org/10.1186/s40478-016-0288-5 |
| Satellite Glial Cells | Tissue Inhibitor of Metalloprotease 3 (TIMP3) | Mouse | R&D Systems | MAB973 | 1:500 | Deidentified biospecimens | Fixed with 4% PFA 2h, cryoprotected 30% Sucrose. 12 µm thick cuts, primaries incubated Overnight 4C, 2aries 1h RT | Tonello, R. | 2023 | https://doi.org/10.1016/j.bbi.2023.08.005 |
| Satellite Glial Cells - Subset | GFAP | Mouse | Cell Signaling Technology | | 1:1000 | 6 DRGs from 3 patients; each went through surgery where the spinal nerve root was to be sacrificed | Immediately stored 4ºC and then processed. 8 µm longitudinal cuts, overnight 4ºC both 1 & 2 Ab | Li, Y. | 2018 | https://doi.org/10.1523/JNEUROSCI.0899-17.2017 |
| Satellite Glial Cells - Subset | GFAP | Mouse | Millipore | Mab360 | 1:500 | 1 hDRG from a 53 year-old female without chronic pain | Frozen at the time of collection and stored at –80°C . 14 µm thick cuts; fixed 30 min in PFA 4%; blocked with 3%Goat serum, hIgG overnight, other primaries for 2 h | Goebel, A. | 2021 | https://doi.org/10.1172/jci144201 |
| Satellite Glial Cells - Subset | GFAP | | | | | 4 donors for IHC studies | 40 µm cuts, Blocking 10%, primary antibodies overnight; secondary antibodies 1h | Avraham, O. | 2022 | https://doi.org/10.1097/j.pain.0000000000002628 |
| Satellite Glial Cells - Subset | GFAP | | Addgene | 1934450 | | Obtained from organ donors | Embedded, cut at 7µm, and fixed on slide with 4%PFA for 30min at 4°C. Blocked with 3%BSA for 45 min. Primary antibodies overnight at room temperature. | Boyer, K. | 2026 | https://doi.org/10.1101/2025.09.21.677655 |
| Satellite Glial Cells - Subset | GFAP | Rabbit | Origene | DP-014 | 1:100 | 13 patients with brachial plexus injury 3-8months after injury; 7 controls post-mortem during routine autopsy with potential CNS injury | Transfered to ice-cold 4%PFA/PBS fixed ON 4°C, washed 30min PBSand ON 30%Sucrose 4°C. Longitudinally cut in 2, OCT embedded & stored at -80°C. 20 µm cuts. Slides incubated in quenching solution 15 min | Schulte, A. | 2023 | https://doi.org/10.1101/2023.02.06.526934 |
| Satellite Glial Cells - Subset | GFAP | Mouse | NeuroMab | N206A/8 | 2µg/mL | Broad donor screening (90 donors)- Four groups of donors based on diabetes and pain conditions. | Frozen in dry ice, kept at -80°C. Gradually embedded in OCT. 20 cuts and fixed on slide with 10% formalin. Dehydrated. Blocked with 10%NDS with Triton-X 100, and incubated with primary antibodies overnight at 4°C | Shiers, S.I. | 2025 | https://doi.org/10.1038/s41467-025-59538-z |
| Satellite Glial Cells - Subset | SCN7A/NaV2.1 | Rabbit | Novus Biologicals | NB100-81029 | 1:200 | Surgically extracted from organ donors within 4 hours of cross-clamp and frozen in dry ice, stored in -80°C. | 12µm cuts. Rehydrated in PBST after thawing and fixed on slide with 10% formaldehyde RT for 15min. Blocked in 4%NDS in PBST for 1 hour, primary antibodies incubated overnight at 4°C. | Ahlgreen, O.A. | 2025 | https://doi.org/10.1002/advs.202511569 |
| Satellite Glial Cells - Subset | SCN7A/NaV2.1 | Rabbit | BioTechne | NBP1-87075 | | Obtained from organ donors | Embedded, cut at 7µm, and fixed on slide with 4%PFA for 30min at 4°C. Blocked with 3%BSA for 45 min. Primary antibodies overnight at room temperature. | Boyer, K. | 2026 | https://doi.org/10.1101/2025.09.21.677655 |
| Schwann Cells - Myelinating | MBP | | Millipore Sigma | AB9348 | | Obtained from organ donors | Embedded, cut at 7µm, and fixed on slide with 4%PFA for 30min at 4°C. Blocked with 3%BSA for 45 min. Primary antibodies overnight at room temperature. | Boyer, K. | 2026 | https://doi.org/10.1101/2025.09.21.677655 |
| Macrophages | CD68 | Mouse | Santa Cruz | sc-20060 | 4 µg/mL | 15 patients with Friedreich ataxia and 12 "normal" controls obtain during autopsy | 6 µm cuts in paraffin, blocking 10% horse/donkey; overnight 4ºC, horse-radish peroxidase-labeled for secondaries or AF-488 or Cy3 | Koeppen, A.H. | 2016 | https://doi.org/10.1186/s40478-016-0288-5 |
| Macrophages | CD68 | Mouse | Abcam | AB783 | 1:500 | L4 DRGs from 5 control and 5 DPN donors | DRGs held at -20C for cutting, sliced in half, and placed in 10% bufered formalin. 6um cuts; r citrate-based (CD20), or tris/EDTA-based (CD3 and CD68) antigen retrieval; immunodetection with a polymer-based reagent | Hall, B.E. | 2022 | https://doi.org/10.1038/s41598-022-08100-8 |
| Macrophages | FcγRI (Fc-gamma receptor 1) / CD64 | Mouse | Serotec | MCA756g | 1:100 | n=4 ;Brain dead subjects after asystole | 14 µm cuts; postfixed with acetone; primary antibodies overnight rt; | Bersellini Farinotti, A. | 2019 | http://doi.org/10.1084/jem.20181657 |
| Macrophages | IBA1 | Rabbit | Sigma | HPA049234 | 1 µg/mL | 15 patients with Friedreich ataxia and 12 "normal" controls obtain during autopsy | 6 µm cuts in paraffin, blocking 10% horse/donkey; overnight 4ºC, horse-radish peroxidase-labeled for secondaries or AF-488 or Cy3 | Koeppen, A.H. | 2016 | https://doi.org/10.1186/s40478-016-0288-5 |
| Macrophages | IBA1 | Chicken | Synaptic Systems | 234 006 | 1:50 | 6 donors | 25-40 µm cuts; primary antibodies overnight 4ºC, secondary antibodies 2h rt | Lund, H. | 2023 | https://doi.org/10.1084/jem.20230675 |
| Macrophages | IBA1 | Rabbit | FujiFilm Wako | 019-19741 | 1:100/0.5mg/mL | 13 patients with brachial plexus injury 3-8months after injury; 7 controls post-mortem during routine autopsy with potential CNS injury | Transfered to ice-cold 4%PFA/PBS fixed ON 4°C, washed 30min PBSand ON 30%Sucrose 4°C. Longitudinally cut in 2, OCT embedded & stored at -80°C. 20 µm cuts. Slides incubated in quenching solution 15 min | Schulte, A. | 2023 | https://doi.org/10.1101/2023.02.06.526934 |
| Macrophages | IBA1 | Rabbit | Novus Biologicals | NBP2-19019 | | Adult human tissues from organ transplant donors. Thoracic DRG washed with cold PBS and immediately fixed with 4% PFA. | Paraffin-embedded cut at 4 µm, followed TSA Kit protocol with horse radish peroxidase secondary antibodies, after counterstaining repairing with antigen repair solution at 100°C for 10 min. | Wu, Z. | 2025 | https://doi.org/10.1016/j.cell.2025.02.007 |
| Macrophage subset | CD163 | Mouse | Novus Biologicals | NB110-40686 | 1:500 | 6 donors | 25-40 µm cuts; primary antibodies overnight 4ºC, secondary antibodies 2h rt | Lund, H. | 2023 | https://doi.org/10.1084/jem.20230675 |
| Macrophage subset | MRC1 | Mouse | Thermo | MA5-44147 | 1:50 | 6 donors | 25-40 µm cuts; primary antibodies overnight 4ºC, secondary antibodies 2h rt | Lund, H. | 2023 | https://doi.org/10.1084/jem.20230675 |
| Macrophage subset | MRC1 | Rabbit | Cell Signaling Technology | # 24595 | | Adult human tissues from organ transplant donors. Thoracic DRG washed with cold PBS and immediately fixed with 4% PFA. | Paraffin-embedded cut at 4 µm, followed TSA Kit protocol with horse radish peroxidase secondary antibodies, after counterstaining repairing with antigen repair solution at 100°C for 10 min. | Wu, Z. | 2025 | https://doi.org/10.1016/j.cell.2025.02.007 |
| Macrophage subset | CX3CR1 | Rat | BioSite | D070-3 | 1:50 | 6 donors | 25-40 µm cuts; primary antibodies overnight 4ºC, secondary antibodies 2h rt | Lund, H. | 2023 | https://doi.org/10.1084/jem.20230675 |
| Macrophage subset | P2RY12 | Rabbit | Abcam | ab300140 | | Adult human tissues from organ transplant donors. Thoracic DRG washed with cold PBS and immediately fixed with 4% PFA. | Two protocols: Paraffin-embedded cut at 4 µm, followed TSA Kit protocol with horse radish peroxidase secondary antibodies, after counterstaining repairing with antigen repair solution at 100°C for 10 min \|Cryoprotected in 30% sucrose for 24 hours, OCT embedded and cut at 50µm. Free floating sections blocked with 1%BSA, primary antibodies incubated overnight at 4°C. | Wu, Z. | 2025 | https://doi.org/10.1016/j.cell.2025.02.007 |
| B-Cells | CD20 | Mouse | Abcam | AB9475 | 1:100 | L4 DRGs from 5 control and 5 DPN donors | DRGs held at -20C for cutting, sliced in half, and placed in 10% bufered formalin. 6um cuts; r citrate-based (CD20), or tris/EDTA-based (CD3 and CD68) antigen retrieval; immunodetection with a polymer-based reagent | Hall, B.E. | 2022 | https://doi.org/10.1038/s41598-022-08100-8 |
| T-Cells | CD3 | Mouse | Abcam | AB17143 | 1:100 | L4 DRGs from 5 control and 5 DPN donors | DRGs held at -20C for cutting, sliced in half, and placed in 10% bufered formalin. 6um cuts; r citrate-based (CD20), or tris/EDTA-based (CD3 and CD68) antigen retrieval; immunodetection with a polymer-based reagent | Hall, B.E. | 2022 | https://doi.org/10.1038/s41598-022-08100-8 |

| | | | | | | | | | | |
|---|---|---|---|---|---|---|---|---|---|---|
| Mithocondrial | Frataxin | Mouse | Abcam | ab110328 | 10 µg/mL | 15 patients with Friedreich ataxia and 12 "normal" controls obtain during autopsy | 6 µm cuts in paraffin, blocking 10% horse/donkey; overnight 4ºC, horse-radish peroxidase-labeled for secondaries or AF-488 or Cy3 | Koeppen, A.H. | 2016 | https://doi.org/10.1186/s40478-016-0288-5 |
| Initiation of translation | phospho-eIF4E | Rabbit | Abcam | ab76256 | 2µg/mL | Broad donor screening (90 donors)- Four groups of donors based on diabetes and pain conditions. | Frozen in dry ice, kept at -80°C. Gradually embedded in OCT. 20 cuts and fixed on slide with 10% formalin. Dehydrated. Blocked with 10%NDS with Triton-X 100, and incubated with primary antibodies overnight at 4°C | Shiers, S.I. | 2025 | https://doi.org/10.1038/s41467-025-59538-z |
