## Supplementary Table 2 for "Tissue-to-Analysis Framework Enables Multiscale Mapping of the Architectural and Cellular Organization in the Human Dorsal Root Ganglion"

| Cell population targeted | Marker | Comments | Host | Vendor | Catalog | Dilution | Verified methods |
|---|---|---|---|---|---|---|---|
| **Neuronal markers** | | | | | | | |
| Pan-neuronal | Beta tubulin 3 | | Mouse | Promega | G7121(A) | 1:100 ; 1:500 | iFix & iFr-F |
| Pan-neuronal | NF200 /NFH | | Chicken | Neuromics | CH22104 | 1:500, 1:1000 | iFix & iFr-F |
| Pan-neuronal | Peripherin | Brighter in small neurons | Chicken | Neuromics | CH22111 | 1:100, 1:500 | iFix & iFr-F |
| Pan-neuronal | PGP 9.5 | **Fixation variability:** Weak staining in iFr-F samples | Chicken | Novus Biological | NB110-58872S | 1:200 | iFix |
| Pan-neuronal | PGP 9.5 | **Fixation variability:** Weak staining in iFr-F samples | Mouse | Biorad | 7863-2004 | 1:200 | iFix |
| Pan-neuronal | PGP 9.5 | **Fixation variability:** Weak staining in iFr-F samples | Rabbit | Nordic Biosite (Zytomed) | RBK064-05 | 1:200,1:2000 (TSA) | iFix |
| Pan-neuronal | NeuN | High percentage of neurons with dim or absent staining. Appears to also mark lipofuscin | Mouse | Merck/ Millipore | MAB377 | 1:100 | iFix & iFr-F |
| Pan-neuronal | NeuN | High percentage of neurons with dim or absent staining. Appears to also mark lipofuscin | Mouse | Merck/ Millipore | MAB377 X | 1:100 | iFix & iFr-F |
| Pan-neuronal - Membrane | PIRT | **Fixation variability:** Weaker staining in iFix samples | Rabbit | Proteintech | 20990-1-AP | 1:100 | iFix & iFr-F |
| Pan-neuronal - Membrane | KCNA2B | **Fixation variability:** Weaker staining in iFix samples | Rabbit | Invitrogen | PA5-115447 | 1:100 | iFr-F |
| Neuron subpopulation - Peptidergic neurons | CGRP | | Rabbit | Sigma-Aldrich | C8198 | 1:500 | iFix & iFr-F |
| Neuron subpopulation - Peptidergic neurons | CGRPalpha | **Batch variability:** May mark neurons dimmly, with high immune cells signal in iFix samples | Guinea Pig | Peninsula Laboratories | T-5027 | 1:500 | iFr-F |

| | | | | | | | |
|---|---|---|---|---|---|---|---|
| Neuron subpopulation - classical non peptidergic | P2X3 | Dim staining | Guinea Pig | Neuromics | GP10108 | 1:500 | iFix & iFr-F |
| Neuron subpopulation - Nociceptors | SCN10A / NaV 1.8 | | Rabbit | Alomone | ASC016 | 1:500 | iFix & iFr-F |
| Neuron subpopulation - Nociceptors | SCN10A / NaV 1.8 | Dim staining | Rabbit | Abcam | ab66743 | 1:200 | iFix & iFr-F |
| Neuron subpopulation - Putative small peptidergic nociceptors | TRKA / NTRK1 | | Goat | R&D Systems | AF175 | 1:50 | iFix & iFr-F |
| Neurons and fibroblasts | P75NTR /NGFR | Not tested in iFr-F; also stains fibroblasts | Rabbit | Abcam | ab52987 | 1:200 | iFix |
| Neuron subpopulation - Putative nociceptor population | Substance P / TAC1 | | Rabbit | Immunostar | 20064 | 1:500 | iFix & iFr-F |
| Neuron subpopulation - Themal sensing | TRPV1 | Batch & sample variability: Staining might present high background and dim signal | Guinea Pig | Neuromics | GP14100 | 1:500, 1:100 | iFix & iFr-F |
| Neuron subpopulation - Themal sensing | TRPA1 | | rabbit | Abnova | PAB11992 | 1:100 | iFix & iFr-F |
| Neuron subpopulation - Themal sensing | TRPA1 | Not tested in iFr-F | Rabbit | Novus Biological | NB110-40763SS | 1:200 | iFix |
| Neuron subpopulation - Silent nociceptor | CHRNA3 | | Rabbit | Alomone | ANC-003 | 1:50 | iFix & iFr-F |
| Neuron subpopulation - Propioceptors | Parvalbumin | **Fixation variability:** Weak staining in iFr-F samples | Sheep | R&D Systems | AF5058 | 1:50 | iFix |
| Neuron subpopulation - A fiber subpopulation | TrkC/ NTRK3 | **Sample variability:** May present high signal from surrounding cells | Sheep | Osenses | OST00119W | 1:200, 1:500 | iFix & iFr-F |

| | | | | | | | |
|---|---|---|---|---|---|---|---|
| Neuron subpopulation - Cold sensing | TRPM8 | **Sample variability**: May stain different ammount of neurons and nuclear staining in some samples | Rabbit | Origene | TA336867 | 1:200 | iFix & iFr-F |
| Neuron subpopulation - Itch sensation sensory neurons | OSMR | | Rabbit | Abcam | ab232684 | 1:50 | iFix & iFr-F |
| Neurodegeneration | ATF3 | **Fixation variability:** Additionally to neuronal signal, may present a dim staining of non-neuronal nuclei in iFr-F tissue | Rabbit | Santa Cruz | sc-188 | 1:200 | iFix & iFr-F |
| Synaptic vesicles | Synapsin 1/2 | | Chicken | Synaptic Systems | 1006006 | 1:200 | iFix & iFr-F |
| Neuronal membrane | Neurexin-2 | Dim signal | Rabbit | ThermoFisher | BS-11104R | 1:50 | iFix & iFr-F |
| **Non-neuronal markers** | | | | | | | |
| Blood Vessels - Pan-endothelial | CD31 | Clean blood vessel signal. | Rabbit | Abcam | Ab28364 | 1:50 | iFix & iFr-F |
| Blood Vessels - Pan-endothelial | CD31 | **Fixation variability:** Weak staining in iFix samples. **Sample variability**: May stain macrophages. | Sheep | R&D Systems | AF806 | 1:50, 1:500, 1: 1000 | iFix & iFr-F |
| Blood Vessels - Pan-endothelial | CD31 | Some non-specific staining appearing in neurons . | Goat | R&D Systems | AF3628 | 1:100, 1:200 | iFix & iFr-F |
| Blood Vessels - Veins & capillaries | Aquaporin 1 | Not tested in iFix samples. | Mouse | Santa Cruz | sc-25287 | 1:50 | iFr-F |
| Blood Vessels - Veins & capillaries | Caveolin 1 (D46G3) | Not tested in iFix samples. | Rabbit | Cell Signaling | 3267 | 1:500, 1:800 | iFr-F |
| Blood Vessels - Veins & capillaries | CD34 | Predominant Blood vessel staining, dim fibroblast signal. | Mouse | DAKO | M7165 | 1:50 | iFr-F |
| Blood Vessels - Veins & capillaries | PLVAP | **Fixation variability:** Weak staining in iFix samples. | Mouse | Abcam | ab81719 | 1:200 | iFr-F |
| Blood Vessels - Arteries & capillaries | CLDN5 | High non-specific staining in neurons. **Fixation variability:** Weak staining in iFix samples. | Rabbit | Invitrogen | 34-1600 | 1:50 | iFix & iFr-F |
| Smooth Muscle Cells | ACTA2 (alpha SMA) | | Mouse | Santa Cruz | c-32251 AF64 | 1:500 | iFix & iFr-F |
| Pericytes | NG2 | Also highly present in SGCs and other cells | Rabbit-Biotin Conjugated | Abcam | AB5320B | 1:100 | iFix & iFr-F |

| | | | | | | | |
|---|---|---|---|---|---|---|---|
| Fibroblasts | EGFR | Some neuronal staining, specially high in neuronal nuclei. | Rabbit | Invitrogen | PA1-1100 | 1:100 | iFix & iFr-F |
| Fibroblasts | Glut-1 (SLC2A1) | | Rabbit | EMD Milipore | 07-1401 | 1:500 | iFix & iFr-F |
| Fibroblasts, neurons | P75NTR / NGFR | Not tested in iFr-F samples; it is also a neuronal marker. | Rabbit | Abcam | ab52987 | 1:200 | iFix |
| Fibroblasts & glial cells | Vimentin | Express at variable degree in most non-neuronal cells. | Goat | R&D Systems | AF2105 | 1:100 | iFix & iFr-F |
| Neural crest-Schwann cells & Satellite Glial Cells | S100beta | Also present in some neurons. | Rabbit | Abcam | ab52642 | 1:500 | iFix |
| Neural crest-Schwann cells & Satellite Glial Cells | SOX10 | | Goat | R&D Systems | AF2864-SP | 1:200 | iFix & iFr-F |
| Neural crest-Schwann cells & Satellite Glial Cells | SOX10 | | Mouse | Santa Cruz | Sc-365692 | 1:200 | iFix & iFr-F |
| Myelinating Schwann cells | MBP | | Rat | Abcam | AB7349 | 1:200 | iFix & iFr-F |
| Satellite Glial Cells | Connexin 43 | Not tested in iFix samples. | Rabbit | Sigma | C6219-25UL | 1:200 | iFr-F |
| Satellite Glial Cells | FABP7 | **Fixation variability:** Loss of specific SGC signal and high neuronal background in iFr-F samples. | Rabbit | Invitrogen | PA5-24949 | 1:100 | iFix |
| Satellite Glial Cells | FABP7 | Unspecific fiber and small neuron labeling. **Fixation variability:** Loss of specific SGC signal in iFr-F samples. | Goat | R&D Systems | AF3166 | 1:200 | iFix |
| Satellite Glial Cells | FASN | **Sample variability:**May present high signal/noise ratio (solvable with TSA). | Rabbit | Abcam | ab128870 | 1:100, 1:500, 1:5000(TSA) | iFix & iFr-F |
| Satellite Glial Cells | Kir 4.1 | Not tested in iFix samples. | Rabbit | Milipore | AB5818 | 1:500, 1:1000 | iFr-F |
| Satellite Glial Cells | Kir 4.1 | Not tested in iFix samples. | Rabbit | Alomone | APC-035 | 1:200 | iFr-F |
| Satellite Glial Cells - Subset | GFAP | Present in a subset of glia cells and other non-neuronal cells. | Rabbit | DAKO | Z0334 | 1:1000 | iFix & iFr-F |
| Macrophages - pan-marker | IBA1 | **Fixation variability:** May not stain nuclear region in iFr-F samples. | Chicken | Synaptic Systems | 234009 | 1:500 | iFix & iFr-F |
| Macrophages - pan-marker | IBA1 | **Fixation variability:** May not stain nuclear region in iFr-F samples. | Rabbit | Wako | 019-19741 | 1:200 | iFix & iFr-F |
| Macrophages | CD32 | | Goat | R&D Systems | AF1330 | 1:50 | iFix & iFr-F |

| | | | | | | | |
|---|---|---|---|---|---|---|---|
| Macrophages and antigen presenting cells | HLA-Dr | | Rabbit | Abcam | ab103998 | 1:50 | iFix & iFr-F |
| Macrophage - Perivascular macrophages | CD163 | | Mouse | Novus Biologicals | NB110-40686 | 1:500 | iFix & iFr-F |
| Macrophage subset | FOLR2 | **Fixation variability**: May stain dim in iFix samples | Mouse | Abcam | 327002 | 1:500 | iFix & iFr-F |
| Macrophage subset | MRC1 | Not tested in iFix samples. | Mouse | Thermo | MA5-44147 | 1:50 | iFr-F |
| Macrophage subset | VSIG4 | Appears to stain all macrophages. | Mouse | BioTechne | MAB46463-SP | 1:50 | iFix & iFr-F |
| Macrophages - Perineuronal macrophages | CX3CR1 | **Fixation variability**: Weaker in iFix samples. | Rat | Nordic Biosite | 341602 | 1:50 | iFix & iFr-F |
| B-Cells | CD19 | Not tested in iFr-F samples. | Rabbit | Abcam | ab134114 | 1:50, 1:500 | iFix |
| T-Cells | CD3 | | Rabbit | DAKO | A0452 | 1:200 | iFix & iFr-F |
| Monocytes & other immune cells | S100A4 | Bright staining of monocytes and dimmer staining in some T-cells. | Mouse | Atlas Antibodies | AMAb90599 | 1:3000 | iFix & iFr-F |
| Mast Cells | FCeR1alpha | Dim staining, difficult to distinguish from background. | mouse | Invitrogen | 12-5899-42 | 1:50 | iFix & iFr-F |
| Mast Cells | KIT / C-KIT / CD177 | Also in FCER1A- cells. | Mouse | BD | 333233 | 1:100 | iFix & iFr-F |
| Neutrophils | CD15 | **Sample variability**: May present different cell numbers. | Mouse | DAKO | GA062 | 1:1 (Sold as solution ready to stain) | iFix & iFr-F |
| Neutrophils | Neutrophil Elastase | Also present in cells with no trilobular nuclei. Sample variability: May present different cell numbers. | Rabbit | Abcam | B131260-100 | 1:50 | iFix & iFr-F |
| Melanocytes | MelanA | | Sheep | R&D Systems | AF8008 | 1:200 | iFix & iFr-F |
| Cell structure - F-Actin | Phalloidin | | Amanita phalloides | Abcam | AB176753 | 1:2000 | iFix & iFr-F |
| Mitochondrial activity | TSPO | | Rabbit | Abcam | ab109497 | 1:500 | iFix & iFr-F |
| Cell Proliferation | KI67 | **Sample variability:** May present different staining levels. | Rat | Invitrogen | 14-5698-82 | 1:100 | iFix & iFr-F |
| Cell senescence | P16 | **Sample variability:** May present different staining levels, mostly glial cells. | Mouse | Invitrogen | MA5-17054 | 1:500 | iFix & iFr-F |
| **Not Working** | | | | | | | |
| ***Neuronal markers*** | | | | | | | |

| | | | | | | | |
|---|---|---|---|---|---|---|---|
| Neuron subpopulation - Silent nociceptor | CHRNA3 | Suboptimal staining: High signal from surrounding cells | Rabbit | BiCell Scientific | 15183 | 1:100, 1:1000 (TSA) | iFix & iFr-F |
| Neuron subpopulation - Cold sensing | TRPM8 | Only neuronal nuclei and scarce fibers might be stained. High unspecific backround from surrounding cells | Rabbit | Novus Biological | NBP1-97311S | 1:100, 1:1000 (TSA) | iFix & iFr-F |
| Pan-neuronal - membrane | GNAL | Nuclear staining, present in most cells but stronger in neurons. | Rabbit | Invitrogen | PA5-27964 | 1:500 | iFix & iFr-F |
| Neuron subpopulation - Themal sensing | TRPV1 | Despite neuronal signal, human cross-reactivity was not expected and staining pattern doesn't match completely to the human specific TRPV1 (Guinea Pig). | Goat | Neuromics | GT15129 | 1:200 | iFix & iFr-F |
| Neuron subpopulation - Themal sensing | TRPV1 | Neuronal signal, but the antibody was not checked for cross-reactivity in human. | Rabbit | Santa Cruz | SC-28759 | 1:500 | iFix & iFr-F |
| Neuron subpopulation - Putative nociceptors | SCN10A / NaV 1.8 | No signal. | Mouse | LifeSpan | LS-C109037 | 1:100 | iFix & iFr-F |
| Putative A-fiber nociceptor | CHRNA7 | Signal mostly in glia (SGC and Schwann Cells) | Rabbit | Invitrogen | PA5-115651 | 1:200 | iFix & iFr-F |
| Neuron subpopulation - Putative non peptidergic | IB4 | No neuronal signal. | *Griffonia simplicifolia* | 121411 | Thermo Fisher Scientific | 1:200 | iFix & iFr-F |
| Neuron subpopulation - Putative cold sensing population | STUM | Scarce signal from small cells, not present in neurons. | Rabbit | Atlas Antibodies | HPA074381 | 1:50; 1:500 (TSA) | iFix & iFr-F |
| Neuron subpopulation - Themal sensing | TRPV1 | No signal. | Guinea Pig | Novus Biologicals | NB300-122 | 1:500, 1:200 | iFr-F |
| Peptidergic neurons | CGRP | Very low signal/noise ratio, high nuclear signal in neurons. | Goat | Abcam | ab36001 | 1:500 | iFix & iFr-F |
| Neuron subpopulation - A fiber subpopulation | TRKB / NTRK2 | No signal. | Rabbit | Cell Signaling | 4603; lot 3 | 1:100 | iFix & iFr-F |
| Neuron subpopulation - Inflammation and itch | SST | No signal. | Rat | Millipore | MAB354 | 1:50 | iFix & iFr-F |
| Neurons | GABBR2 | Unexpected nuclear signal. | Rabbit | | | 1:250 | |

| | | | | | | | |
|---|---|---|---|---|---|---|---|
| Neuron subpopulation - Degenaration | ATF3 | No signal. | Mouse | Santa Cruz | sc-81189 | 1:1000 | iFix & iFr-F |
| Synaptic vesicles | Synaptophysin | No specific signal. | Mouse | Millipore | S5768 | 1:500 | iFix & iFr-F |
| Neuron subpopulation - Putative nociceptor modulation | NMDA / GRIN1 | Dim signal, difficult to distinguish from background. | Goat | Santa Cruz | sc-1467 | 1:50 | iFix & iFr-F |
| Neuron subpopulation - peptidergic | TRKA | Unspecific SGC binding. | Mouse | R&D Systems | MAB175 | 1:200 | iFix & iFr-F |
| ***Non neuronal markers*** | | | | | | | |
| Fibroblasts | DCN | No signal. | Goat | R&D Systems | AF1060 | 1:200 | iFix & iFr-F |
| Pericytes/Fibroblasts | PDGFRb | No signal. | Rabbit | Invitrogen | 14-1402-81 | 1:50 | iFr-F |
| Satellite Glial Cells | Aquaporin 4 | Dim staining, unexpected Macrophages signal instead of SGCs. | Mouse | Novus Biologicals | NB600-716 | 1:50 | iFix & iFr-F |
| Schwann cells / Glial Cells | S100 | No signal. | Mouse | Abcam | ab4066 | 1:200 | iFr-F |
| Schwann cells / Glial Cells | S100beta | Dim signal, difficult to distinguish from background. | Goat | R&D Systems | AF1820 | 1:50 | iFr-F |
| Schwann cells | Myelin PLP | No signal. | Mouse | Abcam | AB9311 | 1:500 | iFix & iFr-F |
| Satellite Glial Cells | GLAST | Low signal. | Mouse | Militenyi Biotec | 130-118-483 ;clone ACSA-1 | 1:50 | iFix & iFr-F |
| Satellite Glial Cells | GLAST | Not specific to SGCs, high neuronal signal. | Rabbit | Novus Biologicals | NB100-1869 | 1:250 | iFix |
| Satellite Glial Cells | GFAP | Dim staining with high background. | mouse | Milipore | MAB360 | 1:100 | iFix & iFr-F |
| Glial progenitor | Sox 2 | No signal. | mouse | R&D Systems | MAB2018 | 1:200 | iFix & iFr-F |
| Macrophages | CSF1R | No signal. | mouse | Thermo / Invitrogen | MA5-38504 | 1:500 | iFix & iFr-F |
| Macrophages / Lymphatic vessels | LYVE1 | No signal. | Rat | Invitrogen | 50-0443-82 | 1:50 | iFr-F |
| Said to be expressed in neurons in DRG by Yin C. et al., 2024 | CXCR2 | No signal. | Rabbit | GeneTex | GTX14935 | 1:50 | iFr-F |
| Macrophages and T-Cells | CCR2 | No immune signal. Unexpected neuronal signal. | Rabbit | LSBio | LS-A1899/6085 | 1:100 | iFr-F |
| T-Cells | CD3 | No signal. | Rat | BioRad | MCA1477 | 1:50 | iFix |
| T-Cells | CD3 | No signal. | rat | Biolegend | 100201 | 1:50 | iFix |

| | | | | | | | |
|---|---|---|---|---|---|---|---|
| T-Cells | CD3 | No signal. | mouse | DAKO | M7193 | 1:250, 1:100. 1:50 | iFix |
| T-Cells (Helper T cell) | CD4 | No signal. | Rat | Biolegend | 100401 | 1:50 | iFix |
| T-Cells (Cytotoxic T cell) | CD8a | No signal. | Rat | Biolegend | 100701 | 1:500 | iFix & iFr-F |
| Dendritic Cells | CD1a | No signal. | mouse | BioRad | MCA80 | 1:500 | iFr-F |
| Melanocytes | MlanA | No signal. | Rabbit | Abcam | ab210546 | 1:1000 | iFix & iFr-F |
| Cell structure: Beta-actin | Beta-actin | Mostly noise, dispersed and scarce signal. | Mouse | Cell Signaling | 3700 | 1:400 | iFix & iFr-F |

### Secondary antibodies

| | Vendor | Host | Anti- | Fluorophore | Catalog | Dilution | |
|---|---|---|---|---|---|---|---|
| Secondary antibodies | Jackson Immuno Research | Donkey | Rabbit | Cy2 | 711-225-152 | 1:200 | |
| | | Donkey | Rabbit | Cy3 | 711-165-152 | 1:600 | |
| | | Donkey | Rabbit | Cy5 | 711-175-152 | 1:500 | |
| | | Donkey | Mouse | Cy2 | 715-225-151 | 1:200 | |
| | | Donkey | Mouse | Cy3 | 715-165-151 | 1:600 | |
| | | Donkey | Mouse | Cy5 | 715-175-151 | 1:500 | |
| | | Donkey | Rat | Cy2 | 712-225-150 | 1:200 | |
| | | Donkey | Rat | Cy3 | 712-165-150 | 1:600 | |
| | | Donkey | Rat | Cy5 | 712-175-153 | 1:500 | |
| | | Donkey | Chicken | Cy2 | 703-225-155 | 1:200 | |
| | | Donkey | Chicken | Cy3 | 703-165-155 | 1:600 | |
| | | Donkey | Chicken | Cy5 | 703-175-155 | 1:500 | |
| | | Donkey | Goat | Cy2 | 705-225-147 | 1:200 | |
| | | Donkey | Goat | Cy3 | 705-165-147 | 1:600 | |
| | | Donkey | Goat | Cy5 | 705-175-147 | 1:500 | |
| | | Donkey | Guinea Pig | Cy2 | 706-225-148 | 1:200 | |
| | | Donkey | Guinea Pig | Cy3 | 706-165-148 | 1:600 | |
| | | Donkey | Guinea Pig | Cy5 | 706-175-148 | 1:500 | |
| | Invitrogen | Donkey | Sheep | AF488 | A-11015 | 1:500 | |
| | | Donkey | Sheep | AF555 | A-21436 | 1:500 | |
| | | Donkey | Mouse | AF488 | A11015 | 1:500 | |
| | | Streptavidin antibody | Biotin | AF647 | S32357 | 1:500 | Used for NG2 |
