## Supplementary Table 3 for "Tissue-to-Analysis Framework Enables Multiscale Mapping of the Architectural and Cellular Organization in the Human Dorsal Root Ganglion"

| Neuronal markers | | | | | | | |
|---|---|---|---|---|---|---|---|
| **Cell population targeted** | **Marker** | **Host** | **Vendor** | **Catalog** | **Dilution** | **Verified methods** | **Comments** |
| Pan-neuronal | Beta tubulin 3* | Mouse | Promega | G7121(A) | 1:100 ; 1:500 | iFix & iFr-F | |
| Pan-neuronal | NF200 /NFH* | Chicken | Neuromics | CH22104 | 1:500, 1:1000 | iFix & iFr-F | |
| Pan-neuronal | Peripherin* | Chicken | Neuromics | CH22111 | 1:100, 1:500 | iFix & iFr-F | Brighter in small neurons |
| Pan-neuronal | PGP 9.5* | Chicken | Novus Biological | NB110-58872S | 1:200 | iFix | **Fixation variability:** Weak staining in iFr-F samples |
| Pan-neuronal | PGP 9.5 | Mouse | Biorad | 7863-2004 | 1:200 | iFix | **Fixation variability:** Weak staining in iFr-F samples |
| Pan-neuronal | PGP 9.5 | Rabbit | Nordic Biosite (Zytomed) | RBK064-05 | 1:200,1:2000 (TSA) | iFix | **Fixation variability:** Weak staining in iFr-F samples |
| Pan-neuronal | NeuN | Mouse | Merck/ Millipore | MAB377 | 1:100 | iFix & iFr-F | High percentage of neurons with dim or absent staining. Appears to also mark lipofuscin |
| Pan-neuronal | NeuN* | Mouse | Merck/ Millipore | MAB377 X | 1:100 | iFix & iFr-F | High percentage of neurons with dim or absent staining. Appears to also mark lipofuscin |
| Pan-neuronal - Membrane | PIRT* | Rabbit | Proteintech | 20990-1-AP | 1:100 | iFix & iFr-F | **Fixation variability:** Weaker staining in iFix samples |
| Pan-neuronal - Membrane | KCNA2B* | Rabbit | Invitrogen | PA5-115447 | 1:100 | iFr-F | **Fixation variability:** Weaker staining in iFix samples |
| Neuron subpopulation - Peptidergic neurons | CGRP* | Rabbit | Sigma-Aldrich | C8198 | 1:500 | iFix & iFr-F | |
| Neuron subpopulation - Peptidergic neurons | CGRPalpha | Guinea Pig | Peninsula Laboratories | T-5027 | 1:500 | iFr-F | **Batch variability:** May mark neurons dimmly, with high immune cells signal in iFix samples |
| Neuron subpopulation - classical non peptidergic | P2X3* | Guinea Pig | Neuromics | GP10108 | 1:500 | iFix & iFr-F | Dim staining |
| Neuron subpopulation - Nociceptors | SCN10A / NaV 1.8* | Rabbit | Alomone | ASC016 | 1:500 | iFix & iFr-F | |
| Neuron subpopulation - Nociceptors | SCN10A / NaV 1.8 | Rabbit | Abcam | ab66743 | 1:200 | iFix & iFr-F | Dim staining |
| Neuron subpopulation - Putative small peptidergic nociceptors | TRKA / NTRK1* | Goat | R&D Systems | AF175 | 1:50 | iFix & iFr-F | |
| Neurons and fibroblasts | P75NTR /NGFR | Rabbit | Abcam | ab52987 | 1:200 | iFix | Not tested in iFr-F; also stains fibroblasts |
| Neuron subpopulation - Putative nociceptor population | Substance P / TAC1* | Rabbit | Immunostar | 20064 | 1:500 | iFix & iFr-F | |
| Neuron subpopulation - Themal sensing | TRPV1* | Guinea Pig | Neuromics | GP14100 | 1:500, 1:100 | iFix & iFr-F | Batch & sample variability: Staining might present high background and dim signal |
| Neuron subpopulation - Themal sensing | TRPA1* | rabbit | Abnova | PAB11992 | 1:100 | iFix & iFr-F | |
| Neuron subpopulation - Themal sensing | TRPA1 | Rabbit | Novus Biological | NB110-40763SS | 1:200 | iFix | Not tested in iFr-F |
| Neuron subpopulation - Silent nociceptor | CHRNA3* | Rabbit | Alomone | ANC-003 | 1:50 | iFix & iFr-F | |
| Neuron subpopulation - Propioceptors | Parvalbumin* | Sheep | R&D Systems | AF5058 | 1:50 | iFix | **Fixation variability:** Weak staining in iFr-F samples |
| Neuron subpopulation - A fiber subpopulation | TrkC/ NTRK3* | Sheep | Osenses | OST00119W | 1:200, 1:500 | iFix & iFr-F | **Sample variability:** May present high signal from surrounding cells |
| Neuron subpopulation - Cold sensing | TRPM8* | Rabbit | Origene | TA336867 | 1:200 | iFix & iFr-F | **Sample variability**: May stain different ammount of neurons and nuclear staining in some samples |
| Neuron subpopulation - Itch sensation sensory neurons | OSMR* | Rabbit | Abcam | ab232684 | 1:50 | iFix & iFr-F | |
| Neurodegeneration | ATF3* | Rabbit | Santa Cruz | sc-188 | 1:200 | iFix & iFr-F | **Fixation variability:** Additionally to neuronal signal, may present a dim staining of non-neuronal nuclei in iFr-F tissue |
| Synaptic vesicles | Synapsin 1/2 | Chicken | Synaptic Systems | 1006006 | 1:200 | iFix & iFr-F | |
| Neuronal membrane | Neurexin-2 | Rabbit | ThermoFisher | BS-11104R | 1:50 | iFix & iFr-F | Dim signal |

*: Used in manual quantification
