## Supplementary Table 4 for "Tissue-to-Analysis Framework Enables Multiscale Mapping of the Architectural and Cellular Organization in the Human Dorsal Root Ganglion"

| Non-neuronal markers | | | | | | | |
|---|---|---|---|---|---|---|---|
| **Cell population targeted** | **Marker** | **Host** | **Vendor** | **Catalog** | **Dilution** | **Verified method** | **Comments** |
| Blood Vessels - Pan-endothelial | CD31 | Rabbit | Abcam | Ab28364 | 1:50 | iFix & iFr-F | Clean blood vessel signal. |
| Blood Vessels - Pan-endothelial | CD31 | Sheep | R&D Systems | AF806 | 1:50, 1:500, 1: 1000 | iFix & iFr-F | **Fixation variability:** Weak staining in iFix samples. **Sample variability**: May stain macrophages. |
| Blood Vessels - Pan-endothelial | CD31 | Goat | R&D Systems | AF3628 | 1:100, 1:200 | iFix & iFr-F | Some non-specific staining appearing in neurons . |
| Blood Vessels - Veins & capillaries | Aquaporin 1 | Mouse | Santa Cruz | sc-25287 | 1:50 | iFr-F | Not tested in iFix samples. |
| Blood Vessels - Veins & capillaries | Caveolin 1 (D46G3) | Rabbit | Cell Signaling | 3267 | 1:500, 1:800 | iFr-F | Not tested in iFix samples. |
| Blood Vessels - Veins & capillaries | CD34 | Mouse | DAKO | M7165 | 1:50 | iFr-F | Predominant Blood vessel staining, dim fibroblast signal. |
| Blood Vessels - Veins & capillaries | PLVAP | Mouse | Abcam | ab81719 | 1:200 | iFr-F | **Fixation variability:** Weak staining in iFix samples. |
| Blood Vessels - Arteries & capillaries | CLDN5 | Rabbit | Invitrogen | 34-1600 | 1:50 | iFix & iFr-F | High non-specific staining in neurons. **Fixation variability:** Weak staining in iFix samples. |
| Smooth Muscle Cells | ACTA2 (alpha SMA) | Mouse | Santa Cruz | sc-32251 AF647 | 1:500 | iFix & iFr-F | |
| Pericytes | NG2 | Rabbit-Biotin Conjugated | Abcam | AB5320B | 1:100 | iFix & iFr-F | Also highly present in SGCs and other cells |
| Fibroblasts | EGFR | Rabbit | Invitrogen | PA1-1100 | 1:100 | iFix & iFr-F | Some neuronal staining, specially high in neuronal nuclei. |
| Fibroblasts | Glut-1 (SLC2A1) | Rabbit | EMD Milipore | 07-1401 | 1:500 | iFix & iFr-F | |
| Fibroblasts, neurons | P75NTR / NGFR | Rabbit | Abcam | ab52987 | 1:200 | iFix | Not tested in iFr-F samples; it is also a neuronal marker. |
| Fibroblasts & glial cells | Vimentin | Goat | R&D Systems | AF2105 | 1:100 | iFix & iFr-F | Express at variable degree in most non-neuronal cells. |
| Neural crest- Schwann cells & Satellite Glial Cells | S100beta | Rabbit | Abcam | ab52642 | 1:500 | iFix | Also present in some neurons. |
| Neural crest- Schwann cells & Satellite Glial Cells | SOX10 | Goat | R&D Systems | AF2864-SP | 1:200 | iFix & iFr-F | |
| Neural crest- Schwann cells & Satellite Glial Cells | SOX10 | Mouse | Santa Cruz | Sc-365692 | 1:200 | iFix & iFr-F | |
| Myelinating Schwann cells | MBP | Rat | Abcam | AB7349 | 1:200 | iFix & iFr-F | |
| Satellite Glial Cells | Connexin 43 | Rabbit | Sigma | C6219-25UL | 1:200 | iFr-F | Not tested in iFix samples. |
| Satellite Glial Cells | FABP7 | Rabbit | Invitrogen | PA5-24949 | 1:100 | iFix | **Fixation variability:** Loss of specific SGC signal and high neuronal background in iFr-F samples. |
| Satellite Glial Cells | FABP7 | Goat | R&D Systems | AF3166 | 1:200 | iFix | Unspecific fiber and small neuron labeling. **Fixation variability:** Loss of specific SGC signal in iFr-F samples. |
| Satellite Glial Cells | FASN | Rabbit | Abcam | ab128870 | 1:100, 1:500, 1:5000(TSA) | iFix & iFr-F | **Sample variability:**May present high signal/noise ratio (solvable with TSA). |
| Satellite Glial Cells | Kir 4.1 | Rabbit | Milipore | AB5818 | 1:500, 1:1000 | iFr-F | Not tested in iFix samples. |
| Satellite Glial Cells | Kir 4.1 | Rabbit | Alomone | APC-035 | 1:200 | iFr-F | Not tested in iFix samples. |
| Satellite Glial Cells - Subset | GFAP | Rabbit | DAKO | Z0334 | 1:1000 | iFix & iFr-F | Present in a subset of glia cells and other non-neuronal cells. |
| Macrophages - pan-marker | IBA1 | Chicken | Synaptic Systems | 234009 | 1:500 | iFix & iFr-F | **Fixation variability:** May not stain nuclear region in iFr-F samples. |
| Macrophages - pan-marker | IBA1 | Rabbit | Wako | 019-19741 | 1:200 | iFix & iFr-F | **Fixation variability:** May not stain nuclear region in iFr-F samples. |
| Macrophages | CD32 | Goat | R&D Systems | AF1330 | 1:50 | iFix & iFr-F | |
| Macrophages and antigen presenting cells | HLA-Dr | Rabbit | Abcam | ab103998 | 1:50 | iFix & iFr-F | |
| Macrophage - Perivascular macrophages | CD163 | Mouse | Novus Biologicals | NB110-40686 | 1:500 | iFix & iFr-F | |
| Macrophage subset | FOLR2 | Mouse | Abcam | 327002 | 1:500 | iFix & iFr-F | **Fixation variability:** May stain dim in iFix samples |
| Macrophage subset | MRC1 | Mouse | Thermo | MA5-44147 | 1:50 | iFr-F | Not tested in iFix samples. |
| Macrophage subset | VSIG4 | Mouse | BioTechne | MAB46463-SP | 1:50 | iFix & iFr-F | Appears to stain all macrophages. |
| Macrophages - Perineuronal macrophages | CX3CR1 | Rat | Nordic Biosite | 341602 | 1:50 | iFix & iFr-F | **Fixation variability:** Weaker in iFix samples. |
| B-Cells | CD19 | Rabbit | Abcam | ab134114 | 1:50, 1:500 | iFix | Not tested in iFr-F samples. |
| T-Cells | CD3 | Rabbit | DAKO | A0452 | 1:200 | iFix & iFr-F | |
| Monocytes & other immune cells | S100A4 | Mouse | Atlas Antibodies | AMAb90599 | 1:3000 | iFix & iFr-F | Bright staining of monocytes and dimmer staining in some T-cells. |
| Mast Cells | FCeR1alpha | mouse | Invitrogen | 12-5899-42 | 1:50 | iFix & iFr-F | Dim staining, difficult to distinguish from background. |
| Mast Cells | KIT / C-KIT / CD177 | Mouse | BD | 333233 | 1:100 | iFix & iFr-F | Also in FCER1A- cells. |
| Neutrophils | CD15 | Mouse | DAKO | GA062 | 1:1 (Sold as solution ready to stain) | iFix & iFr-F | **Sample variability:** May present different cell numbers. |
| Neutrophils | Neutrophil Elastase | Rabbit | Abcam | AB131260-1001 | 1:50 | iFix & iFr-F | Also present in cells with no trilobular nuclei. Sample variability: May present different cell numbers. |
| Melanocytes | MelanA | Sheep | R&D Systems | AF8008 | 1:200 | iFix & iFr-F | |
| Cell structure - F-Actin | Phalloidin | Amanita phalloides | Abcam | AB176753 | 1:2000 | iFix & iFr-F | |
| Mitochondrial activity | TSPO | Rabbit | Abcam | ab109497 | 1:500 | iFix & iFr-F | |
| Cell Proliferation | KI67 | Rat | Invitrogen | 14-5698-82 | 1:100 | iFix & iFr-F | **Sample variability:** May present different staining levels. |
| Cell senescence | P16 | Mouse | Invitrogen | MA5-17054 | 1:500 | iFix & iFr-F | **Sample variability:** May present different staining levels, mostly glial cells. |

0.16 0.11
