## Supplementary Table 5 for "Tissue-to-Analysis Framework Enables Multiscale Mapping of the Architectural and Cellular Organization in the Human Dorsal Root Ganglion"

| Donor | Age | Sex | Fixation time-point | Cause of Death |
|---|---|---|---|---|
| 1 | 40 | M | iFix & iFr-F | Cerebral hemorrhage |
| 2 | 55 | M | iFix & iFr-F | Cranial trauma / Anoxia |
| 3 | 61 | M | iFix & iFr-F | Suicide (CO Anoxia) |
| 4 | 33 | F | iFix & iFr-F | Suicide (Hanging) |
| 5 | 47 | F | iFix & iFr-F | Vein thrombosis at the transverse sinus |
| 6 | 72 | F | iFix & iFr-F | Cerebral hemorrhage (Car accident) |
| 7 | 35 | M | iFix | Suicide (Hanging, anoxia) |
| 8 | 60 | M | iFix | Trauma (Close head injury) |
| 9 | 45 | F | iFix | Brain hemorrhage |
| 10 | 54 | F | iFix | Stroke |
| 11 | 56 | F | iFix | Brain death (Cerebral venous sinus thrombosis) |
| 12 | 69 | M | iFr-F & iFr-FOS | Brain hemorrhage |
| 13 | 27 | F | iFr-F & iFr-FOS | Suicide (Hanging) |
| 14 | 52 | F | iFr-F & iFr-FOS | Brain hemorrhage |
| 15 | 53 | F | iFr-F & iFr-FOS | Cranial trauma |
| 16 | 38 | F | iFr-F & iFr-FOS | Brain hemorrhage |
| 17 | 47 | M | iFr-FOS | Brain hemorrhage |

iFix: Immediately fixed at tissue collection; iFr-F: Immediately frozen, fixed before cryosectioning; iFr-FOS: Immediately frozen, fixed on slide
